# Whole organism snRNA-seq reveals systemic peripheral changes in Alzheimer’s Disease fly models

**DOI:** 10.1101/2024.03.10.584317

**Authors:** Ye-Jin Park, Tzu-Chiao Lu, Tyler Jackson, Lindsey D. Goodman, Lindsey Ran, Jiaye Chen, Chung-Yi Liang, Erin Harrison, Christina Ko, Ao-Lin Hsu, Shinya Yamamoto, Yanyan Qi, Hugo J. Bellen, Hongjie Li

## Abstract

Peripheral tissues become disrupted in Alzheimer’s Disease (AD). However, a comprehensive understanding of how the expression of AD-associated toxic proteins, Aβ42 and Tau, in neurons impacts the periphery is lacking. Using *Drosophila*, a prime model organism for studying aging and neurodegeneration, we generated the Alzheimer’s Disease Fly Cell Atlas (AD-FCA): whole-organism single-nucleus transcriptomes of 219 cell types from adult flies neuronally expressing human Aβ42 or Tau. In-depth analyses and functional data reveal impacts on peripheral sensory neurons by Aβ42 and on various non-neuronal peripheral tissues by Tau, including the gut, fat body, and reproductive system. This novel AD atlas provides valuable insights into potential biomarkers and the intricate interplay between the nervous system and peripheral tissues in response to AD-associated proteins.

## Introduction

Alzheimer’s Disease (AD) has been associated with disruptions in the gut microbiota, cardiovascular function, and hormone homeostasis (Menengiç et al., 2020; Stampfer, 2006; Vogt et al., 2017; Xiong et al., 2022). Moreover, peripheral changes, such as inflammation and immune decline, contribute to AD pathogenesis, highlighting the involvement of peripheral tissues as potential contributors to disease (Bettcher et al., 2021). Understanding AD-induced peripheral changes is important for identifying novel biomarkers and gaining insights into how peripheral alterations contribute to AD. Amyloid plaques and hyperphosphorylated Tau tangles are the key hallmark pathologies used to define AD and most studies focus on the impact of expressing these AD-associated proteins in neurons (Scheltens et al., 2016). Neuronal changes will also impact other tissues outside of the brain. Hence, a comprehensive comparison across all cell types throughout the entire body is needed to provide a further understanding of human disease.

The fruit fly, *Drosophila melanogaster,* is a prime experimental model system for studying aging and age-related diseases, including AD. Many key aging pathways are conserved from flies to mammals and have been primarily characterized using fly models. This includes the role of insulin signaling, oxidative stress, dietary restriction, and circadian rhythm in aging and disease (Piper and Partridge, 2018; Tatar et al., 2014). Further, *Drosophila* is a valuable organism for identifying non-cell autonomous mechanisms relevant to human disease. For example, research initiated in flies revealed an uncharacterized mechanism by which glia protect neurons from reactive oxygen species (Liu et al., 2015). This pathway involves the formation of glial lipid droplets which contain peroxidized lipids sourced from neurons (Ioannou et al., 2019; Liu et al., 2015). Importantly, research is accumulating using flies and rodents, with supportive human data that this pathway is disrupted in aging and in AD (Claes et al., 2021; Liu et al., 2017; Marschallinger et al., 2020; Moulton et al., 2021). Overall, this highly conserved pathway was identified and characterized successfully in vivo using *Drosophila* due to the breadth of genetic tools available in this organism.

Advances in single-cell RNA sequencing (scRNA-seq) have promoted its use in defining new cell-types and progressive changes in individual cell types relevant to the aging process (Almanzar et al., 2020; Davie et al., 2018; Li et al., 2022; Lu et al., 2023; Roux et al., 2023; Scheltens et al., 2016; Siletti et al., 2023; Sziraki et al., 2023; Zhang et al., 2020). *Drosophila* offers a unique advantage in scRNA-seq research as it allows a system-level, single-cell survey of progressive transcriptional changes in the whole organism. Our single-nucleus RNA sequencing (snRNA-seq) platform has enabled the compilation of the Fly Cell Atlas (FCA; the first system-level transcriptomic dataset at single-cell resolution of adult *Drosophila* (Li et al., 2022)) and the Aging Fly Cell Atlas (AFCA; a comprehensive single-cell transcriptomic dataset interrogating aging features in *Drosophila* (Lu et al., 2023)). Building on this experimental and bioinformatics platform, we can now achieve a comprehensive characterization of all peripheral tissues across the whole organism in response to expression of genes associated with AD.

Here, we present the Alzheimer’s Disease Fly Cell Atlas (AD-FCA), which profiles single-nucleus transcriptomes of two established AD fly models during the aging process using the whole organism. These established AD models express either a secreted human amyloid-β 42 peptide (Aβ42) or wild-type human Tau (hTau), specifically in neurons of the fly brain, and cause neurodegenerative effects such as reduced lifespans, motor defects, and neuronal cell loss (Chouhan et al., 2016; Iijima et al., 2004; Prüßing et al., 2013; Scheltens et al., 2021; Wittmann et al., 2001). We include both fly AD models to explore the potential differential impacts of expressing Aβ42 versus hTau in neurons. Hereafter, we refer to them as Aβ42 flies and hTau flies. Our data revealed that Aβ42, but not hTau, primarily impacts neurons. Specifically, our snRNA-seq data and functional studies revealed that sensory neurons, including olfactory receptor neurons and auditory neurons, are among the most vulnerable cell types in Aβ42 flies. In contrast to Aβ42, our snRNA-seq and functional data revealed that neuronal expression of hTau impacts a number of non-neuronal peripheral cell types, including gut, fat body, and reproductive system. We highlight defects in cell-cell interactions that may be driving these effects. Interestingly, many of the peripheral changes in the hTau fly model mimic aging phenotypes, suggesting that neuronal expression of hTau causes accelerated aging in peripheral tissues.

Overall, this whole organism AD atlas provides a valuable resource for the neurodegeneration and AD research community. The AD-FCA enables us to uncover many progressive changes that likely occur in non-neurological tissues in response to expression of AD genes, offering fresh insights into the realm of brain-body communication. All data and annotations can be accessed through the visualization and analysis portal at https://hongjielilab.org/adfca/ (figs. S3C, and S6-7).

## Result

### Single-nucleus transcriptomes of the entire fly from two AD models

To generate the AD-FCA, we first characterized the lifespan and motor activity, using the negative geotaxis assay, of flies expressing Aβ42 or hTau. To avoid effects of expressing these proteins through development, we used a temperature-controlled pan-neuronal GAL4 system (*tub-GAL80^ts^; nSyb-GAL4*) that restricts the expression of Aβ42 (*UAS-Aβ42*) or hTau (*UAS-hTau^0N4R^*) to neurons at the adult stage (Fig. 1A). To minimize the genetic background effect, we expressed *UAS-empty* in control flies. Aβ42 or hTau expression in the brain was confirmed by immunofluorescence staining using anti-Aβ42 or anti-phospho-Tau (AT8) antibodies (fig. S1). The lifespan and motor activity experiments showed that 10 days and 20 days of Aβ42-expression represented early and late time-points for disease progression and flies from these time-points were selected for transcriptomic profiling (Fig. 1B, C, and fig. S2). For hTau flies, 20 days and 30 days represented early and late time-points for disease progression and were selected for transcriptomic profiling. Control flies were collected in parallel at all three ages (10-, 20- and 30-day). We performed droplet-based single-nucleus RNA-seq (snRNA-seq) on whole head tissue or whole body (excluding the head) tissue in males and females separately (Fig. 1D, E). We followed the same snRNA-seq pipeline that was used in the Fly Cell Atlas and Aging Fly Cell Atlas except for the modification that high-throughput kits were used in this study to improve the yield of nuclei as well as the number of genes detected (fig. S3A, B). In total, we obtained 624,458 high-quality nuclei covering 18 broad cell classes (Fig. 1F).

**Fig. 1.**
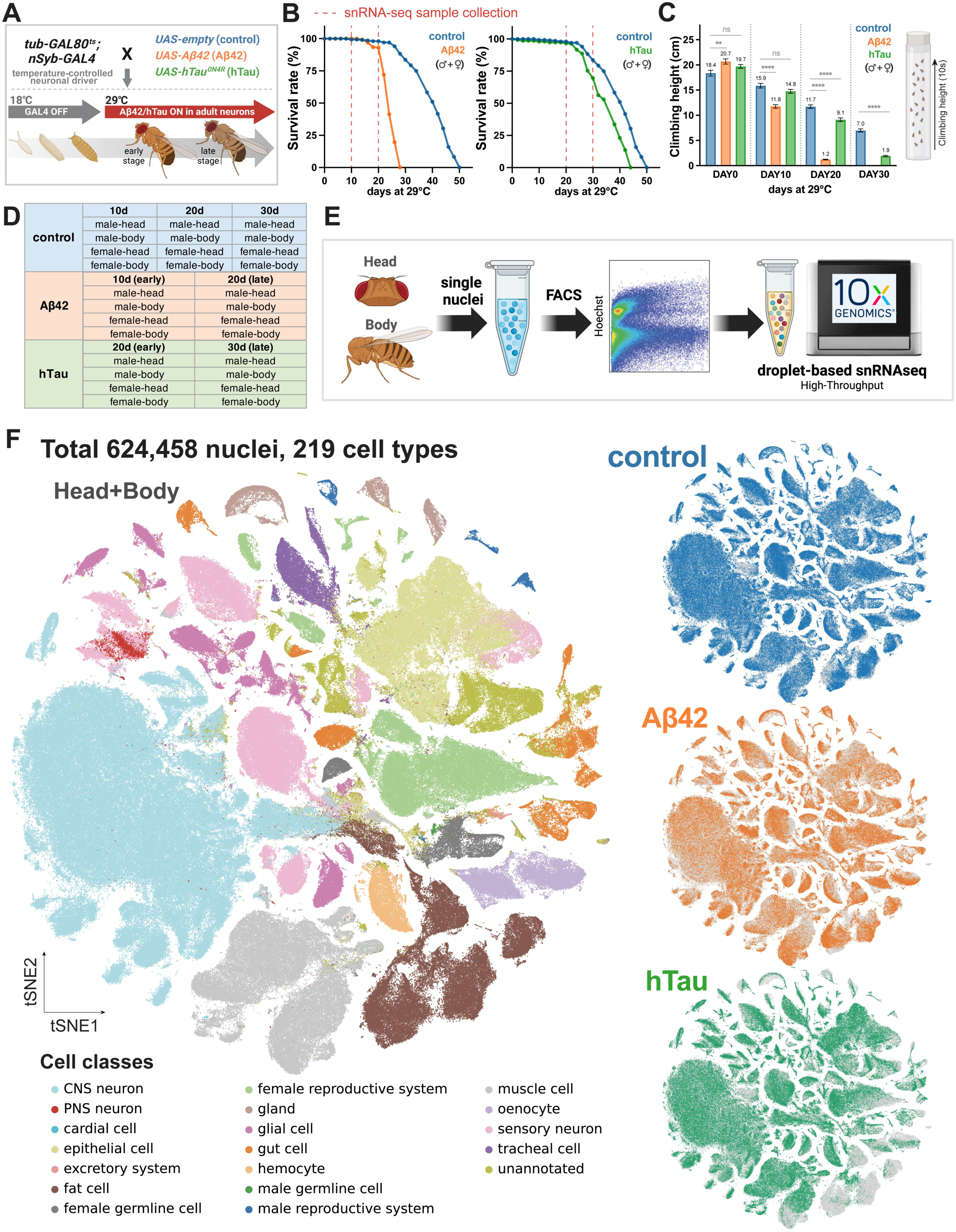
Overview of the AD-FCA. (A) AD models of AD-FCA. Temperature-controlled pan-neuronal driver flies crossed to *UAS-empty*, *UAS-Aβ42*, or *UAS-hTau* transgenic flies. To avoid effects of transgene expression during the developmental process, flies were kept at 18°C for cross and development. Progenies were collected after eclosion, and moved to 29°C to express the transgenes in pan-neurons during the adult stage. (B) Survival curves at 29°C (female and male combined). Female and male flies were counted separately. N=300 (150 female and 150 male) per genotype. See separate lifespan curves in fig. S2. (C) Aβ42 (10d and 20d) and hTau (20d and 30d) flies have climbing defects compared to control in a negative geotaxis assay. 20 flies per genotype per set. Flies climbed 10 times per set for 10 seconds. 5 sets of 0d (N=100, 40 female and 60 male per genotype), 8 sets of 10d (N=160, 80 female and 80 male per genotype), 10 sets of 20d (N=200, 100 female and 100 male per genotype), 10 sets of 30d (N=200, 100 female and 100 male per genotype). Bars represent mean ± SEM. ** p ≤ 0.01,**** p < 0.0001. P-values were computed using the parametric unpaired t-test. The numbers above the bars indicate mean values. Female and male flies were counted separately. See fig. S2. (D) The heads and bodies of females and males from each genotype were collected separately (50 heads or 15 bodies per sample). Aβ42 flies were collected at 10d and 20d, and hTau flies were collected at 20d and 30d. Control flies were collected at 10d, 20d, and 30d. (E) Flowchart of the snRNA-seq experiment. FACS (fluorescence-activated cell sorting) was used to sort nuclei. 10X Genomics Chromium Next GEM Single Cell 3’ HT (high-throughput) kits were used for library generation to capture over 20,000 nuclei per sample. (F) t-distributed Stochastic Neighbor Embedding (tSNE) visualizations showing 18 broad cell classes of datasets (head + body) integrated across different samples (left). tSNE visualizations separated by genotype (right).

Next, we employed a supervised machine learning-based method for annotating the cell types based on reference datasets from the Fly Cell Atlas, the Aging Fly Cell Atlas, and a comprehensive single-cell dataset from the fly optical lobe (Li et al., 2022; Lu et al., 2023; Özel et al., 2021). We manually validated each annotation using cell type-specific markers. We were able to annotate most cell clusters from the body, but many neurons from the head remained unannotated. This is largely due to the high complexity of the fly central brain and the fact that many types of central nervous system (CNS) neurons only represent a very small number of cells. We annotated these cells as uncharacterized CNS neurons. Overall, we identified 219 different cell types, including 144 cell types from the head and 75 cell types from the body displaying marked improvement from the Aging Fly Cell Atlas annotation (total 163 cell types) (Fig. 1F, and figs. S4-5).

### Sensory neurons are among the top vulnerable cell types in the head of Aβ42 flies

We next developed a comprehensive comparison among all different cell types, including CNS neurons, peripheral sensory neurons, glia, and other non-neural cells across the whole head in these two AD models.

First, we assessed whether and how Aβ42 or hTau expression impacts cellular composition across the whole head (Fig. 2A). Note that we based our measurements on nuclear composition as we performed single-nucleus sequencing. Because there is very limited neurogenesis or gliosis in adult fly brains (Li and Hidalgo, 2020), we expect that most cellular composition changes will be caused by cell loss. The snRNA-seq data revealed that Aβ42, but not hTau, caused a decrease of the nuclear ratio in many cell types including different CNS neurons and sensory neurons (fig. S9A). Consistently, we found that Aβ42 fly brains show more dramatic vacuolization than hTau flies (fig. S8). This is also consistent with a previous study (Chouhan et al., 2016). Among the top ten most affected cell types in Aβ42 flies are 7 CNS neurons, 2 sensory neurons, and 1 glial cell type (Fig. 2B, C). Interestingly, these cell types are directly associated with sensory functions, including vision, audition, and olfaction (Fig. 2C), suggesting that the sensory system is vulnerable to Aβ42 expression. To validate these findings, we stained for neuronal nuclei in the second segment of the antenna where the neuronal cell bodies of the auditory sensory neurons are located. We observed a significant reduction in the number of auditory sensory neurons in Aβ42 flies compared with control flies (Fig. 2D). The impact on these sensory neurons was specific to Aβ42 expression as it was not detected in control flies. It is not due to differential expression of the *GAL4/UAS* system used to express Aβ42 as we found that there is a negligible correlation coefficient (r=-0.09, p=0.68) between *nSyb* expression levels and the nuclear ratio changes in neurons (fig. S9C). Hence, sensory neurons are among the top cell types that are susceptible to Aβ42-associated toxicity.

**Fig. 2.**
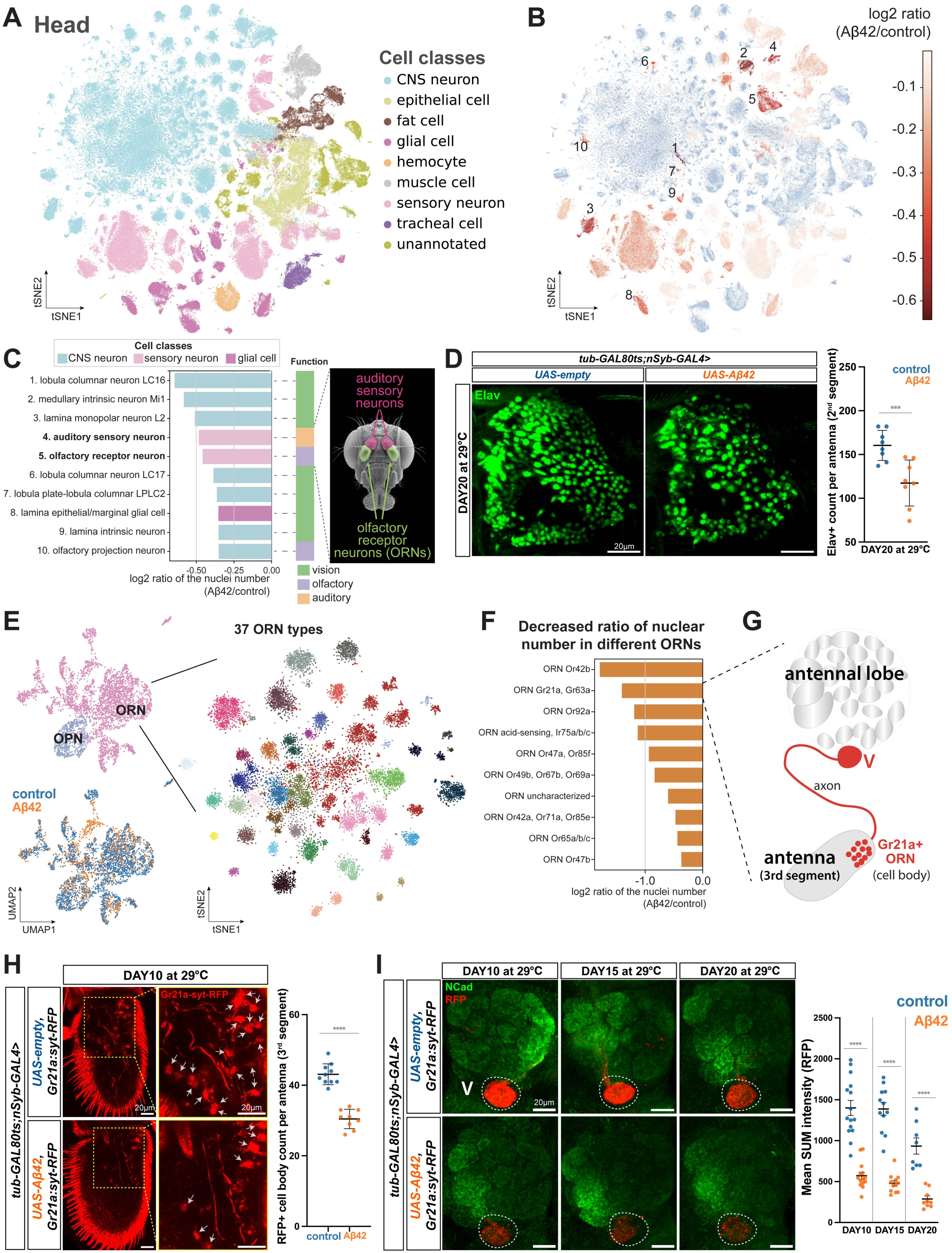
Cell vulnerability in the head of Aβ42 flies. (A) Broad cell classes of different head cell types. (B) Decrease of nuclear ratios in Aβ42 flies. Nuclear ratios are compared between Aβ42 and control flies and the top 10 cell types with the biggest decrease are marked with the ranking on the tSNE plot. Numbers associated with the cell types are shown in Fig. 2C. (C) The top 10 cell types with the lowest nuclear ratios in Aβ42 flies. Each cell type is colored by the corresponding broad cell classes and associated with its biological function in the adjacent color bar. The cartoon illustrates the locations of two types of sensory neurons. (D) Auditory neurons located in the 2nd segment of the antenna are shown by anti-Elav staining. Elav+ nuclei were counted from whole Z series images. N=8 per genotype. Scatter plot with mean ± SD. *** p < 0.001. The representative images were generated by MAX intensity of all Z planes. (E) Olfactory receptor neuron (ORN) and olfactory projection neuron (OPN) clusters are shown with corresponding cell types and genotypes (left). ORNs are further sub-clustered into 37 detailed ORN types (right). See fig. S10 for annotation. (F) ORN subtypes dislaying the largest decrease in nuclear ratios. (G) Schematic of *Gr21a+* ORN cell bodies and axon projection (red). (H) The number of *Gr21a+* ORN cell bodies in the 3rd segment of antenna is significantly reduced in Aβ42 flies compared to control at 10d (yellow dotted box). The RFP+ cell bodies (red) were counted from whole Z series images. Control N=10, Aβ42 N=9. Scatter plot with mean ± SD. ** p ≤ 0.01. The representative images were generated by MAX intensity of Z planes containing these cell bodies. The off-target signal detected outside of the yellow dotted box region is due to the laser reflection on the cuticle. (I) The axon terminal RFP signals of *Gr21a+* ORNs are significantly decreased compared to age-matched control flies. anti-RFP (red, Gr21a+ ORN axon terminal) and anti-NCad (green, neuropil of the antennal lobe). The mean RFP signal intensity/area in V region was quantified from SUM intensity images by averaging measurements from both sides of the antennal lobes. N=8-15 per genotype/age. Scatter plot with mean ± SEM. ** p ≤ 0.01. The representative images were generated by MAX intensity of all Z planes. P-values in D and I were computed using the parametric unpaired t-test.

Second, we performed differentially expressed gene (DEG) analysis on each cell type to compare Aβ42 or hTau flies to corresponding controls (fig. S9B). Three sensory neuronal types, including outer photoreceptors (R1-R6), auditory sensory neurons, and olfactory receptor neurons (ORNs), are among the top ten cell types with the highest number of DEGs in Aβ42 flies. We also observed that Aβ42 or hTau expression caused very different DEG patterns that were dependent on the cell type. For example, in auditory sensory neurons, Aβ42 showed about 400 DEGs while hTau showed only a few DEGs. Note that some non-neuronal cells, such as head fat body cells, also showed high DEGs, suggesting that neuronal exposure to Aβ42 and hTau may precipitate non-cell-autonomous effects on these cells (see below).

These analyses indicate that the fly sensory systems, including sensory neurons which directly encounter the environmental stimuli and send information to the brain, are among the most susceptible cell types to Aβ42 expression. This is consistent with several human AD studies which showed that olfactory decline is one of the earliest symptoms in AD patients (Pacyna et al., 2023; Tian et al., 2022).

### Different subtypes of ORNs show different vulnerability to Aβ42

The decline in olfactory sensitivity is prevalent in different types of dementia: nearly 100% in Alzheimer’s disease (AD), 90% in Parkinson’s disease, and 15% in vascular dementias (Doty, 2012; Duff et al., 2002). However, olfactory dysfunction is frequently overlooked by both physicians and patients, as well as in AD model studies, due to three major challenges. First, aging can also cause olfactory decline and it is difficult to distinguish age-versus AD-caused olfactory decline. Second, there are many types of ORNs (about 800 in humans) that detect different odors and it is difficult to assess which ORNs are specifically affected by AD. Third, the olfactory neural circuit contains ORNs and downstream neurons that relay the olfactory information to the higher brain center, and it is largely unknown which neurons are primarily affected.

Our current data provide a novel opportunity to address these questions. Since we sequenced age-matched controls with the Aβ42 flies, we can distinguish the effects caused by Aβ42 expression versus aging. In flies, there are about 50 pairs of olfactory neurons, including primary ORNs and secondary olfactory project neurons (OPNs), providing a simplified system to examine neuronal specificity. Our snRNA-seq data captured both ORNs and OPNs, allowing for direct comparisons. Interestingly, we found that ORNs are more affected than OPNs based on both cell loss analysis (ranked as #5 for ORNs and #10 for OPNs from the top ten cell types; Fig. 2C) and DEG analysis (fig. S9B). Further, compared with age-matched controls, Aβ42 ORNs showed more cell loss (Fig. 2C), suggesting that ORNs are primarily impacted by Aβ42-associated toxicity versus aging. Next, we sub-clustered all ORNs, and they formed 37 distinct clusters, each with the expression of specific olfactory receptors (Fig. 2E and fig. S10). At this resolution, we were able to rank these ORN types based on cell loss and observed that *Or42b* and *Gr21a/Gr63a* ORNs are the top two affected types (Fig. 2F). To further validate the sensitivity of ORNs to Aβ42, we expressed the ORN reporter, *Gr21a:syt-RFP*, in Aβ42 and control flies and found that both cell body numbers and axon terminal signals of syt-RFP labeled ORNs are significantly decreased in response to Aβ42 (Fig. 2G-I); this reporter expresses an RFP-tagged synaptotagmin protein under control of the *Gr21a* promoter (Jones et al., 2007). Hence, sensory neuron loss is primarily driven by Aβ42-associated toxicity, not by aging.

### Aβ42-specific neuron cluster and endoplasmic reticulum (ER) stress response

Next, we sought to identify possible mechanisms for Aβ42-induced cell loss of different neuronal types. We observed an Aβ42-specific cluster which expresses neuronal markers and separates from other neurons identified in hTau and control flies (Fig. 3A, and fig. S11A, B). There are two possibilities for the origin of this cluster: 1) these cells arise from a certain neuronal type whose transcriptome is uniquely changed by Aβ42 expression, or 2) these cells represent many different types of neurons that converge to have a similar transcriptional state due to Aβ42 expression.

**Fig. 3.**
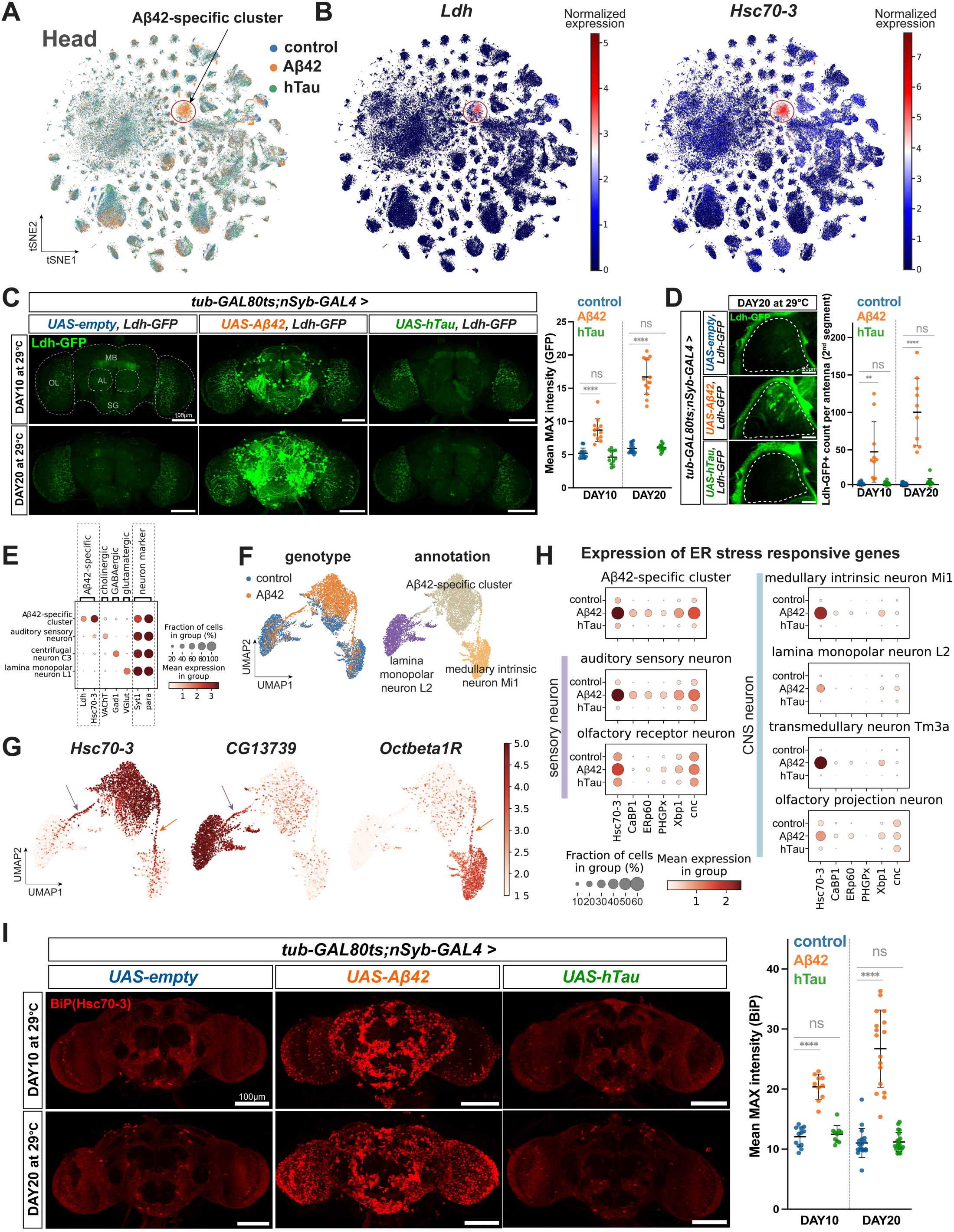
Characterization of the Aβ42-specific cluster. (A) The tSNE plot is colored by different genotypes and the Aβ42-specific cluster is circled and highlighted. (B) The expression of two genes enriched in the Aβ42-specific cluster. (C, D) Ldh-GFP (green) is highly expressed in brains (C) and auditory neurons (D) of Aβ42 flies, but not in control or hTau flies. (C) The mean GFP signal intensity/area in the whole brain was quantified from MAX intensity images. N=12-15 per genotype/age. Scatter plot with mean ± SD. **** p ≤ 0.0001. The representative images were generated by MAX intensity of all Z planes. (D) The GFP+ cell bodies were counted from whole Z series images. Control N=10-12 per genotype/age. Scatter plot with mean ± SD. ** p ≤ 0.01, **** p ≤ 0.0001. The representative images were generated by MAX intensity of Z planes containing these cell bodies. The off-target signal detected outside of the circled region is due to the laser reflection on the cuticle. (E) Dot plot showing marker genes that are specific to the Aβ42-specific cluster and three types of neurons, as well as two pan-neuronal marker genes. (F) Co-clustering of the Aβ42-specific cluster and two other neuronal types that are vulnerable to Aβ42 expression. The Uniform Manifold Approximation and Projection (UMAP) plots are colored by corresponding genotypes and cell types. (G) UMAPs showing the expression of marker genes specific to the Aβ42-specific cluster, lamina monopolar neuron L2, and medullary intrinsic neuron Mi1. “Bridge” cells connecting different clusters are indicated by arrows. (H) Expression of ER stress markers in cell types that are vulnerable to Aβ42 expression. (I) The Hsc70-3/BiP expression shown by antibody staining in the whole brain is increased in Aβ42 flies, but not in Tau flies, compared with control flies. The mean BiP signal intensity/area in the whole brain was quantified from MAX intensity images. N=10-20 per genotype/age. Scatter plot with mean ± SD. **** p ≤ 0.0001. The representative images were generated by MAX intensity of all Z planes. P-values in C, D, and I were computed using the parametric unpaired t-test.

To test these possibilities, we first identified genes that are specifically expressed in the Aβ42-specific cluster, finding that *lactate dehydrogenase (Ldh)* and *Hsc70-3* are significantly upregulated in these neurons (Fig. 3B, and fig. S11C). To validate that *Ldh* does become upregulated in response to Aβ42, we expressed a *Ldh-GFP* reporter (Bawa et al., 2020) in flies that pan-neuronally expressed *UAS-empty, UAS-Aβ42,* or *UAS-hTau*. This reporter expresses a GFP-tagged Ldh protein under control of the *Ldh* promoter. Indeed, Aβ42 flies exhibit a significant increase in Ldh-GFP levels compared to control or hTau flies (Fig. 3C, D, and fig. S12). Interestingly, Ldh-GFP positive cells are identifiable across different brain regions and in the antenna of Aβ42 flies, demonstrating that they are not localized to a specific neuron population. Consistent with this cluster including different types of neurons, in our snRNA-seq data we found that the neurons within the Aβ42-specific cluster express *VaChT* (marks cholinergic neurons), *Gad1* (marks GABAergic neurons), and *VGlut* (marks glutamatergic neurons), all at low levels (Fig. 3E). To further confirm this, we performed sub-clustering analysis for the Aβ42-specific cluster with two distinct neuronal types, lamina monopolar neuron L2 (L2) and medullary intrinsic neuron Mi1 (Mi1). These two cell types are the top impacted cell types in Aβ42 flies (see Fig. 2C). Intriguingly, both L2 and Mi2 clusters form a bridge-like pattern connecting them with the Aβ42-specific cluster (Fig. 3F). When we checked the expression of genetic markers for the Aβ42-specific cluster (*Hsc70-3*), L2 neuron (*CG13739*), or Mi1 neuron (*Octbeta1R*), we found that these genes are all expressed in the connecting “bridge” cells (Fig. 3G). Altogether, our data suggest that cells in the Aβ42-specific cluster originate from many different types of neurons.

To better understand why this Aβ42-specific cluster has a unique genetic profile, we performed gene-ontology enrichment (GO) analysis on DEGs from these cells (fig. S11D). The top GO terms identified are all related to neuronal physiology and function with one exception, *response to endoplasmic reticulum (ER) stress*. Representative genes in this GO term include *Hsc70-3, CaBP1, ERp60, PHGPx, Xbp1,* and *cnc*, all displaying higher expression levels in the top affected cell types from Aβ42 flies when compared with control or hTau flies (Fig 3H, and fig. S11E). We confirmed that Hsc70-3 protein levels are increased in Aβ42 flies, but not in hTau flies, compared to control flies, by performing immunofluorescence in brains using an anti-Hsc70-3 (BiP) antibody (Fig. 3I). Consistently, these Hsc70-3+ cells in Aβ42 flies are also localized across different brain regions. As the *UAS-Aβ42* construct carries a secretion signal peptide (ssp), which directs the Aβ42 peptides to the ER during the process of protein synthesis and secretion, we considered if the ER stress response seen in Aβ42 flies was a general effect caused by high expression of a secreted peptide. Using a newly generated *UAS-ssp.HA.tag* transgenic fly line, we did not observe increased Hsc70-3 levels when expressed pan-neuronally (fig. S13A). Hence, the ER stress response observed in our Aβ42 flies is specific to the expression of this AD-associated peptide.

It has been shown that persistent activation of ER stress can lead to cell death in various cell types, while its role in the initiation and progression of AD is just beginning to be characterized (Sano and Reed, 2013). We observe that an apoptosis-related gene, *Calr* (Osman et al., 2017), is highly enriched in the Aβ42-specific cluster (fig. S13B). Consistent with this observation, immunofluorescence data showed that Aβ42 brains have higher levels of the apoptosis marker, cleaved Caspase-3, compared to control and hTau brains (fig. S14A). In addition, many Hsc70-3/BiP+ cells in the Aβ42 brain are positive for cleaved Caspase-3(fig. S14B), demonstrating that the ER stressed neurons are undergoing apoptosis. Overall, these data indicate that Aβ42-induced ER stress may cause neuronal loss.

### Neuronal expression of hTau induces various changes in peripheral tissues

To characterize how neuronal expression of AD-associated proteins impact non-neuronal peripheral tissues, we next focused on the snRNA-seq data derived from the fly bodies. (Fig. 4A, B). We found that hTau flies show more obvious changes than Aβ42 flies when compared to control flies for several major tissues including the fat body, oenocyte, gut, and reproductive system. The observed changes are defined based on the overall cell distribution shift in the clustering plot which reflects their transcriptomic status (Fig. 4A). To quantify those changes, we measured differences in nuclear ratio and the number of DEGs between hTau and control flies (Fig. 4C, fig. S15A, B). Indeed, cell types within these tissues displayed divergent transcriptomes and cell compositions, corroborating the distinct clustering shown in Fig. 4A. Functionally, these top impacted peripheral cell types fall into three categories: fat metabolism, digestion, and reproduction. To validate these changes and understand them in detail, we performed further analyses and conducted complementary in vivo assays.

**Fig. 4.**
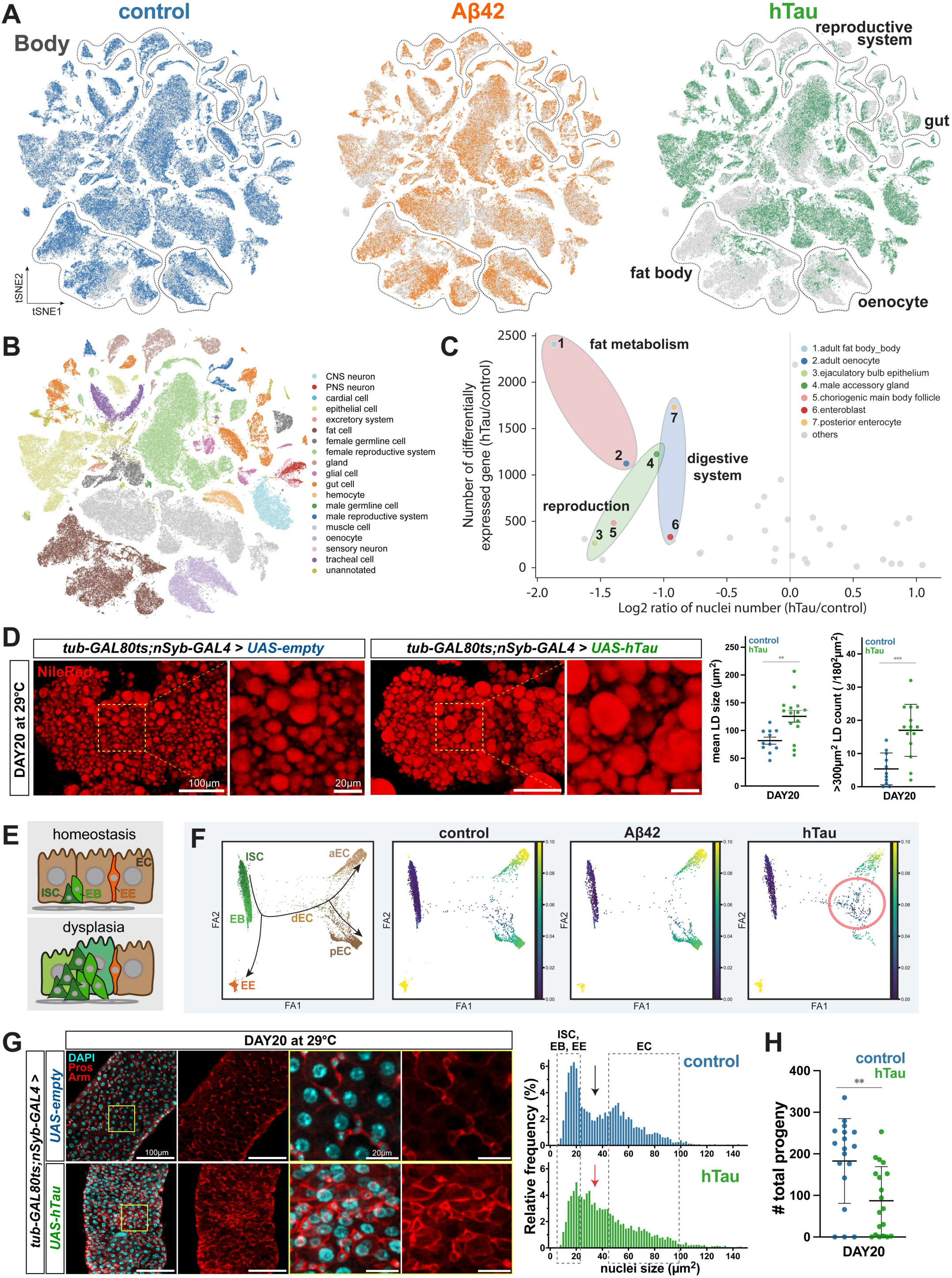
Systemic effects of neuronal hTau expression on peripheral tissues. (A) tSNE plots of body samples colored by genotypes. Clusters showing obvious changes in the hTau flies are indicated. (B) tSNE plot of broad cell classes in body samples. (C) Cell types with significant changes between hTau and control flies regarding DEG number and nuclear ratios. (D) Nile Red staining to visualize lipid droplets (LDs) in the fat body. The 20d hTau flies exhibit significantly enlarged LDs compared to control. The size of each LD was measured from the same area of each sample by StarDist2D and ImageJ. Control N=11, hTau N=15. Scatter plot with mean ± SD. ** p ≤ 0.01, *** p ≤ 0.001. The representative images were generated by MAX intensity of all Z planes. (E) Illustration of the dysplasia phenotype in the fly gut. ISC, intestinal stem cell; EB, enteroblast; EE, enteroendocrine cell; EC, enterocyte. (F) Pseudotime analysis of fly gut cell types from different genotypes. Outlined on the right plot is a group of cells between EBs and mature ECs. aEC, anterior enterocyte; pEC, posterior enterocyte; dEC, differentiating EC. (G) 20d hTau fly guts show a decrease of mature polyploidy enterocytes (large nuclei) and an increase of cells with intermediate-size nuclei with strong armadillo staining. The right two plots show nuclear size distribution. ISCs, EBs, and EEs are diploid cells with small nuclei and mature ECs are polyploidy cells with big nuclei. DAPI (cyan), DAPI (cyan), anti-Prospero (red signals in nuclei, EEs) & anti-armadillo (red signals in cell membrane, stronger in ISCs and EBs than ECs). The size of each nucleus (cyan) was measured from the same area of each sample by StarDist2D and ImageJ. Control N=12, hTau N=16. (H) The 20d hTau male flies show reduced fecundity compared to control. Control N=18, hTau N=19. Scatter plot with mean ± SD. ** p ≤ 0.01. P-values were computed using the parametric unpaired t-test.

To further probe for potential defects in the fat body, we dissected this tissue from hTau or control flies. The fat body is the fly adipose tissue and central storage depot of nutrients and energy reserves. Hence, it can be stained using the neutral lipid dye, Nile red, to visualize lipid droplets (LDs). In 20-day hTau flies, we observed significantly enlarged LDs compared to LDs in age-matched control flies (Fig. 4D and fig. S16). Metabolic pathway analysis on DEGs identified in the fat body from hTau flies also showed that lipid degradation pathway is perturbed in these animals (fig. S17A). For example, several genes involved in the fatty acid degradation pathway are significantly down-regulated, such as *whd (withered)* and *Mcad (Medium-chain acyl-CoA dehydrogenase)* (fig. S17C). Collectively, we found a profound alteration in lipid metabolism that occurs within the fat body in response to neuronal hTau expression.

We next functionally investigated hTau-associated changes in the gut identified in our snRNA-seq analysis. The *Drosophila* gut epithelium is maintained and regenerated by intestinal stem cells (ISCs) through proper proliferation and differentiation in young flies (Micchelli and Perrimon, 2006; Ohlstein and Spradling, 2006). Aging and different stresses, such as oxidative stress induced by oxidants or inflammatory stress triggered by pathogenic gut infection, can cause dysplasia phenotypes (Fig. 4E) (Li and Jasper, 2016). In these conditions, ISCs become hyperproliferative, and their daughter cells, enteroblasts (EBs), cannot properly differentiate into mature enterocytes (ECs) (Li and Jasper, 2016). In a pseudotime analysis, we observed that the ratio of mature enterocytes is reduced and the ratio of transitioning between EBs and mature ECs is increased (Fig. 4F), which indicates there is a dysplasia phenotype. In dissected gut tissue from hTau or control flies, we further found two gut dysplasia-related phenotypes in hTau flies: 1) a decrease of mature polyploidy enterocytes (characterized by their large nuclei) and 2) an increase of cells with intermediate-size nuclei that retain strong anti-armadillo staining signals (Fig. 4G and fig. S18). The strong anti-armadillo signal is normally seen in ISCs and EBs of homeostatic guts.

Note that pan-neuronal drivers, including *elav-GAL4* and *nSyb-GAL4*, are expressed at low levels in EEs (Holsopple et al., 2022a, 2022b; Katheder et al., 2023), consistent with our findings that *nSyb* transcripts are at lower levels in EEs than neurons (fig. S19A). This could impact the identified gut dysplasia phenotypes in hTau flies. It is interesting that hTau expression using *nSyb-GAL4* causes more DEGs in non-EE gut cells (fig. S19B), suggesting that the observed changes may not be solely EE dependent. Furthermore, we document widespread impacts of hTau on many cell types outside the digestive system (see Fig. 5, and Fig. 6). These data support the overall peripheral changes are primarily driven by neuronal hTau expression.

**Fig. 5.**
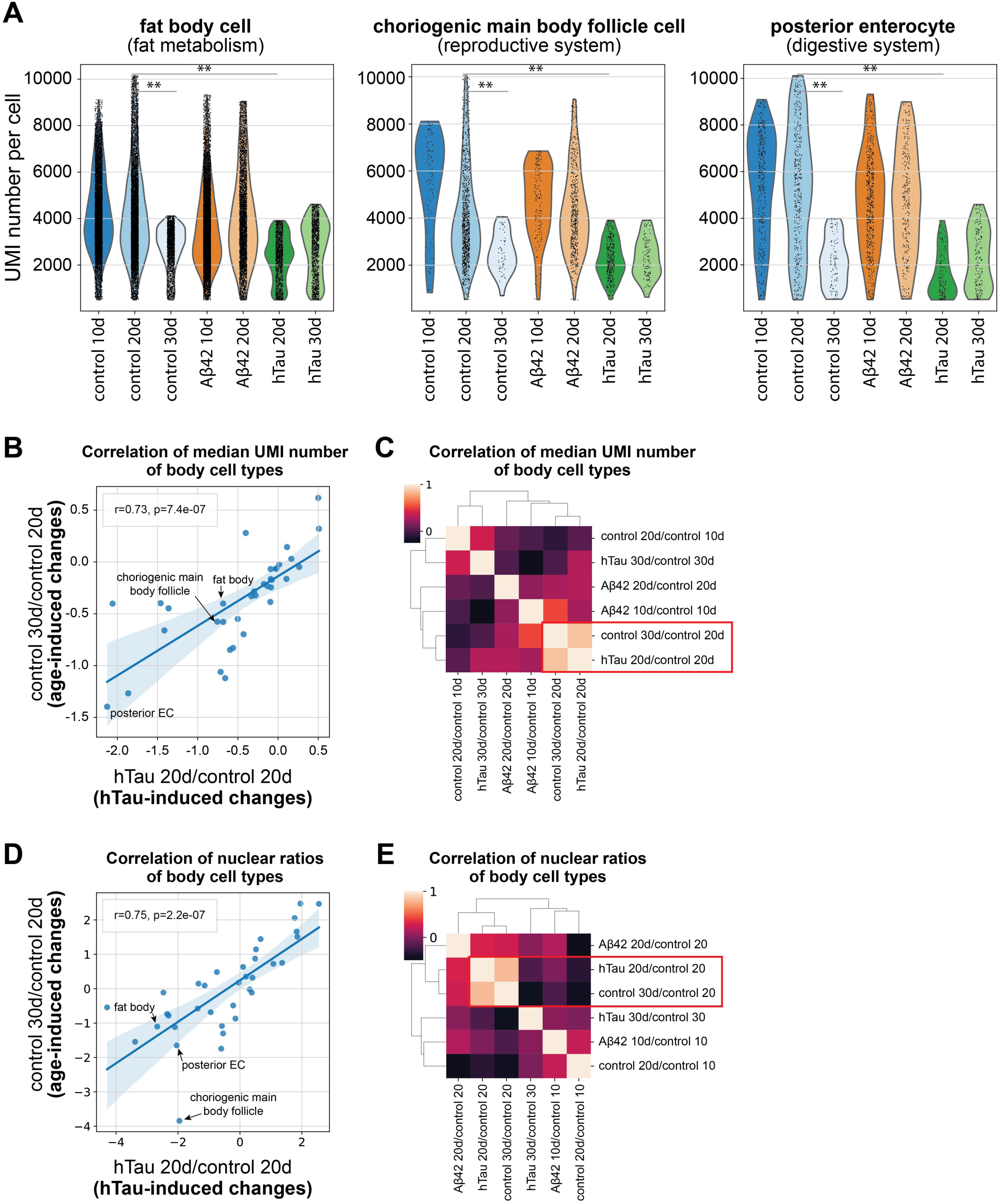
Accelerated aging in hTau flies. (A) Comparison of the number of transcripts per cell (measured by the UMIs) across various ages and genotypes. **: Kolmogorov–Smirnov test, False Discovery Rate < 1e-35. UMI: Unique Molecular Identifier. (B) Pearson correlation of median UMI numbers between age-related changes (30d-control flies versus 20d-control flies) and hTu-induced changes (20d-hTau flies versus 20d-control flies) across different cell types. (C) Spearman correlation of median UMI comparisons between different genotypes and ages. Two comparisons showing the highest correlation are indicated. (D) Pearson correlation of nuclear ratios between age-related changes (30d-control flies at 30d versus 20d-control flies) and hTu-induced changes (20d-hTau flies at 20d versus 20d-control flies)at 20d across different cell types. (E) Spearman correlation of nuclear ratios between different genotypes and ages. Two comparisons showing the highest correlation are indicated.

**Fig. 6.**
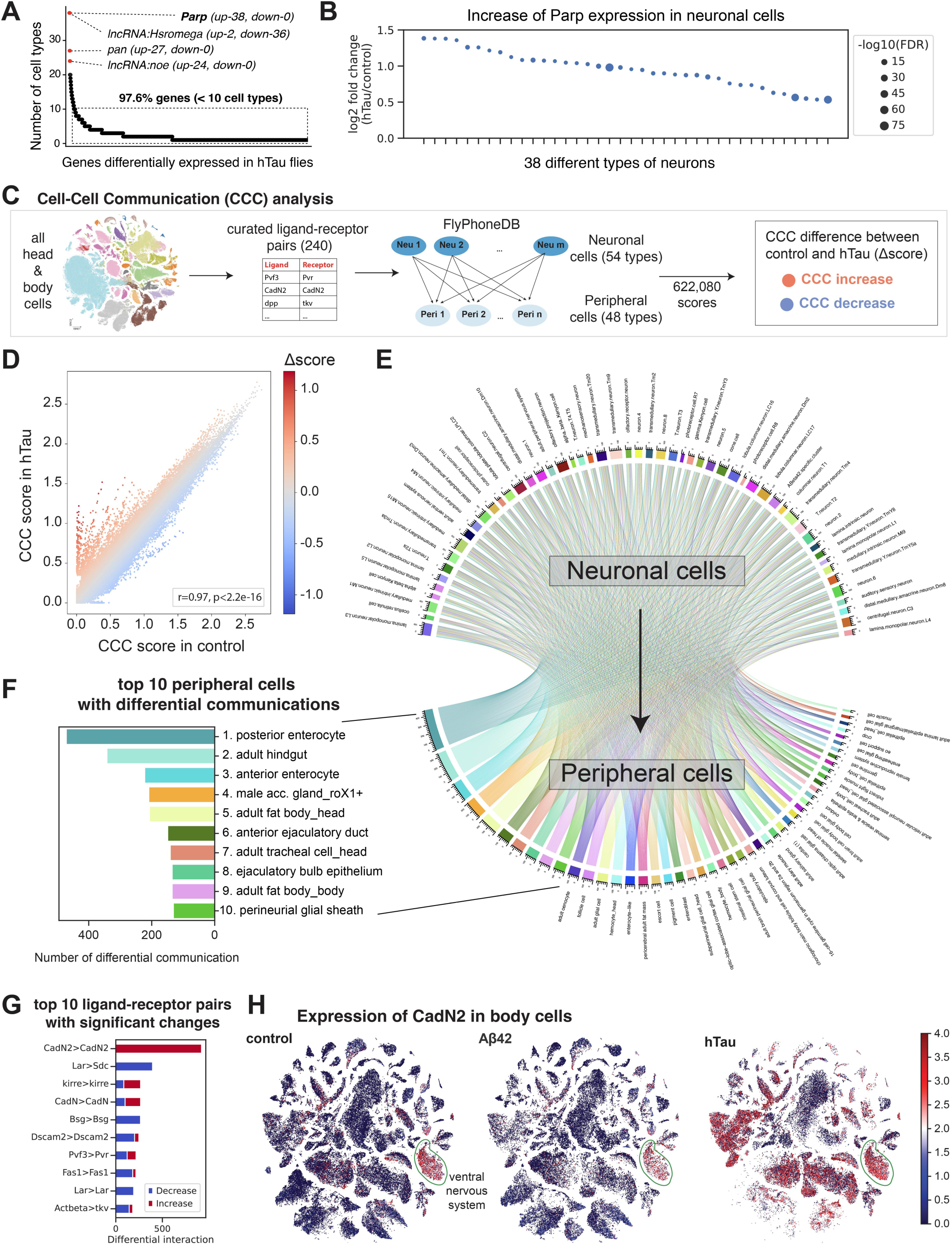
Intrinsic neuronal changes and altered brain-body communications in hTau flies. (A) Identification of genes differentially expressed in hTau neurons compared with control flies. Genes are ranked based on the number of cell types in which the genes are significantly up- or down-regulated (see Methods). (B) The expression of *Parp* is increased in 38 neuronal types. (C) Flowchart showing the cell-cell communication (CCC) analysis and the identification of altered CCC scores in hTau flies. (D) CCC scores from control and hTau flies. Each dot represents a ligand-receptor pair, which includes the ligand from a neuronal cell type and the receptor from a peripheral cell type. The Δscores are results from CCC scores in hTau flies minus CCC scores from control flies. (E) Chord diagram showing all brain-body communications that are significantly altered in hTau flies compared to control flies (Z score > 2). Periphery cell types are ranked by the number of altered CCCs. (F) Top 10 periphery cell types with the highest number of differential CCCs. (G) Top 10 ligand-receptor pairs with significant CCC changes between hTau and control flies. (H) tSNE of body cells showing the CadN2 expression in three different genotypes.

Lastly, our snRNA-seq data showed changes in the reproductive system in response to neuronal hTau expression. Hence, we examined the fecundity of control and hTau male flies and found that the number of progenies which result from mating young, female wild-type flies with 20-day hTau male flies was significantly lower than that of control (Fig. 4H). This reflects that hTau flies may have reduced reproductive function.

In summary, we found that hTau induces obvious and significant changes for several peripheral non-neuronal cell types, which regulate fat metabolism, digestion, and reproduction using snRNA-seq and functional studies.

### Neuronal expression of hTau induces accelerated aging in peripheral tissues

Interestingly, the observed changes in the periphery of hTau expressing flies – defects in lipid metabolism, gut dysplasia, and reduced fecundity (see Fig. 4) – are all associated with aging. Hence, we hypothesized that neuronal expression of hTau may induce accelerated aging phenotypes in peripheral tissues.

To test this, we considered how changes in the number of expressed transcripts (measured by the number of unique molecular identifiers, UMIs) per cell may be impacted with age in addition to considering changes in the nuclear ratio and gene expression pattern. We previously used this approach to identify cell type specific alterations associated with the aging process (Lu et al., 2023). We compared the distribution of expressed transcript numbers from the seven available conditions – three time points for control flies, two time points for Aβ42 flies, and two time points for hTau flies – and found that many different peripheral cell types from both early and late hTau flies showed a similar distribution pattern with old control flies. In contrast, both early and late Aβ42 flies showed a similar pattern with young and middle-aged control flies. These cell types include fat body cells, follicle main body cells, and enterocytes from the gut (Fig. 5A). These data support that neuronal expression of hTau accelerates transcriptional changes associated with aging in peripheral tissues. To further validate this idea, we extended our analysis to more cell types and found that age-related changes (30d-old control versus 20d-old control) and hTau-induced changes (20d-old hTau versus 20d-old control) are highly correlated for both transcript numbers and nuclear ratios (Fig. 5B-E). Changes in the transcript number are not due to the different sequencing depth of our samples because the sequencing saturations across samples with different ages, genotypes, and sexes are similar (fig. S20). Altogether, our data suggest that hTau expression has a broad impact on many peripheral tissues, particularly on cell types from the fat body, gut and reproductive systems, and it induces peripheral changes that reflect accelerated aging phenotypes.

### Intrinsic neuronal changes and brain-body communication alterations in hTau flies

What is the molecular mechanism underlying hTau-induced aging phenotypes in peripheral tissues? We reasoned that neuronal expression hTau will first cause intrinsic changes in neurons and such changes will induce accelerated aging phenotypes in peripheral tissues through the brain-body communication. Hence, we took two approaches to identify potential molecules and pathways that may contribute to these processes.

First, we performed a DEG analysis on neurons between control and hTau flies, aiming to identify genes that are broadly up- or down-regulated in response to neuronal hTau expression. We found that the majority (97.6%) of DEGs are associated with a small subset of neurons (1 to 10 cell types). Importantly, only four genes – *Poly-(ADP-ribose) polymerase (Parp)*, *lncRNA:Hsromega, pangolin* (*pan*)*, lncRNA:noe –* are differentially expressed in a larger subset of neurons (>20 cell types) (Fig. 6A). The top hit, *Parp*, is mildly but significantly up-regulated in 38 types of neurons (Fig. 6B). *Parp* encodes a chromatin-associated enzyme, Poly ADP-ribose polymerase, that modifies various nuclear proteins and is involved in several cellular processes, including DNA repair, maintenance of genomic stability, and regulation of gene expression. *Parp* upregulation in neurons may contribute to hTau-induced peripheral changes by multiple mechanisms, including by inducing inflammation (Luo and Kraus, 2012) and by altering metabolite levels like NAD+ (Cohen, 2020). Importantly, we found there is an upregulation of innate immune response genes in the fat body from hTau flies (fig. S17B), consistent with previous data showing that mild activation of Parp can induce inflammation, immunodefense, and senescence (Luo and Kraus, 2012). This may also be consistent with our previous data showing gut dysplasia occurs in hTau flies (Fig. 4G) as this phenotype can be caused by inflammatory stress (Li and Jasper, 2016). Overall, these data suggest that one mechanism, by which *Parp* upregulation is altering brain-body communication in hTau flies, may involve in the induction of inflammatory response.

Second, we performed a comprehensive cell-cell communication (CCC) analysis between all neuronal types and peripheral cell types using FlyPhoneDB, a quantification algorithm that calculates cell-cell interaction scores in *Drosophila* based on ligand-receptor pairs for the major signaling pathways (Liu et al., 2022). As many protein-protein interacting pairs that are used in the fly neural system are not included in the ligand-receptor list of the FlyPhoneDB database (196 pairs), such as CadN2-CadN2 and Lar-Sdc pairs, we expanded this list to include 44 additional pairs by searching through literature and FlyBase annotations (Table S1). Note that, although called ligand-receptor pairs, this database consists of both typical “ligand-receptor” pairs (e.g., Pvf3-Pvr, dpp-tkv), as well as atypical “surface protein-surface protein” pairs (e.g., CadN-CadN, Dscam2–Dscam2), as both can mediate cell-cell interactions.

We calculated 622,080 brain-body communications based on 54 neuronal cell types, 48 peripheral cell types, and 240 ligand-receptor pairs (Fig. 6C, see Methods). 93,524 CCC scores were significantly (p-value < 0.05) detected in hTau or control flies. Each score corresponds to one unique ligand-receptor pair, with the ligand originating from neuronal cells and the receptor from peripheral cells. Next, we compared the CCC scores from hTau and control flies. As expected, CCC scores between these two genotypes are highly correlated (R=0.97), suggesting most brain-body interactions are maintained between these two genotypes (Fig. 6D). However, hTau flies have a number of outliers that significantly deviate from the correlation plots based on Delta scores (absolute Z score > 2), reflecting possible perturbed brain-body communications (Fig. 6D, and fig. S21). Interestingly, we identified more positive outliers than negative ones, suggesting that neuronal expression of hTau creates more ectopic brain-body communications.

To characterize which peripheral cell types and which ligand-receptor pairs are perturbed in hTau flies, we first grouped the ligand-receptor pairs with significant Delta scores by their corresponding cell types (Fig. 6E). The top ten peripheral cell types with the most differential CCCs consist of eight cell types that belong to fat metabolism tissues (fat body from the head and body), digestive system (posterior enterocyte, hindgut, anterior enterocyte), as well as the reproductive system (male accessory gland, anterior ejaculatory duct, ejaculatory bulb epithelium) (Fig. 6F). This is consistent with our previous analyses showing that these three peripheral systems are significantly altered by hTau expression (Fig. 4 and 5). The remaining two cell types are perineural glia (critical for providing metabolic and nutritional support for the nervous system and a major component for the blood-brain-barrier), and tracheal cells (form a branched network of epithelial tubes that ramifies throughout the body and transports oxygen to all fly tissues), suggesting that these two cell types are also profoundly impacted by hTau expression and may contribute to peripheral aging phenotypes in hTau flies.

Next, we ranked differential CCCs by the types of ligand-receptor pairs to identify which pairs are mostly influenced by hTau (Fig. 6G). The top 10 altered ligand-receptor pairs include: CadN2–CadN2, Lar–Sdc, Kirre–Kirre, CadN–CadN, Bsg–Bsg, Dscam2–Dscam2, Pvf3–Pvr, Fas1–Fas1, Lar-Lar, and Actbeta–Tkv. Note that some of these pairs were identified due to differential gene expression in peripheral cells instead of in neuronal cells. For example, CadN2, which is a member of the cadherin superfamily and a key regulator of cell-cell adhesion during development, is normally only expressed in neurons in wild-type flies but shows wide-spread expression in many peripheral cell types of hTau flies (Fig. 6H).

In summary, our thorough CCC analyses identify the top affected peripheral cell types in hTau flies. These cell types largely regulate fat metabolism, digestion and reproduction. Further, we identified significantly changed ligand-receptor pairs that may contribute to the decline of brain-body communications during neurodegeneration, highlighting novel candidates for future functional and mechanistic characterization in AD.

## Discussion

Our AD-FCA presents a comprehensive examination of how neuronally expressed AD-associated proteins, Aβ42 or Tau, impact both neural and peripheral tissues using *Drosophila* models. Leveraging whole-organism single-nucleus transcriptomics, we profiled 624,458 nuclei across 219 cell types in AD fly models expressing human Aβ42 or Tau specifically in adult neurons. Our analysis unveiled an unexpected vulnerability of sensory neurons to Aβ42. We also found widespread peripheral tissue disruptions are caused by hTau, resembling an accelerated aging phenotype. Notably, hTau expression leads to altered lipid droplet size in the fat body, gut dysplasia, and decreased reproductive capacity. Cell-cell communication analyses highlight potential pathways mediating altered brain-body interactions. Overall, this AD-FCA offers new insights into how Aβ42 or hTau may systematically affect a whole-organism during disease progression. This dataset may pave the way for the identification of novel biomarkers and promote understanding brain-body communication in neurodegeneration.

### Different effects caused by Aβ42 or hTau

Prior studies have used either Aβ42 or hTau transgenic models to study AD-related neurodegeneration and underlying molecular mechanisms in flies, mice, and cultured cells. We note that toxicity varies between models due to differences in approaches and reagents and this hinders a systematic comparison of their toxicity and overall impacts. In order to promote our ability to compare toxicity associated with Aβ42 or Tau we employed an unified system which allows us to directly compare different effects caused by Aβ42 and hTau. By using a pan-neuronal, temperature-controlled driver (*tub-GAL80^ts^; nSyb-GAL4*) to express Aβ42 or hTau in neurons starting in the adult stage, we avoid effects seen by expressing these toxic proteins through development. To promote a fair comparison, we defined two time points per model which represented early or late stages of toxicity, and compared each model to an age-matched control. Importantly, we find that despite both Aβ42 and hTau being toxic, the consequences of this toxicity dramatically deviate between the two AD models. While Aβ42 primarily impacts the neurons, hTau induces many unexpected changes in the periphery. This is consistent with the previous studies showing Aβ42 and hTau cause neuronal death and dysfunction, respectively, when expressed in the fly eye and brain (Chouhan et al., 2016). Our findings highlight that the mechanisms that promote neurodegeneration may vary depending on the pathological cause of the disease. In AD, Aβ42-plaques likely form prior to hyperphosphorylated Tau tangles and each pathology is enriched in distinct regions of the brain (Ito et al., 2010). Our findings argue that both Aβ42 and Tau may be contributing to disease progression in succinct ways, while it is possible that the combination of the two impacts the periphery in AD.

### Cell type vulnerability in AD

In AD, cell vulnerability refers to the susceptibility of certain types, particularly neurons, to damage, dysfunction, and eventual degeneration. Studying cell vulnerability in AD is important to understand disease progression, reveal biomarkers, and identify therapeutic targets. snRNA-seq serves as an ideal method to study cell vulnerability and several recent studies have employed snRNA-seq in different AD models and human patients (Lau et al., 2020; Mathys et al., 2023; Sziraki et al., 2023; Wang et al., 2024; Wu et al., 2023). Out of necessity, most of these studies have focused on a specific brain region and identified specific cell types that are selectively vulnerable. For example, in the human prefrontal cortex, it was found that the most vulnerable cell types in AD tissues are three subtypes of SST+ inhibitory neurons (Mathys et al., 2023). In a PS19 AD mouse model, we found that granule cells in the hippocampus show the highest vulnerability among different types of neurons (Wang et al., 2024). Here, we took a novel approach and focused on comprehensively comparing peripheral sensory neurons with CNS neurons in AD models. *Drosophila* was used as it gave the opportunity to perform whole-organism snRNA-seq, a goal not as easily attainable in mammals. We revealed that sensory neurons, including auditory sensory neurons and ORNs, are among the top impacted cell types (see Fig. 2). Our functional data demonstrate that ORN loss is caused by Aβ42-expression more so than by aging, contributing to growing literature that olfaction decline is an early feature of AD (Zou et al., 2016). This model represents a new opportunity to study ORN loss specificity in human AD patients.

### ER stress for AD

Efforts to develop effective therapies to target Aβ42 and Tau in AD are ongoing while outcomes have been relatively unsatisfactory. Hence, there is an urgent need to explore alternative or complementary targets and strategies. ER stress, characterized by the accumulation of unfolded or misfolded proteins within the ER, disrupts cellular homeostasis, and can contribute to cell death (Sano and Reed, 2013). Recent evidence suggests that ER stress plays a role in the initiation and progression of AD by inducing an inflammatory response (Uddin et al., 2020). A comprehensive understanding of the ER stress machinery in AD pathology could be therapeutically relevant. ER-stress response pathway genes are largely conserved from the fly to humans (Ryoo, 2015). Here we observed that several genes in this pathway, such as *Hsc70-3/BiP*, *CaBP1/PDIA6*, and *XBP1*, are upregulated in Aβ42-expressing neurons (Fig. 3). We also found that Hsc70-3 (BiP) protein levels and apoptotic marker (Cleaved Caspase-3) are upregulated in brains from Aβ42 flies (Fig. 3 and fig. S14). Thus, both our snRNA-seq data and in vivo functional data show that the ER stress response is activated in most types of neurons in the Aβ42 brain. It will be important for future studies to further assess how ER stress contributes to Aβ42 induced neuronal loss.

### Peripheral aging phenotypes induced by neuronal expression of hTau

To identify neuronal intrinsic changes that may contribute to peripheral aging, we focused on candidate genes that are significantly and broadly changed in hTau-expressing neurons and identified four genes: *Poly-(ADP-ribose) polymerase* (*Parp)*, *pangolin* (*pan*), and two long non-coding RNA genes (*Hsromega* and *noe*). Of these, we focus on *Parp* as it was the most impacted and it may contribute to relevant peripheral aging phenotypes. Moderate activation of Parp induces inflammatory signals, while strong activation will cause parthanatos cell death (Luo and Kraus, 2012). Considering two facts that hTau does not cause obvious neuronal loss (Fig. 2) and that *Parp* is mildly increased in hTau-expressing neurons (Fig. 6B), it is conceivable that upregulation of *Parp* may induce systemic inflammation and immune-responses and lead to peripheral aging. This idea is supported by the findings in hTau flies, such as an upregulation of innate immune response genes in the fat body and gut dysplasia phenotypes, which can be caused by inflammatory stress, (Fig. 4). In addition, an upregulation of *Parp* may also impact peripheral aging by depleting NAD+ and its precursor storage in the body. It has been shown that Parp requires NAD+ to synthesize PAR chains during DNA damage response (Murata et al., 2019). These PAR chains recruit other DNA repair proteins to the site of damage. Hence, Parp activation will consume NAD+, and NAD+ depletion is associated with aging in many cell types. Increased Parp activity has been associated with different neurodegenerative diseases (Hu et al., 2023; Kam et al., 2018; McGurk et al., 2018), but its mechanistic role in AD and AD-induced peripheral changes has not been characterized so far. Note that investigations into the other genes identified – *pan, lncRNA:Hsromega* and *lncRNA:noe* – would add further insight into how peripheral changes may be driven by neuronal expression of hTau. In summary, we have identified several candidate genes that may, individually or in combination, contribute to hTau-induced peripheral aging.

### Comprehensive cell-cell communication (CCC) analysis

To further characterize hTau-induced aging phenotypes in peripheral tissues, we performed a comprehensive survey on brain-body communications using FlyPhoneDB-based CCC analysis. Existing research in the brain-body communication field has predominantly concentrated on specific pairs of cell types connecting the brain and the body, often centered on the gut-brain axis. We expanded upon previous work and analyzed all cell types throughout the whole organism. By leveraging the whole-organism snRNA-seq from both wild-type and AD-related neurodegeneration models, we revealed previously uncharted pathways and infrequent interactions. Indeed, our thorough analysis not only revealed known brain-body communications, like neuron-glia, brain-gut, and brain-fat body interactions, but also uncovered new interactions, such as brain-tracheal and brain-reproductive systems (Fig. 6E, F). At the molecular level, we also identified many ligand-receptor pairs that show prevalent changes and will increase or decrease communication between certain neurons and peripheral cells. For example, we found that the *cadherin-N2* (*CadN2)*, which is normally expressed in neurons, is broadly upregulated in many peripheral cells of hTau flies. CadN2 proteins localize to the cell surface and form homophilic interactions, connecting cells. Hence, an aberrant upregulation of this protein in the periphery will disrupt normal cell-cell interactions. It is conceivable that these aberrant interactions may extend beyond brain-peripheral communications to encompass peripheral-peripheral interactions, an emerging area of interest in the study of inter-organ communication during aging. In summary, our AD-FCA and CCC results offer an opportunity to identify novel mechanisms by which organs communicate during aging and in AD models.

## Acknowledgments

We thank the Bellen lab and Li lab members for their constructive advice on this project. We thank Drs. Josh Shulman, Yi Zhu, and Hui Zheng for supporting and giving feedback to this project. We thank Dr. Jason Tennessen for sharing Ldh fly lines and Dr. Hermann Steller for sharing the BiP antibody.

## Author contributions

Conceptualization: Y-J.P., T-C.L., H.J.B., H.L.

snRNA-seq: Y-J.P., Y.Q.

Computational analysis: Y-J.P., T-C.L., T.J.

Transgenic fly: S.Y.

Data portal and AD-FCA website: H.L., J.C.

In vivo data validation: Y-J.P., T.J., L.R., C-Y.L., E.H., C.K., A-L.H., Y.Q., H.L.

Writing: Y-J.P., T-C.L., L.D.G., H.L.

Review and Editing: All authors. Supervision: H.J.B., H.L.

Funding Acquisition: H.J.B., L.D.G., H.L.

## Data and materials availability

Raw FASTQ files and processed h5ad files, including expression matrices and cell-type annotations, have been submitted to NCBI/GEO (currently being curated and will be released after NCBI/GEO curation; total size is over 4TB). Codes for data processing and analysis are available from github: https://github.com/MeshifLu/ADFCA. All annotated data are available for visualization, download, and custom analysis at https://hongjielilab.org/adfca.

## Supplementary Materials

Materials and Methods

Figures and Legends S1 to S21

Table S1

## References

Almanzar, N., Antony, J., Baghel, A.S., Bakerman, I., Bansal, I., Barres, B.A., Beachy, P.A., Berdnik, D., Bilen, B., Brownfield, D., et al. (2020). A single-cell transcriptomic atlas characterizes ageing tissues in the mouse. Nature 583, 590–595. 10.1038/s41586-020-2496-1.

Bawa, S., Brooks, D.S., Neville, K.E., Tipping, M., Sagar, M.A., Kollhoff, J.A., Chawla, G., Geisbrecht, B.V., Tennessen, J.M., Eliceiri, K.W., et al. (2020). Drosophila TRIM32 cooperates with glycolytic enzymes to promote cell growth. eLife 9, e52358. 10.7554/eLife.52358.

Bettcher, B.M., Tansey, M.G., Dorothée, G., and Heneka, M.T. (2021). Peripheral and central immune system crosstalk in Alzheimer disease — a research prospectus. Nat Rev Neurol 17, 689–701. 10.1038/s41582-021-00549-x.

Bischof, J., Maeda, R.K., Hediger, M., Karch, F., and Basler, K. (2007). An optimized transgenesis system for Drosophila using germ-line-specific φC31 integrases. Proceedings of the National Academy of Sciences 104, 3312–3317. 10.1073/pnas.0611511104.

Brand, A.H., and Perrimon, N. (1993). Targeted gene expression as a means of altering cell fates and generating dominant phenotypes. Development 118, 401–415. 10.1242/dev.118.2.401.

Chouhan, A.K., Guo, C., Hsieh, Y.-C., Ye, H., Senturk, M., Zuo, Z., Li, Y., Chatterjee, S., Botas, J., Jackson, G.R., et al. (2016). Uncoupling neuronal death and dysfunction in Drosophila models of neurodegenerative disease. Acta Neuropathologica Communications 4, 62. 10.1186/s40478-016-0333-4.

Claes, C., Danhash, E.P., Hasselmann, J., Chadarevian, J.P., Shabestari, S.K., England, W.E., Lim, T.E., Hidalgo, J.L.S., Spitale, R.C., Davtyan, H., et al. (2021). Plaque-associated human microglia accumulate lipid droplets in a chimeric model of Alzheimer’s disease. Molecular Neurodegeneration 16, 50. 10.1186/s13024-021-00473-0.

Cohen, M.S. (2020). Interplay between compartmentalized NAD+ synthesis and consumption: a focus on the PARP family. Genes Dev. 34, 254–262. 10.1101/gad.335109.119.

Davie, K., Janssens, J., Koldere, D., De Waegeneer, M., Pech, U., Kreft, Ł., Aibar, S., Makhzami, S., Christiaens, V., Bravo González-Blas, C., et al. (2018). A Single-Cell Transcriptome Atlas of the Aging Drosophila Brain. Cell 174, 982–998.e20. 10.1016/j.cell.2018.05.057.

Doty, R.L. (2012). Olfactory dysfunction in Parkinson disease. Nat Rev Neurol 8, 329–339. 10.1038/nrneurol.2012.80.

Duff, K., McCaffrey, R.J., and Solomon, G.S. (2002). The Pocket Smell Test. JNP 14, 197–201. 10.1176/jnp.14.2.197.

Fang, Z., Liu, X., and Peltz, G. (2023). GSEApy: a comprehensive package for performing gene set enrichment analysis in Python. Bioinformatics 39, btac757. 10.1093/bioinformatics/btac757.

Fleming, S.J., Chaffin, M.D., Arduini, A., Akkad, A.-D., Banks, E., Marioni, J.C., Philippakis, A.A., Ellinor, P.T., and Babadi, M. (2023). Unsupervised removal of systematic background noise from droplet-based single-cell experiments using CellBender. Nat Methods 1–13. 10.1038/s41592-023-01943-7.

Germain, P.-L., Lun, A., Meixide, C.G., Macnair, W., and Robinson, M.D. (2022). Doublet identification in single-cell sequencing data using *scDblFinder*. 10.12688/f1000research.73600.2.

Holsopple, J.M., Cook, K.R., and Popodi, E.M. (2022a). Enteroendocrine cell expression of split-GAL4 drivers bearing regulatory sequences associated with panneuronally expressed genes in Drosophila melanogaster. microPublication Biology 10.17912/micropub.biology.000628.

Holsopple, J.M., Cook, K.R., and Popodi, E.M. (2022b). Identification of novel split-GAL4 drivers for the characterization of enteroendocrine cells in the Drosophila melanogaster midgut. G3 Genes|Genomes|Genetics 12, jkac102. 10.1093/g3journal/jkac102.

Hu, M.-L., Pan, Y.-R., Yong, Y.-Y., Liu, Y., Yu, L., Qin, D.-L., Qiao, G., Law, B.Y.-K., Wu, J.-M., Zhou, X.-G., et al. (2023). Poly (ADP-ribose) polymerase 1 and neurodegenerative diseases: Past, present, and future. Ageing Research Reviews 91, 102078. 10.1016/j.arr.2023.102078.

Iijima, K., Liu, H.-P., Chiang, A.-S., Hearn, S.A., Konsolaki, M., and Zhong, Y. (2004). Dissecting the pathological effects of human Aβ40 and Aβ42 in Drosophila: A potential model for Alzheimer’s disease. Proceedings of the National Academy of Sciences 101, 6623–6628. 10.1073/pnas.0400895101.

Ioannou, M.S., Jackson, J., Sheu, S.-H., Chang, C.-L., Weigel, A.V., Liu, H., Pasolli, H.A., Xu, C.S., Pang, S., Matthies, D., et al. (2019). Neuron-Astrocyte Metabolic Coupling Protects against Activity-Induced Fatty Acid Toxicity. Cell 177, 1522–1535.e14. 10.1016/j.cell.2019.04.001.

Ito, K., Ahadieh, S., Corrigan, B., French, J., Fullerton, T., and Tensfeldt, T. (2010). Disease progression meta-analysis model in Alzheimer’s disease. Alzheimer’s & Dementia 6, 39–53. 10.1016/j.jalz.2009.05.665.

Jones, W.D., Cayirlioglu, P., Grunwald Kadow, I., and Vosshall, L.B. (2007). Two chemosensory receptors together mediate carbon dioxide detection in Drosophila. Nature 445, 86–90. 10.1038/nature05466.

Kam, T.-I., Mao, X., Park, H., Chou, S.-C., Karuppagounder, S.S., Umanah, G.E., Yun, S.P., Brahmachari, S., Panicker, N., Chen, R., et al. (2018). Poly(ADP-ribose) drives pathologic α-synuclein neurodegeneration in Parkinson’s disease. Science 362, eaat8407. 10.1126/science.aat8407.

Kanehisa, M., and Goto, S. (2000). KEGG: Kyoto Encyclopedia of Genes and Genomes. Nucleic Acids Research 28, 27–30. 10.1093/nar/28.1.27.

Katheder, N.S., Browder, K.C., Chang, D., De Maziere, A., Kujala, P., van Dijk, S., Klumperman, J., Lu, T.-C., Li, H., Lai, Z., et al. (2023). Nicotinic acetylcholine receptor signaling maintains epithelial barrier integrity. eLife 12, e86381. 10.7554/eLife.86381.

Klopfenstein, D.V., Zhang, L., Pedersen, B.S., Ramírez, F., Warwick Vesztrocy, A., Naldi, A., Mungall, C.J., Yunes, J.M., Botvinnik, O., Weigel, M., et al. (2018). GOATOOLS: A Python library for Gene Ontology analyses. Sci Rep 8, 10872. 10.1038/s41598-018-28948-z.

Korsunsky, I., Millard, N., Fan, J., Slowikowski, K., Zhang, F., Wei, K., Baglaenko, Y., Brenner, M., Loh, P., and Raychaudhuri, S. (2019). Fast, sensitive and accurate integration of single-cell data with Harmony. Nat Methods 16, 1289–1296. 10.1038/s41592-019-0619-0.

Lau, S.-F., Cao, H., Fu, A.K.Y., and Ip, N.Y. (2020). Single-nucleus transcriptome analysis reveals dysregulation of angiogenic endothelial cells and neuroprotective glia in Alzheimer’s disease. Proceedings of the National Academy of Sciences 117, 25800–25809. 10.1073/pnas.2008762117.

Li, G., and Hidalgo, A. (2020). Adult Neurogenesis in the Drosophila Brain: The Evidence and the Void. International Journal of Molecular Sciences 21, 6653. 10.3390/ijms21186653.

Li, H., and Jasper, H. (2016). Gastrointestinal stem cells in health and disease: from flies to humans. Disease Models & Mechanisms 9, 487–499. 10.1242/dmm.024232.

Li, H., Janssens, J., De Waegeneer, M., Kolluru, S.S., Davie, K., Gardeux, V., Saelens, W., David, F.P.A., Brbić, M., Spanier, K., et al. (2022). Fly Cell Atlas: A single-nucleus transcriptomic atlas of the adult fruit fly. Science 375, eabk2432. 10.1126/science.abk2432.

Liu, L., Zhang, K., Sandoval, H., Yamamoto, S., Jaiswal, M., Sanz, E., Li, Z., Hui, J., Graham, B.H., Quintana, A., et al. (2015). Glial Lipid Droplets and ROS Induced by Mitochondrial Defects Promote Neurodegeneration. Cell 160, 177–190. 10.1016/j.cell.2014.12.019.

Liu, L., MacKenzie, K.R., Putluri, N., Maletić-Savatić, M., and Bellen, H.J. (2017). The Glia-Neuron Lactate Shuttle and Elevated ROS Promote Lipid Synthesis in Neurons and Lipid Droplet Accumulation in Glia via APOE/D. Cell Metabolism 26, 719–737.e6. 10.1016/j.cmet.2017.08.024.

Liu, Y., Li, J.S.S., Rodiger, J., Comjean, A., Attrill, H., Antonazzo, G., Brown, N.H., Hu, Y., and Perrimon, N. (2022). FlyPhoneDB: an integrated web-based resource for cell–cell communication prediction in Drosophila. Genetics 220, iyab235. 10.1093/genetics/iyab235.

Lu, T.-C., Brbić, M., Park, Y.-J., Jackson, T., Chen, J., Kolluru, S.S., Qi, Y., Katheder, N.S., Cai, X.T., Lee, S., et al. (2023). Aging Fly Cell Atlas identifies exhaustive aging features at cellular resolution. Science 380, eadg0934. 10.1126/science.adg0934.

Luo, X., and Kraus, W.L. (2012). On PAR with PARP: cellular stress signaling through poly(ADP-ribose) and PARP-1. Genes Dev. 26, 417–432. 10.1101/gad.183509.111.

Marschallinger, J., Iram, T., Zardeneta, M., Lee, S.E., Lehallier, B., Haney, M.S., Pluvinage, J.V., Mathur, V., Hahn, O., Morgens, D.W., et al. (2020). Lipid-droplet-accumulating microglia represent a dysfunctional and proinflammatory state in the aging brain. Nat Neurosci 23, 194–208. 10.1038/s41593-019-0566-1.

Mathys, H., Peng, Z., Boix, C.A., Victor, M.B., Leary, N., Babu, S., Abdelhady, G., Jiang, X., Ng, A.P., Ghafari, K., et al. (2023). Single-cell atlas reveals correlates of high cognitive function, dementia, and resilience to Alzheimer’s disease pathology. Cell 186, 4365–4385.e27. 10.1016/j.cell.2023.08.039.

McGurk, L., Gomes, E., Guo, L., Mojsilovic-Petrovic, J., Tran, V., Kalb, R.G., Shorter, J., and Bonini, N.M. (2018). Poly(ADP-Ribose) Prevents Pathological Phase Separation of TDP-43 by Promoting Liquid Demixing and Stress Granule Localization. Molecular Cell 71, 703–717.e9. 10.1016/j.molcel.2018.07.002.

McLaughlin, C.N., Qi, Y., Quake, S.R., Luo, L., and Li, H. (2022). Isolation and RNA sequencing of single nuclei from Drosophila tissues. STAR Protocols 3, 101417. 10.1016/j.xpro.2022.101417.

Menengiç, K.N., Güngör, F., Ovacık, U., Çınar, N., and Yeldan, İ. (2020). Relation of Respiratory Function and Functional Capacity, Body Composition in Individuals with Alzheimer’s Disease. European Respiratory Journal 56. 10.1183/13993003.congress-2020.2973.

Micchelli, C.A., and Perrimon, N. (2006). Evidence that stem cells reside in the adult Drosophila midgut epithelium. Nature 439, 475–479. 10.1038/nature04371.

Moulton, M.J., Barish, S., Ralhan, I., Chang, J., Goodman, L.D., Harland, J.G., Marcogliese, P.C., Johansson, J.O., Ioannou, M.S., and Bellen, H.J. (2021). Neuronal ROS-induced glial lipid droplet formation is altered by loss of Alzheimer’s disease–associated genes. Proceedings of the National Academy of Sciences 118, e2112095118. 10.1073/pnas.2112095118.

Murata, M.M., Kong, X., Moncada, E., Chen, Y., Imamura, H., Wang, P., Berns, M.W., Yokomori, K., and Digman, M.A. (2019). NAD+ consumption by PARP1 in response to DNA damage triggers metabolic shift critical for damaged cell survival. MBoC 30, 2584–2597. 10.1091/mbc.E18-10-0650.

Ohlstein, B., and Spradling, A. (2006). The adult Drosophila posterior midgut is maintained by pluripotent stem cells. Nature 439, 470–474. 10.1038/nature04333.

Osman, R., Tacnet-Delorme, P., Kleman, J.-P., Millet, A., and Frachet, P. (2017). Calreticulin Release at an Early Stage of Death Modulates the Clearance by Macrophages of Apoptotic Cells. Frontiers in Immunology 8. .

Özel, M.N., Simon, F., Jafari, S., Holguera, I., Chen, Y.-C., Benhra, N., El-Danaf, R.N., Kapuralin, K., Malin, J.A., Konstantinides, N., et al. (2021). Neuronal diversity and convergence in a visual system developmental atlas. Nature 589, 88–95. 10.1038/s41586-020-2879-3.

Pacyna, R.R., Han, S.D., Wroblewski, K.E., McClintock, M.K., and Pinto, J.M. (2023). Rapid olfactory decline during aging predicts dementia and GMV loss in AD brain regions. Alzheimer’s & Dementia 19, 1479–1490. 10.1002/alz.12717.

Palladino, M.J., Hadley, T.J., and Ganetzky, B. (2002). Temperature-Sensitive Paralytic Mutants Are Enriched For Those Causing Neurodegeneration in Drosophila. Genetics 161, 1197–1208. 10.1093/genetics/161.3.1197.

Park, Y.-J., Kim, S., Shim, H.-P., Park, J.H., Lee, G., Kim, T.-Y., Jo, M.-C., Kwon, A.-Y., Lee, M., Lee, S., et al. (2021). Phosphatidylserine synthase plays an essential role in glia and affects development, as well as the maintenance of neuronal function. iScience 24. 10.1016/j.isci.2021.102899.

Piper, M.D.W., and Partridge, L. (2018). *Drosophila* as a model for ageing. Biochimica et Biophysica Acta (BBA) - Molecular Basis of Disease 1864, 2707–2717. 10.1016/j.bbadis.2017.09.016.

Prüßing, K., Voigt, A., and Schulz, J.B. (2013). Drosophila melanogaster as a model organism for Alzheimer’s disease. Molecular Neurodegeneration 8, 35. 10.1186/1750-1326-8-35.

Roux, A.E., Yuan, H., Podshivalova, K., Hendrickson, D., Kerr, R., Kenyon, C., and Kelley, D. (2023). Individual cell types in *C. elegans* age differently and activate distinct cell-protective responses. Cell Reports 42, 112902. 10.1016/j.celrep.2023.112902.

Ryoo, H.D. (2015). Drosophila as a model for unfolded protein response research. BMB Reports 48, 445–453. 10.5483/BMBRep.2015.48.8.099.

Ryoo, H.D., Domingos, P.M., Kang, M., and Steller, H. (2007). Unfolded protein response in a Drosophila model for retinal degeneration. The EMBO Journal 26, 242–252. 10.1038/sj.emboj.7601477.

Sano, R., and Reed, J.C. (2013). ER stress-induced cell death mechanisms. Biochimica et Biophysica Acta (BBA) - Molecular Cell Research 1833, 3460–3470. 10.1016/j.bbamcr.2013.06.028.

Scheltens, P., Blennow, K., Breteler, M.M.B., Strooper, B. de, Frisoni, G.B., Salloway, S., and Flier, W.M.V. der (2016). Alzheimer’s disease. The Lancet 388, 505–517. 10.1016/S0140-6736(15)01124-1.

Scheltens, P., De Strooper, B., Kivipelto, M., Holstege, H., Chételat, G., Teunissen, C.E., Cummings, J., and van der Flier, W.M. (2021). Alzheimer’s disease. The Lancet 397, 1577–1590. 10.1016/S0140-6736(20)32205-4.

Siletti, K., Hodge, R., Mossi Albiach, A., Lee, K.W., Ding, S.-L., Hu, L., Lönnerberg, P., Bakken, T., Casper, T., Clark, M., et al. (2023). Transcriptomic diversity of cell types across the adult human brain. Science 382, eadd7046. 10.1126/science.add7046.

Stampfer, M.J. (2006). Cardiovascular disease and Alzheimer’s disease: common links. Journal of Internal Medicine 260, 211–223. 10.1111/j.1365-2796.2006.01687.x.

Stevens, M., Nanou, A., Terstappen, L.W.M.M., Driemel, C., Stoecklein, N.H., and Coumans, F.A.W. (2022). StarDist Image Segmentation Improves Circulating Tumor Cell Detection. Cancers (Basel) 14, 2916. 10.3390/cancers14122916.

Sziraki, A., Lu, Z., Lee, J., Banyai, G., Anderson, S., Abdulraouf, A., Metzner, E., Liao, A., Banfelder, J., Epstein, A., et al. (2023). A global view of aging and Alzheimer’s pathogenesis-associated cell population dynamics and molecular signatures in human and mouse brains. Nat Genet 55, 2104–2116. 10.1038/s41588-023-01572-y.

Tatar, M., Post, S., and Yu, K. (2014). Nutrient control of Drosophila longevity. Trends in Endocrinology & Metabolism 25, 509–517. 10.1016/j.tem.2014.02.006.

Tian, Q., Bilgel, M., Moghekar, A.R., Ferrucci, L., and Resnick, S.M. (2022). Olfaction, Cognitive Impairment, and PET Biomarkers in Community-Dwelling Older Adults. Journal of Alzheimer’s Disease 86, 1275–1285. 10.3233/JAD-210636.

Uddin, Md.S., Tewari, D., Sharma, G., Kabir, Md.T., Barreto, G.E., Bin-Jumah, M.N., Perveen, A., Abdel-Daim, M.M., and Ashraf, G.M. (2020). Molecular Mechanisms of ER Stress and UPR in the Pathogenesis of Alzheimer’s Disease. Mol Neurobiol 57, 2902–2919. 10.1007/s12035-020-01929-y.

Venken, K.J.T., He, Y., Hoskins, R.A., and Bellen, H.J. (2006). P[acman]: A BAC Transgenic Platform for Targeted Insertion of Large DNA Fragments in D. melanogaster. Science 314, 1747–1751. 10.1126/science.1134426.

Vogt, N.M., Kerby, R.L., Dill-McFarland, K.A., Harding, S.J., Merluzzi, A.P., Johnson, S.C., Carlsson, C.M., Asthana, S., Zetterberg, H., Blennow, K., et al. (2017). Gut microbiome alterations in Alzheimer’s disease. Sci Rep 7, 13537. 10.1038/s41598-017-13601-y.

Wang, B., Martini-Stoica, H., Qi, C., Lu, T.-C., Wang, S., Xiong, W., Qi, Y., Xu, Y., Sardiello, M., Li, H., et al. (2024). TFEB–vacuolar ATPase signaling regulates lysosomal function and microglial activation in tauopathy. Nat Neurosci 27, 48–62. 10.1038/s41593-023-01494-2.

Wittmann, C.W., Wszolek, M.F., Shulman, J.M., Salvaterra, P.M., Lewis, J., Hutton, M., and Feany, M.B. (2001). Tauopathy in Drosophila: Neurodegeneration Without Neurofibrillary Tangles. Science 293, 711–714. 10.1126/science.1062382.

Wolf, F.A., Angerer, P., and Theis, F.J. (2018). SCANPY: large-scale single-cell gene expression data analysis. Genome Biology 19, 15. 10.1186/s13059-017-1382-0.

Wu, T., Deger, J.M., Ye, H., Guo, C., Dhindsa, J., Pekarek, B.T., Al-Ouran, R., Liu, Z., Al-Ramahi, I., Botas, J., et al. (2023). Tau polarizes an aging transcriptional signature to excitatory neurons and glia. eLife 12, e85251. 10.7554/eLife.85251.

Xiong, J., Kang, S.S., Wang, Z., Liu, X., Kuo, T.-C., Korkmaz, F., Padilla, A., Miyashita, S., Chan, P., Zhang, Z., et al. (2022). FSH blockade improves cognition in mice with Alzheimer’s disease. Nature 603, 470–476. 10.1038/s41586-022-04463-0.

Zhang, W., Zhang, S., Yan, P., Ren, J., Song, M., Li, J., Lei, J., Pan, H., Wang, S., Ma, X., et al. (2020). A single-cell transcriptomic landscape of primate arterial aging. Nat Commun 11, 2202. 10.1038/s41467-020-15997-0.

Zou, Y., Lu, D., Liu, L., Zhang, H., and Zhou, Y. (2016). Olfactory dysfunction in Alzheimer’s disease. NDT 12, 869–875. 10.2147/NDT.S104886.

