## Supplemental Figures for "Whole organism snRNA-seq reveals systemic peripheral changes in Alzheimer’s Disease fly models"

#### Materials and Methods

##### *Drosophila* husbandry, fly lines, and genotypes

Flies were maintained on standard cornmeal-molasses medium with a 12-hour light–dark cycle. *tub-GAL80$^{ts}$; nSyb-GAL4* virgin flies crossed to *UAS-empty*, *UAS-Aβ42*, or *UAS-hTau* transgenic male flies. Each UAS line was combined with *Gr21a:syt-RFP* or *Ldh-GFP* flies for Gr21a ORN and Ldh related experiments. To avoid developmental expression-caused defects, flies were kept at 18ºC during cross and developmental process. Progenies were allowed to mate for the first 1-4 days at 18ºC, and moved to 29ºC to express transgenes in pan-neurons from the adult stage. **Table** below details the fly lines used.

| Fly lines | genotype | Source/reference | notes |
|---|---|---|---|
| *tub-GAL80$^{ts}$* | *w*;P{tubP-GAL80ts}10;TM2/TM6 B, Tb1* | BDSC #7108 | |
| *nSyb-GAL4* | *y1 w*; P{nSyb-GAL4.S}3* | BDSC #51635 | |
| *UAS-empty* | *w[1118];;UAS-empty/TM6* | (Moulton et al., 2021) | pGW vector injected to VK33 |
| *UAS-Aβ42 (II)* | *yw;P{UAS-APP.Abeta42.B}m26a (II)* | (Chouhan et al., 2016) | Used for most of the crosses |
| *UAS-Aβ42 (III)* | *yw;; P{UAS-APP.Abeta42.B}m17a (III)* | (Chouhan et al., 2016) | Only used for combining with *Ldh-GFP (II)* |
| *UAS-hTau* | *w[1118];; UAS-humanTau$^{0N4R}$/TM3, Sb[1]* | (Wittmann et al., 2001) | Isoform 0N4R |
| *UAS-ssp.3XHA* | *UAS-ssp.3xHA (II)* | This study | See methods |
| *Gr21a:syt-RFP* | *Gr21a:syt-RFP (III)* | (Jones et al., 2007) | |
| *Ldh-GFP* | *w*; P{Ldh-optGFP}attP40* | BDSC #94704 | |

##### Generation of *UAS-ssp.3XHA* transgenic fly lines

A transgenic construct that allows overexpression of a neutral 3xHA polypeptide with a signal sequence under the control of the UAS/GAL4 system (Brand and Perrimon, 1993) was generated as follows. We PCR amplified the rat Preproenkephalin signal sequence (MAQFLRLCIWLLALGSCLLATVQA) with primers to add a *Drosophila* codon optimized 3xHA tag (YPYDVPDYAYPYDVPDYAYPYDVPDYA) at its C-terminus from pUAST-SSP::Abeta42 (Iijima et al., 2004) (a gift from Dr. Koichi Iijima). Restriction enzyme sites (BglII and XhoI) were also added for cloning purposes. The amplified fragment was first cloned into pGEM-T Easy plasmid (Promega) via TA cloning and subsequently shuttled to the pUAST.attB plasmid (Bischof et al., 2007). A Sanger sequence validated clone was injected into the VK37 docking site on the 2nd chromosome (Venken et al., 2006).

**Sample collection, lifespan and negative geotaxis assays**

Progenies were collected 3 days after the first fly hatched at 18°C. Flies were allowed to mate for 1 more day at 18 ℃. Then male and female flies were separated into different cages (150 flies per cage) and incubated at 29°C. Fly food was changed every 2 days. For sample collection, fly heads and bodies were dissected at different ages, 10d, 20d and 30d, put into 1.5ml RNAase free Eppendorf tube, flash-frozen using liquid nitrogen, and then stored at –80℃. We put 50 heads or 15 bodies in each tube. For longevity assay, dead or censored flies was recorded every 2 days. Female and male flies were counted separately. Survival rates were calculated from the total population. For negative geotaxis assays, we used a climbing kit (used in (Park et al., 2021)) to compare up to 5 groups at once. 0, 10, 20, or 30-day-old flies were transferred to an empty vial (20 flies per genotype per set). The climbing kit was gently tapped to force all flies to the bottom and then videotaped for 10 seconds. Flies climbed 10 times per set. Climbing height at 10 seconds was measured by the Tracker software (Brown, 2008) (https://tracker.physlets.org/) for the final data. Female and male flies were separated for this assay. 5 sets of 0d (N=100, 40 female and 60 male per genotype), 8 sets of 10d (N=160, 80 female and 80 male per genotype), 10 sets of 20d (N=200, 100 female and 100 male per genotype), 10 sets of 30d (N=200, 100 female and 100 male per genotype). Survival curves and negative geotaxis plots were generated in Graphpad Prism.

**Single-nucleus RNA sequencing**

Single-nucleus suspensions were prepared following the protocol we described previously (McLaughlin et al., 2022). Next, we used the BD AriaIII FACS sorter for collecting nuclei. Nuclei were stained by Hoechst-33342 on ice (1:1000; >5min). Hoechst+ nuclei were collected during sorting. Since polyploidy is common for many tissue types in *Drosophila*, we observed different populations of nuclei according to DNA content (Hoechst signal). In order to include all cell populations with different nuclear sizes, we included all nuclear populations from the FACS in samples for single-nucleus RNA sequencing. Individual nuclei were collected into one 1.5ml RNAase free Eppendorf tube with 200μl 1x PBS with 0.5% BSA as the receiving buffer (RNase inhibitor added). For each 10x Genomics run, 200k nuclei were collected. Nuclei were spun down for 10 min at 950g at 4℃, and then resuspended using 80μl or desired amount of 1x PBS with 0.5% BSA (RNase inhibitor added). 2μl of nucleus suspension was used for counting the nuclei with hemocytometers to calculate the concentration. We loaded 50-60K nuclei to the 10x controller to target >20k nuclei for each channel.

Next, we performed snRNA-seq using the 10x Genomics platform with the Chromium Next GEM Single Cell 3' HT(high-throughput) Reagent Kits v3.1 (Dual Index) with following settings. All PCR reactions were performed using the BioRad C1000 Touch Thermal cycler with 96-Deep Well Reaction Module. The recommended cycle numbers from 10x protocol were used for cDNA amplification and sample index PCR. As per 10x protocol, 1:10 dilutions of amplified cDNA and final libraries were evaluated on a bioanalyzer. The final library was sent to

Novogene Corporation Inc. for Illumina NovaSeq PE150 S4 lane sequencing with the dual index configuration Read 1 28 cycles, Index 1 (i7) 10 cycles, Index 2 (i5) 10 cycles and Read 2 90 cycles. A PhiX control library was spiked in at 0.2 to 1% concentration. The sequencing depth is about 30-40K reads per nucleus.

**snRNA-seq data processing**

Raw snRNA-seq data, in the form of FASTQ files, underwent alignment to the *Drosophila melanogaster* reference genome (FlyBase release 6.31) using the Cell Ranger software (v6.1.2), employing a pre-mRNA gene transfer format (GTF) file generated by the FCA algorithm (Li et al., 2022). Subsequent steps involved the removal of ambient RNA contamination via CellBender (Fleming et al., 2023) and the identification and exclusion of potential doublet cells using scDblFinder (Germain et al., 2022). Quality control criteria necessitated the elimination of cells exhibiting fewer than 200 genes or 500 UMIs. Genes detected in fewer than three nuclei were removed from our analysis. Furthermore, cells with gene or UMI counts exceeding five median absolute deviations from the median were also excluded from analysis. Additionally, cells harboring over 5% of mitochondrial transcripts were eliminated. The majority of the snRNA-seq data analysis was conducted using the Scanpy package (v1.8.2) (Wolf et al., 2018).

**Detection of vacuoles in the brain**

To detect vacuoles in the brain, fly heads were fixed in fresh Carnoy's solution (ethanol:chloroform:acetic acid at 6:3:1) at 4°C for at least 24 hours, washed, serially dehydrated with ethanol, and paraffin-embedded using standard histological procedures (Palladino et al., 2002). Serial 5 µm sections were stained with hematoxylin and eosin and examined under a light microscope. Images were taken with a Nikon NiU Eclipse Upright Microscope.

**Whole brain dissection and immunofluorescence**

Fly brains were dissected in 1X PBS using Dumont #55 forceps (FST, 11255-20), placed into 0.2ml PCR strip tubes, and fixed with 4% paraformaldehyde (PFA) in 1X PBS for 20 min at room temperature (RT). The brains were thoroughly washed with 0.3% PBST (1X PBS + 0.3% Triton-X100) and blocked with a brain blocking buffer (5% normal goat serum (NGS) in 0.3% PBST) for 2 hours at RT. After blocking, the brains were incubated with primary antibodies (see below) in the brain blocking buffer at 4ºC overnight. The brains were thoroughly rinsed with 0.2% PBST and incubated with secondary antibodies (see below) in the brain blocking buffer at RT for 2 hours. The brains were thoroughly rinsed with 0.3% PBST and 1X PBS, and mounted with SlowFade™ Gold Antifade Mountant (Thermo Fisher, S36936). Images were obtained with Leica STELLARIS 5 confocal microscope as Z series with the same interval. Z series images were merged by ImageJ (Image-Stacks-Z projection-Max Intensity or SUM slices), and then the signal intensity and area were measured using ImageJ. Quantification graphs were generated in GraphPad Prism. P-values were computed using the parametric unpaired t-test. For Ldh-GFP brains, endogenous GFP signal was imaged without antibody staining.

*Primary antibodies*

mouse anti-Aβ42 (1:100, BioLegend, #805509)

mouse anti-hyperphosphorylated-Tau (AT8) (1:500, ThermoFisher, #MN1020)
rat anti-NCad (1:40, DSHB, DN-Ex #8)
rabbit anti-dsRed(RFP) (1:500, , )
guinea pig anti-BiP(Hsc70-3) (1:1000, gift from Dr. Hermann Steller, (Ryoo et al., 2007))
mouse anti-HA 16B12 (1:1000, BioLegend, #901501)
rabbit anti-Cleaved Caspase-3 (1:100, Cell Signaling Technology, #9661)

*Secondary antibodies (all 1:250 dilution)*
Alexa Fluor 647 AffiniPure Donkey Anti-Guinea Pig (Jackson ImmunoResearch, 706-605-148)
Alexa Fluor 647 AffiniPure Goat Anti-Rat (Jackson ImmunoResearch, 112-605-167)
Alexa Fluor 488 AffiniPure Donkey Anti-Mouse (Jackson ImmunoResearch, 715-545-151)
Alexa Fluor 647 AffiniPure Donkey Anti-Mouse (Jackson ImmunoResearch, 715-605-150)
Cy3 AffiniPure Donkey Anti-Mouse (Jackson ImmunoResearch, 715-165-151)
Cy3 AffiniPure Donkey Anti-Rabbit (Jackson ImmunoResearch, 711-165-152)

**Antenna dissection and imaging**

For Gr21a:syt-RFP and Ldh-GFP antenna, endogenous RFP/GFP signals were imaged without antibody staining. The 2nd and 3rd segments of antenna were dissected together, mounted with SlowFade™ Gold Antifade Mountant, and imaged immediately with Leica STELLARIS 5 confocal microscope as Z series with the same interval. RFP+/GFP+ cell bodies were counted from whole Z planes. Representative images were merged by ImageJ (Image-Stacks-Z projection-Max Intensity). Quantification graphs are generated in GraphPad Prism. P-values were computed using the parametric unpaired t-test.

For auditory neuron nuclei detection, the 2nd and 3rd segments of antenna were dissected together, placed into 0.2ml PCR and fixed with 4% PFA in 3% PBST (1X PBS + 3% Triton-X100) for 20 min at RT. The antennae were thoroughly washed with 3% PBST and blocked with an antenna blocking buffer (5% normal goat serum (NGS) in 3% PBST) for 2 hours at RT. After blocking, the brains were incubated with mouse anti-Elav (1:40, DSHB, 9F8A9) in the antenna blocking buffer at 4ºC for 2 days. The antennae were thoroughly rinsed with 3% PBST and incubated with Alexa Fluor 488 AffiniPure Donkey Anti-Mouse (1:250, Jackson ImmunoResearch, 715-545-151) in the antenna blocking buffer at RT for 2 hours. The antennae were thoroughly rinsed with 3% PBST, incubated with DAPI stain (1:1000) for 20 min, washed with 1X PBS. mounted SlowFade™ Gold Antifade Mountant (Thermo Fisher, S36936). Images were obtained with Leica STELLARIS 5 confocal microscope as Z series with the same interval. Elav+ cell bodies were counted from whole Z planes. Representative images were merged by ImageJ (Image-Stacks-Z projection-Max Intensity). Quantification graphs were generated in GraphPad Prism. P-values were computed using the parametric unpaired t-test.

**Fat body dissection and LD staining**

For fat body dissection, fly abdomen filets were dissected in 1X PBS using Vannas Spring Scissors (FST, 3mm Cutting Edge, 15000-00) and forceps. The abdomen filets were placed into 1.5ml Eppendorf tube, fixed with 4% PFA for 20 min at RT, and washed with 1X PBS. Filets were incubated in Nile Red buffer (1:1,000 dilution of 1 mg/ml Nile Red (Sigma) in 1X PBS) for

10 min at RT, thoroughly rinsed with 1X PBS, and mounted with SlowFade™ Gold Antifade Mountant. Images were obtained with Leica STELLARIS 5 confocal microscope as Z series with the same interval. Z series images were merged by ImageJ (Image-Stacks-Z projection-Max Intensity), and then the size and number of LD was measured by the ImageJ plugin (see StarDist2D quantification).

**Gut dissection and immunofluorescence**

Fly guts (15-20 females) were dissected in cold 1X PBS, placed into a 24 well plate and fixed with 4% PFA in 1X PBS for 45 min at RT. The guts were thoroughly washed with 0.2% PBST (1X PBS + 0.2% Triton-X100) and blocked with a gut blocking buffer (5% normal goat serum (NGS) in 0.2% PBST) for one hour at 4ºC. After blocking, the guts were incubated with primary antibodies, rabbit anti-pH3 (1:1000, Cell Signaling Technology, #9701) and a combination cocktail of mouse anti-armadillo (1:100, DSHB, N2 7A1) and mouse anti-prospero (1:250, DSHB, MR1A), in the gut blocking buffer at 4ºC overnight. The guts were thoroughly rinsed with 0.2% PBST and incubated with secondary antibodies, Cy3 AffiniPure Donkey Anti-Rabbit (1:250, Jackson ImmunoResearch, 711-165-152) and Alexa Fluor 647 AffiniPure Donkey Anti-Mouse (Jackson ImmunoResearch, 715-605-150) in gut blocking buffer, at RT for 2 hours. The guts thoroughly rinsed with 0.2% PBST, and incubated with DAPI stain (1:1000) for 20 min at 4ºC. The guts were mounted SlowFade™ Gold Antifade Mountant. Images were obtained with Leica STELLARIS 5 confocal microscope as Z series with the same interval. Z series images were merged on half of the Z-stacks (merged = total z-stacks with the gut visible/2) by ImageJ (Image-Stacks-Z projection-Max Intensity), and the DAPI channel was extracted from the image for quantification (see StarDist2D quantification).

**StarDist2D quantification for LD and gut nuclei**

To measure the number and area of fat body LD or gut nuclei for each sample, the ImageJ (Fiji) plugin, StarDist2D, was implemented (Stevens et al., 2022). Nile Red or DAPI channel was extracted from the images. The StarDist2D plugin parameters were variable between images but the most accurate settings were applied on a per image basis. Images were manually curated to remove false positives and add false negatives. The tables of LD/nuclei number and area were produced by the ImageJ measure function. For LD quantification, mean LD size and >300$\mu m^2$ LD count were calculated from each sample using Excel. Quantification graphs were generated by GraphPad Prism. P-values were computed using the parametric unpaired t-test. For gut nuclei size quantification, frequency distribution analysis in GraphPad Prism was used (Parameters: Relative frequency percentages. Bin width 2$\mu m^2$, Bin each replicate, Bar graph).

**Male fecundity assay**

Individual 20d male fly (N>25 per genotypes, control or hTau flies) was crossed with 3 young wild-type female virgin flies. After mating in 29°C for 3 days, female flies were separated into individual vials (one female/vial) and incubated in 25°C. Fly vials were changed every 3 days. Total numbers of progenies from each cross were counted including the mating vial and divided by the number of female flies. Dead flies during mating were censored.

**Gut snRNA-seq trajectory analysis**

Trajectory analysis of the gut cells was performed using Scanpy. The intestinal stem cell lineage was extracted from the data, which include the intestinal stem cells, enteroblasts, adult differentiating enterocytes, enteroendocrine cells, anterior enterocytes of the adult midgut epithelium, and posterior enterocytes of the adult midgut epithelium. The pseudotime of the clusters was inferred using the partition-based graph abstraction (PAGA) function and then the ForceAtlas2 algorithm was implemented to spatially overlay the cells onto the PAGA plot. To compare the cell compositions of each genotype (control, Abeta42, and hTau), the genotypes were subset separately from the final pseudotime plot.

**Cell type annotation in AD-FCA**

Our approach to annotating cell types in AD-FCA closely mirrored the methodology previously established for AFCA annotations (Lu et al., 2023). We integrated AD-FCA samples with existing FCA and AFCA datasets, facilitating their co-clustering. To mitigate batch effects and align dataset variations, the Harmony algorithm (Korsunsky et al., 2019) was applied to the co-clustered data. Subsequent to adjustment, AFCA-derived cell type labels were assigned to AD-FCA cells using a Logistic Regression classifier, with AFCA serving as the training set and AD-FCA as the test set. These initial automated annotations were subsequently subjected to manual validation and correction to enhance reliability.

To enrich the diversity of cell types identified within our head samples, this annotation strategy was extended to incorporate two additional datasets: FCA antenna data and the optical lobe data (Li et al., 2022; Özel et al., 2021). Through this expanded analysis, we successfully annotated a total of 219 distinct cell types.

**DEG analysis**

Differential gene expression analysis for identification of genes with altered expression levels in Aβ42 and hTau flies in comparison to age-matched controls was conducted using the Wilcoxon Rank Sum test. Genes were considered differentially expressed if they exhibited a FDR of less than 0.05. Cell types with more than 700 nuclei were selected for our DEG analyses.

**Analysis of cell composition changes**

Quantitative assessments of cell composition changes involved counting cell numbers for each cell type across different ages and genotypes. To calculate the proportion of a given cell type within a specific genotype, the cell count for that cell type and genotype was normalized to the total cell number of the same genotype. This normalization facilitated the comparison of cell type proportions between experimental (Aβ42 or hTau flies) and age-matched control groups. The resulting ratios for the experimental groups were then divided by those of the control groups, with the differences expressed in log2 scale to quantify relative changes in cell composition. Cell types with more than 700 nuclei were selected for our cell composition analyses.

**Gene ontology and pathway enrichment analysis**

Differential expression analysis yielded genotype specific DEGs, encompassing both upregulated and downregulated genes across various cell types. These DEGs were subjected to Gene Ontology (GO) analysis using the GOATOOLS software (v1.2.3) (Klopfenstein et al., 2018). For this purpose, the gene association dataset (FB2022_04) was retrieved from FlyBase, with a specific focus on Biological Process (BP) GO terms for our investigations (refer to fig. S11).

Pathway enrichment analysis leveraged the Kyoto Encyclopedia of Genes and Genomes (KEGG) database, with *Drosophila*-specific pathways selected to suit our study's requirements (Kanehisa and Goto, 2000). The integration of DEGs with these selected pathways was facilitated through the GSEApy package (Fang et al., 2023), enabling the identification of significantly enriched pathways.

**Correlation analysis of UMI numbers and nuclear ratios**

To evaluate the correlation of UMI numbers across cell types, we initially determined the median UMI counts for each cell type within individual samples. These median UMI counts were then compared across various samples to calculate ratios that reflect differences attributable to genotype or age. We utilized Pearson's or Spearman's correlation coefficients to identify groups exhibiting similar trends in UMI number alterations. Furthermore, we extended our correlation analysis to encompass nuclear ratios. These ratios were derived by comparing nuclear counts between pairs of samples, as outlined in the "Analysis of Cell Composition Changes" section. Subsequently, we compared these nuclear ratios across different groups to discern patterns and trends consistent among various comparisons.

**Ranking of differentially expressed genes in hTau neurons**

In our analysis to delineate global gene expression alterations in hTau neurons relative to control neurons, we categorized each DEG and quantified the number of cell types wherein each gene exhibited differential expression. Remarkably, we observed that 97.6% of DEGs manifested differential expression across fewer than 10 cell types. Genes that were differentially expressed across more than 20 cell types were featured in Figure 6A.

**Analysis of cell-cell communication (CCC)**

Our investigation into CCC was guided by the foundational principles delineated in FlyPhoneDB (Liu et al., 2022), with specific adaptations. Initially, we extracted a collection of genes linked to the GO term "homophilic cell adhesion via plasma membrane adhesion molecules" from FlyBase. This was followed by a manual curation process, resulting in the selection of 44 genes that were subsequently incorporated into the existing ligand-receptor list within the FlyPhoneDB framework (Table S1).

For the analysis, a subset of 102 cell types—comprising 54 neuronal and 48 peripheral cell types, all characterized with more than 700 nuclei—was selected for inclusion in the FlyPhoneDB framework. This facilitated the computation of CCC metrics within each genotype under investigation. Only CCCs exhibiting a p-value less than 0.05 in control or hTau samples were retained for subsequent analyses, ensuring statistical relevance.

Particular emphasis was placed on CCCs emanating from neurons and targeting peripheral cells, enabling an exploration of the communicative alterations induced by neuronal hTau expression. To quantitatively assess these changes, we computed the 'Delta scores' by subtracting control CCC scores from those of the hTau samples. These Delta scores were then standardized to Z scores, based on their corresponding standard deviations. CCCs with absolute Z scores exceeding 2 were classified as differential CCCs, denoting statistically significant communication differences between hTau and control cohorts.

**AD-FCA data portal website**

The AD-FCA web platform was constructed utilizing the Shiny package (version 1.6.0) within the R programming environment (version 4.0.5), facilitating interactive web application development. The platform hosts two principal datasets: the Head dataset and the Body dataset, which encompass snRNA-Seq analytical outcomes derived from head and body tissues, respectively.

For the 'Cell Type' and 'Gene Expression' interface components, data handling and visual representation were executed employing the Tidyverse package (version 1.3.0) and the ggplot2 package (version 3.3.3) in R, ensuring a robust and efficient data processing pipeline. Additionally, the 'Custom Analysis' section of the platform is enriched by the functionalities provided by the ShinyCell package (version 2.1.0), offering users an enhanced and tailored analytical experience.

fig. S1

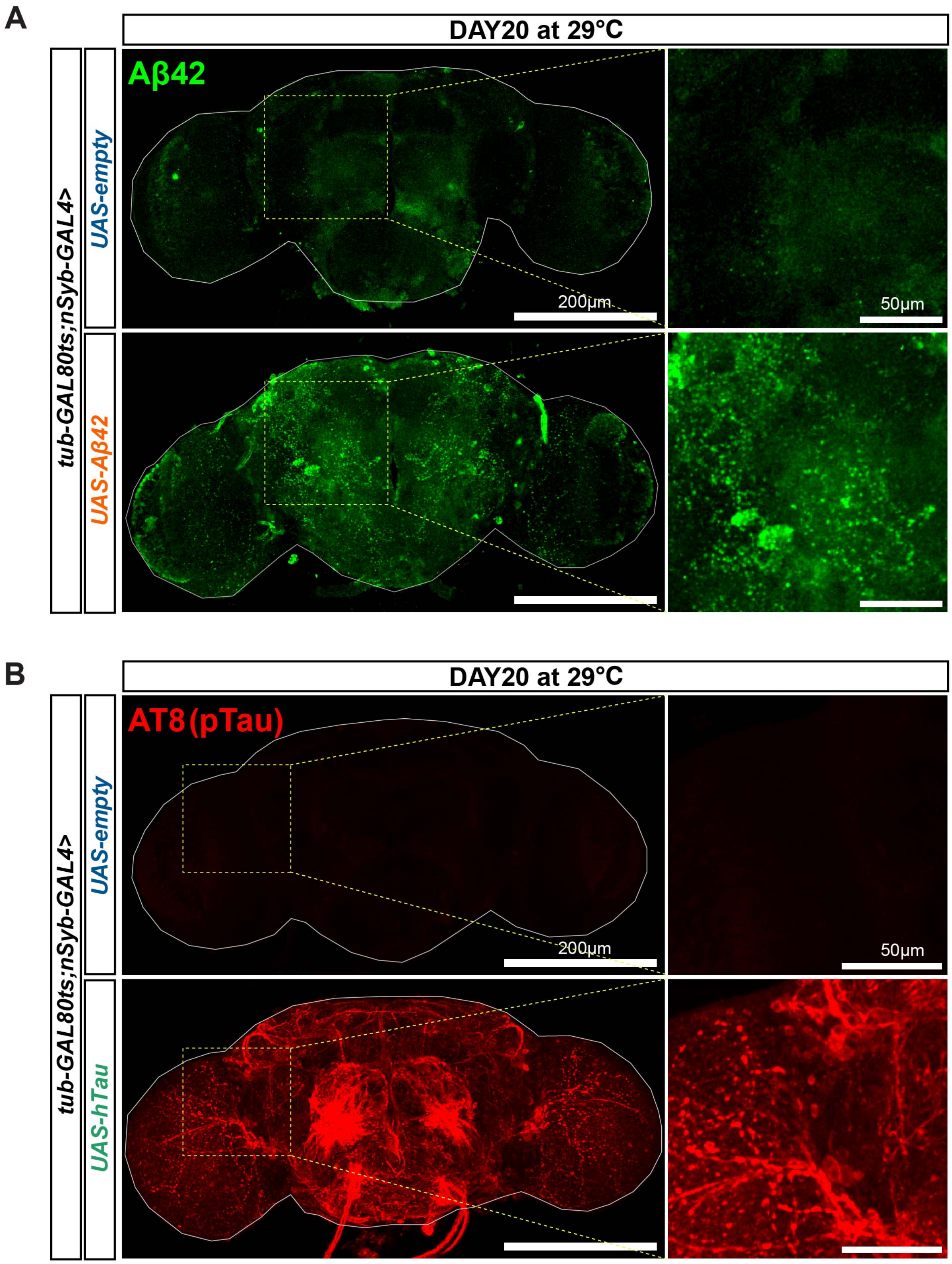
A
DAY20 at 29°C
tub-GAL80ts;nSyb-GAL4>
UAS-empty
UAS-Aβ42
Aβ42
200μm
50μm
B
DAY20 at 29°C
tub-GAL80ts;nSyb-GAL4>
UAS-empty
UAS-hTau
AT8(pTau)
200μm
50μm

**fig. S1. Aβ42 and hTau expression in the brain of 20d AD fly models**

(A) anti-Aβ42 antibody staining of the whole brain in control and Aβ42 flies.

(B) anti-phospho-Tau (AT8) antibody staining of the whole brain in control and hTau flies.

### fig. S2

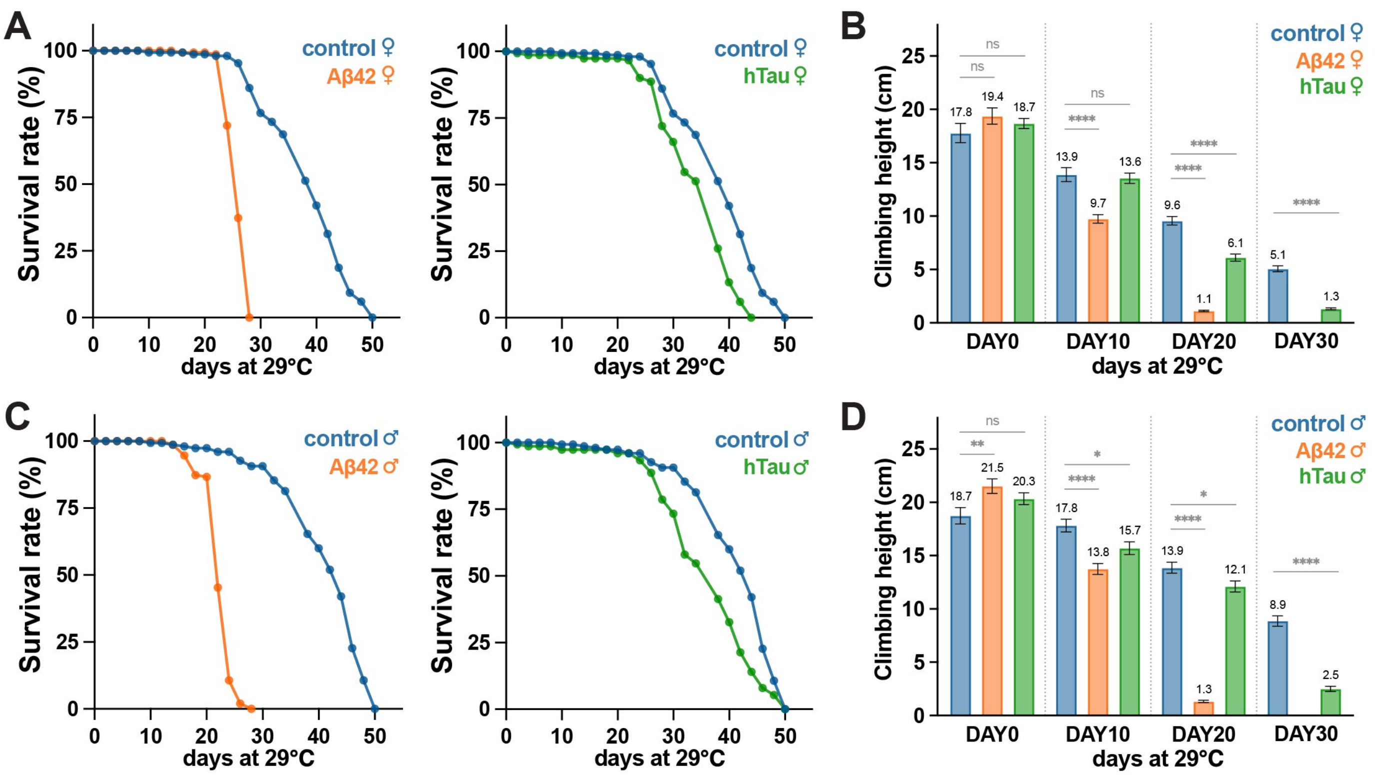

A
B
C
D
Survival rate (%)
days at 29°C
control ♀
Aβ42 ♀
hTau ♀
control ♂
Aβ42 ♂
hTau ♂
Climbing height (cm)
DAY0
DAY10
DAY20
DAY30
ns
**
****
*
17.8
19.4
18.7
13.9
9.7
13.6
9.6
1.1
6.1
5.1
1.3
18.7
21.5
20.3
17.8
13.8
15.7
13.9
1.3
12.1
8.9
2.5

**fig. S2. Separated survival curves and negative geotaxis plots.**

(A) Survival curves of female flies at 29°C. N=150 per genotype. (B) Negative geotaxis plots of female flies at 29°C. 20 flies per genotype per set. Flies climbed 10 times per set for 10 seconds. 2 sets of 0d (N=40 per genotype), 4 sets of 10d (N=80 per genotype), 5 sets of 20d (N=100 per genotype), 5 sets of 30d (N=100 per genotype). Bars represent mean ± SEM. **** $p < 0.0001$. The numbers above the bars indicate mean values.

(C) Survival curves of male flies at 29°C. N=150 per genotype.

(D) Negative geotaxis plots of male flies at 29°C. 20 flies per genotype per set. Flies climbed 10 times per set for 10 seconds. 3 sets of 0d (N=40 per genotype), 4 sets of 10d (N=80 per genotype), 5 sets of 20d (N=100 per genotype), 5 sets of 30d (N=100 per genotype). Bars represent mean ± SEM. **** $p < 0.0001$. The numbers above the bars indicate mean values.

P-values were computed using the parametric unpaired t-test.

fig. S3

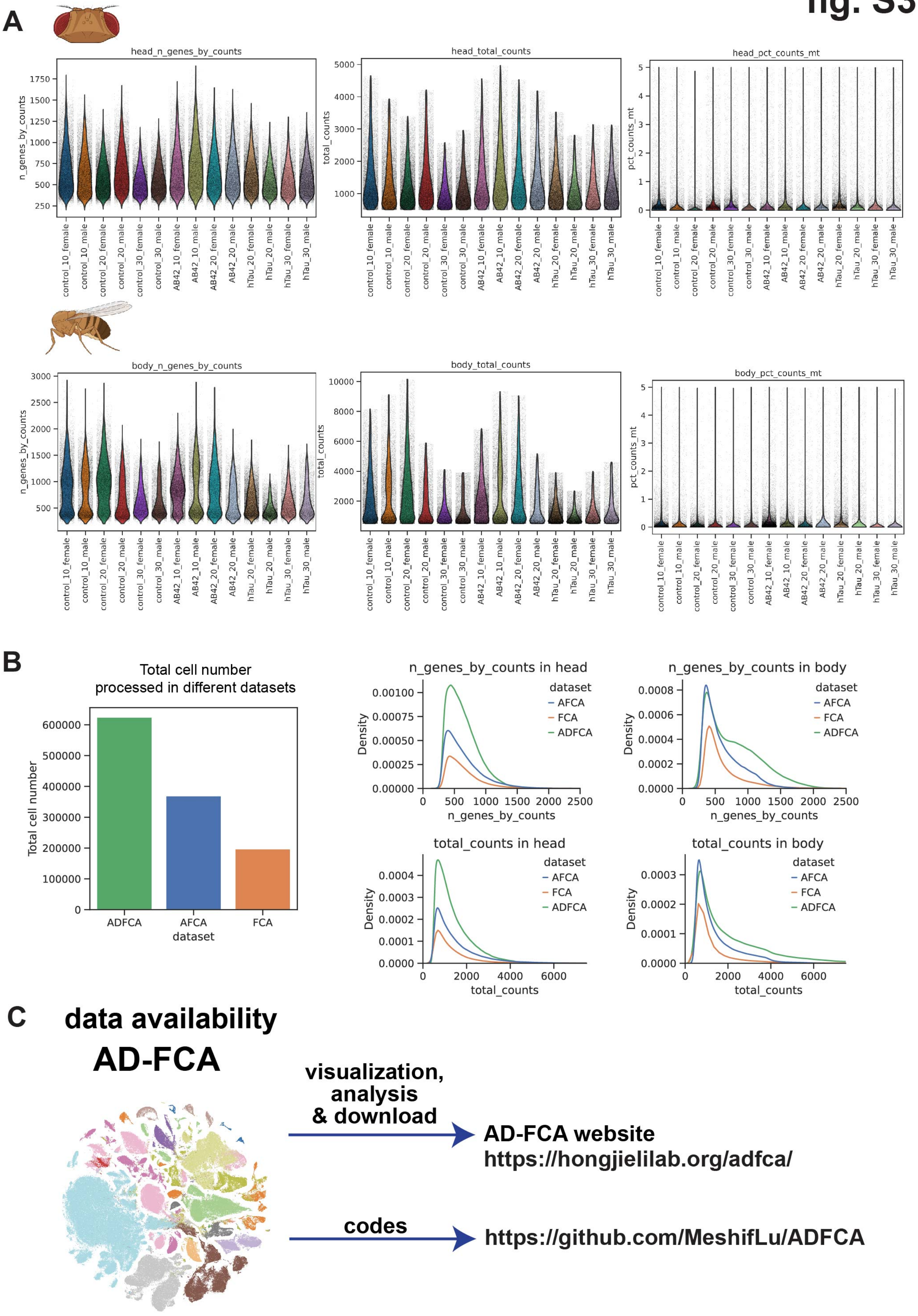

A
head_n_genes_by_counts
n_genes_by_counts
head_total_counts
total_counts
head_pct_counts_mt
pct_counts_mt
body_n_genes_by_counts
body_total_counts
body_pct_counts_mt
control_10_female
control_10_male
control_20_female
control_20_male
control_30_female
control_30_male
AB42_10_female
AB42_10_male
AB42_20_female
AB42_20_male
hTau_20_female
hTau_20_male
hTau_30_female
hTau_30_male
B
Total cell number processed in different datasets
Total cell number
dataset
ADFCA
AFCA
FCA
n_genes_by_counts in head
n_genes_by_counts in body
total_counts in head
total_counts in body
Density
n_genes_by_counts
total_counts
C
data availability
AD-FCA
visualization, analysis & download
AD-FCA website
https://hongjielilab.org/adfca/
codes
https://github.com/MeshifLu/ADFCA

**fig. S3. Data quality of snRNA-seq results and data availability.**

(A) Expressed gene numbers, UMI numbers, and mitochondria transcript ratios of each sample from the AD-FCA data.

(B) High-throughput kit improved both the nuclear yield and the number of detected genes/transcripts in AD-FCA compared to AFCA and FCA.

(C) Data available from AD-FCA website (visualization, analysis and download) and GitHub (codes).

**fig. S4**

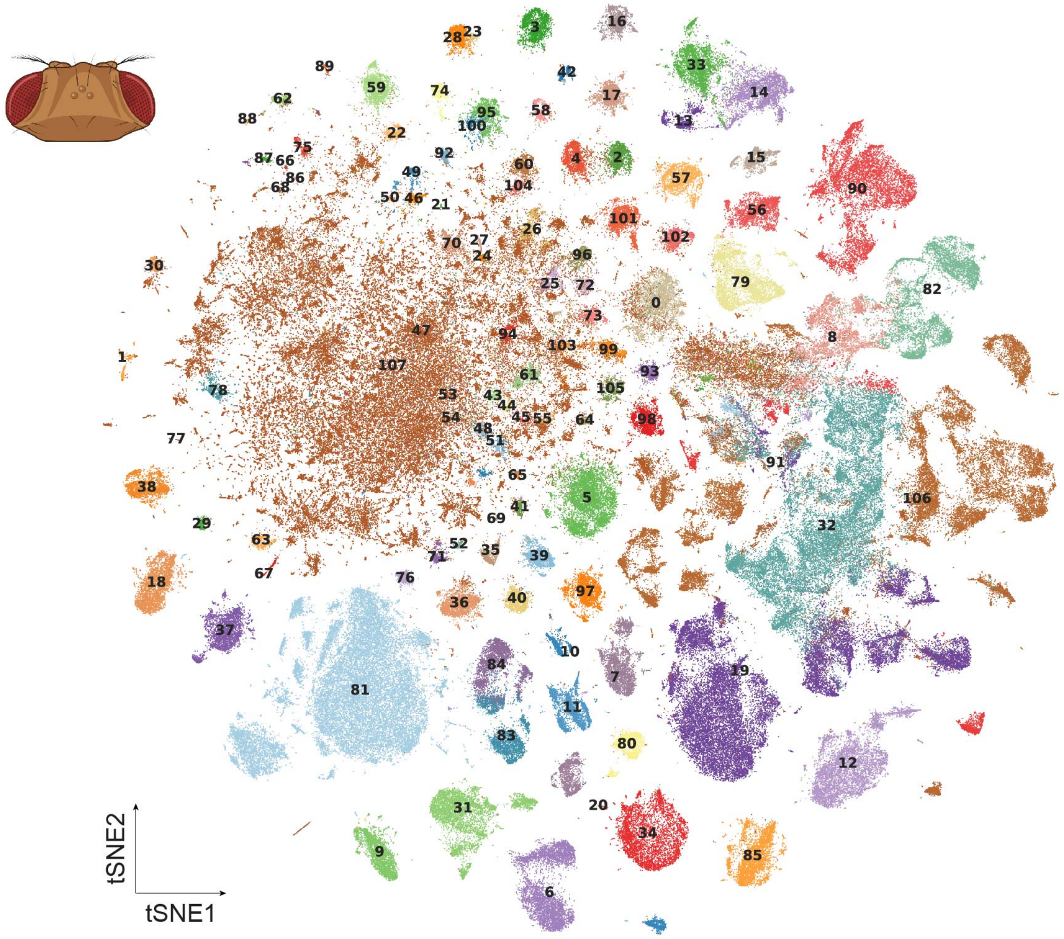

tSNE2
tSNE1



0.ABeta42-specific cluster
1.Poxn neuron
2.T neuron T2
3.T neuron T2a
4.T neuron T3
5.T neuron T4/T5
6.adult brain cell body glial cell
7.adult brain perineurial glial cell
8.adult fat body_head
9.adult lamina epithelial/marginal glial cell
10.adult optic chiasma glial cell
11.adult reticular neuropil associated glial cell_head
12.adult tracheal cell_head
13.alpha'/beta' Kenyon cell
14.alpha/beta Kenyon cell
15.auditory sensory neuron
16.centrifugal neuron C2
17.centrifugal neuron C3
18.columnar neuron T1
19.cone cell
20.crystal cell_head
21.distal medullary amacrine neuron Dm1
22.distal medullary amacrine neuron Dm10
23.distal medullary amacrine neuron Dm11
24.distal medullary amacrine neuron Dm12
25.distal medullary amacrine neuron Dm2
26.distal medullary amacrine neuron Dm3
27.distal medullary amacrine neuron Dm4
28.distal medullary amacrine neuron Dm8
29.distal medullary amacrine neuron Dm9
30.dopaminergic PAM neuron
31.ensheathing glial cell
32.epithelial cell_head
33.gamma Kenyon cell
34.hemocyte_head
35.lamina intrinsic neuron
36.lamina monopolar neuron L1
37.lamina monopolar neuron L2
38.lamina monopolar neuron L3
39.lamina monopolar neuron L4
40.lamina monopolar neuron L5
41.lamina wide-field 1 neuron
42.lamina wide-field 2 neuron
43.lobula columnar neuron LC10a
44.lobula columnar neuron LC10b
45.lobula columnar neuron LC10d
46.lobula columnar neuron LC12
47.lobula columnar neuron LC14
48.lobula columnar neuron LC16
49.lobula columnar neuron LC17
50.lobula columnar neuron LC4
51.lobula columnar neuron LC6
52.lobula plate columnar LPC1
53.lobula plate-lobula columnar LPLC1
54.lobula plate-lobula columnar LPLC2
55.lobula-lobula plate columnar LLPC1
56.mechanosensory neuron
57.medullary intrinsic neuron Mi1
58.medullary intrinsic neuron Mi15
59.medullary intrinsic neuron Mi4
60.medullary intrinsic neuron Mi9
61.neuron 1
62.neuron 10
63.neuron 11
64.neuron 12
65.neuron 13
66.neuron 14
67.neuron 15
68.neuron 16
69.neuron 17
70.neuron 2
71.neuron 4
72.neuron 5
73.neuron 6
74.neuron 8
75.neuron 9
76.ocellus retinula cell
77.octopaminergic/tyraminergic neuron
78.olfactory projection neuron
79.olfactory receptor neuron
80.optic-lobe-associated cortex glial cell
81.outer photoreceptor cell
82.pericerebral adult fat mass
83.photoreceptor cell R7
84.photoreceptor cell R8
85.pigment cell
86.proximal medullary amacrine neuron Pm1
87.proximal medullary amacrine neuron Pm2
88.proximal medullary amacrine neuron Pm3*
89.proximal medullary amacrine neuron Pm4
90.skeletal muscle of head
91.subperineurial glial cell_head
92.transmedullary Y neuron TmY14
93.transmedullary Y neuron TmY3
94.transmedullary Y neuron TmY4
95.transmedullary Y neuron TmY5a
96.transmedullary Y neuron TmY8
97.transmedullary neuron Tm1
98.transmedullary neuron Tm2
99.transmedullary neuron Tm20
100.transmedullary neuron Tm29
101.transmedullary neuron Tm3a
102.transmedullary neuron Tm4
103.transmedullary neuron Tm5ab
104.transmedullary neuron Tm5c
105.transmedullary neuron Tm9
106.unannotated
107.uncharacterized CNS neuron

**fig. S4. tSNE of detailed cell types annotated in the head sample.**
107 head cell types are shown in the tSNE plot.

### fig. S5

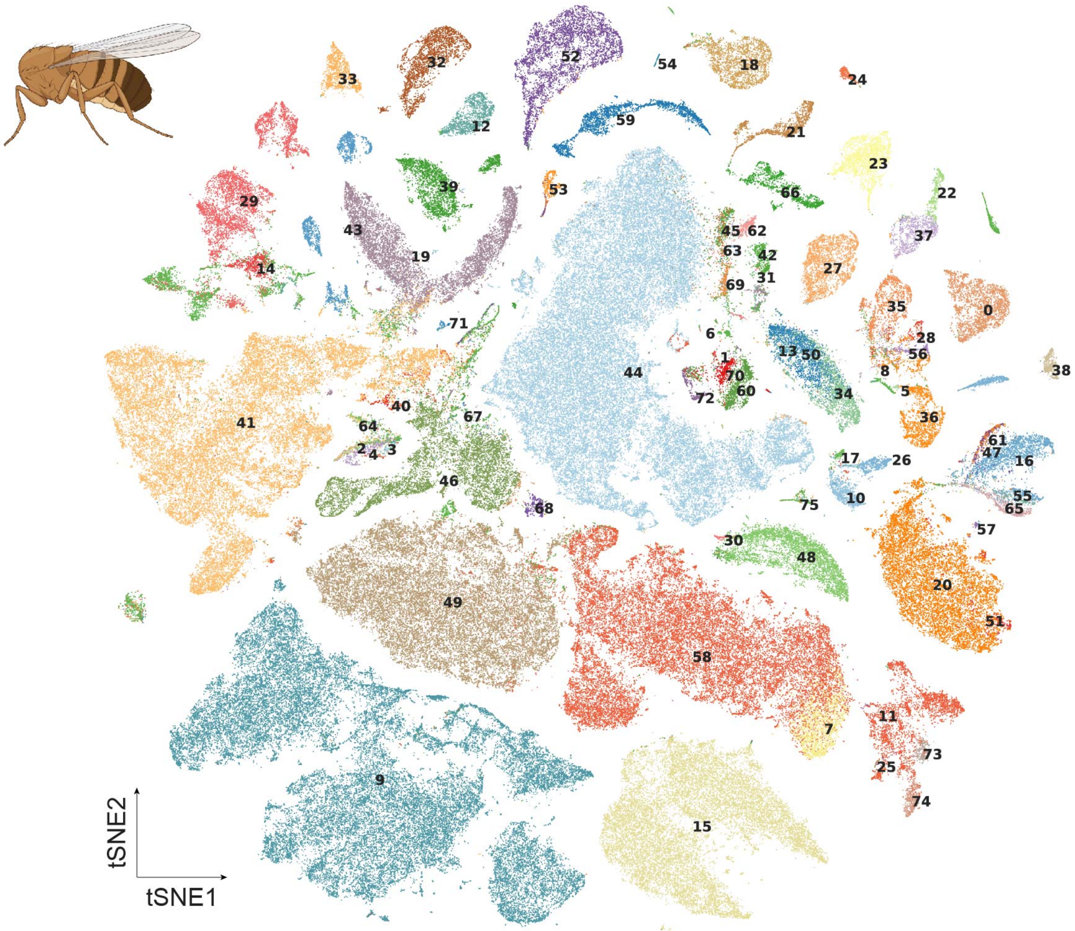


- 0.16-cell germline cyst in germarium region 2a and 2b
- 1.CNS surface associated glial cell
- 2.adult Malpighian tubule principal cell
- 3.adult Malpighian tubule principal cell of initial segment
- 4.adult Malpighian tubule principal cell of lower segment
- 5.adult Malpighian tubule principal cell of lower ureter
- 6.adult Malpighian tubule stellate cell of main segment
- 7.adult alary muscle
- 8.adult differentiating enterocyte
- 9.adult fat body_body
- 10.adult glial cell
- 11.adult heart ventral longitudinal muscle
- 12.adult hindgut
- 13.adult midgut enterocyte
- 14.adult midgut-hindgut hybrid zone
- 15.adult oenocyte
- 16.adult peripheral nervous system
- 17.adult reticular neuropil associated glial cell_body
- 18.adult salivary gland
- 19.adult tracheal cell_body
- 20.adult ventral nervous system
- 21.anterior ejaculatory duct
- 22.antimicrobial peptide-producing cell
- 23.cardia (1)
- 24.cardia (2)
- 25.cardiomyocyte, working adult heart muscle (non-ostia)
- 26.cell body glial cell
- 27.choriogenic main body follicle cell and corpus luteum
- 28.copper cell
- 29.crop
- 30.crystal cell_body
- 31.cyst cell
- 32.ejaculatory bulb
- 33.ejaculatory bulb epithelium
- 34.enteroblast
- 35.enterocyte of anterior adult midgut epithelium
- 36.enterocyte of posterior adult midgut epithelium
- 37.enterocyte-like
- 38.enteroendocrine cell
- 39.eo support cell
- 40.epidermal cell that specialized in antimicrobial response
- 41.epithelial cell_body
- 42.escort cell
- 43.female reproductive system
- 44.follicle cell
- 45.follicle stem cell and prefollicle cell
- 46.germline cell
- 47.gustatory receptor neuron
- 48.hemocyte_body
- 49.indirect flight muscle
- 50.intestinal stem cell
- 51.leg muscle motor neuron
- 52.male accessory gland main cell_roX1+
- 53.male accessory gland main cell_roX1-
- 54.male accessory gland secondary cell
- 55.mechanosensory neuron of haltere
- 56.midgut large flat cell
- 57.multidendritic neuron
- 58.muscle cell
- 59.oviduct
- 60.perineurial glial sheath
- 61.pheromone-sensing neuron
- 62.polar follicle cell
- 63.prefollicle cell/stalk follicle cell
- 64.principal cell*
- 65.scolopidial neuron
- 66.seminal vesicle & testis epithelia
- 67.spermatid
- 68.spermatocyte
- 69.stalk follicle cell
- 70.subperineurial glial cell_body
- 71.unannotated
- 72.uncharacterized glial cell
- 73.visceral muscle of the crop
- 74.visceral muscle of the midgut
- 75.young germ cell

**fig. S5. tSNE of detailed cell types annotated in the body sample.**
75 body cell types are shown in the tSNE plot.

fig. S6

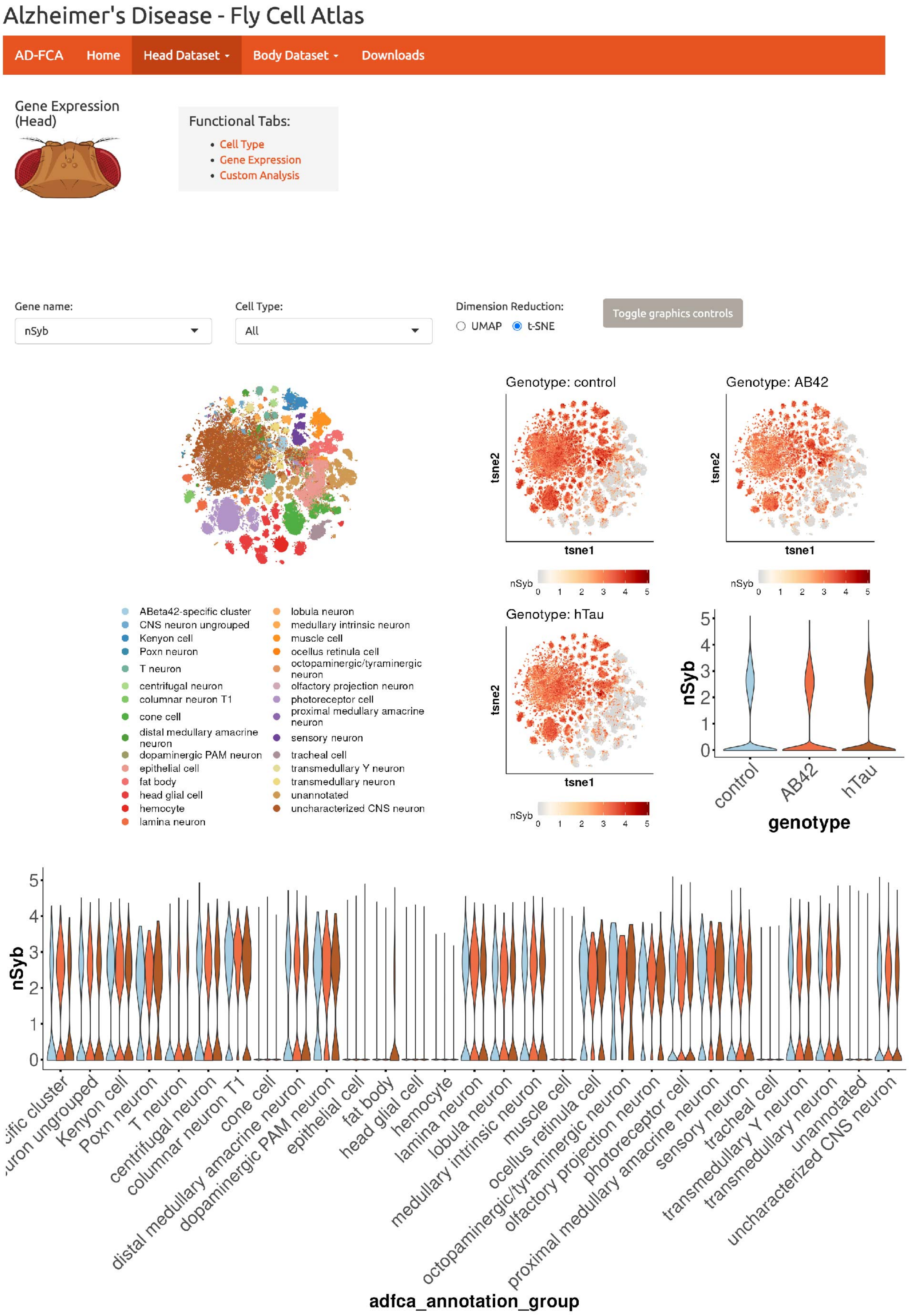

Alzheimer's Disease - Fly Cell Atlas
AD-FCA
Home
Head Dataset
Body Dataset
Downloads
Gene Expression (Head)
Functional Tabs:
Cell Type
Gene Expression
Custom Analysis
Gene name:
nSyb
Cell Type:
All
Dimension Reduction:
UMAP
t-SNE
Toggle graphics controls
Genotype: control
Genotype: AB42
Genotype: hTau
tsne1
tsne2
nSyb
ABeta42-specific cluster
CNS neuron ungrouped
Kenyon cell
Poxn neuron
T neuron
centrifugal neuron
columnar neuron T1
cone cell
distal medullary amacrine neuron
dopaminergic PAM neuron
epithelial cell
fat body
head glial cell
hemocyte
lamina neuron
lobula neuron
medullary intrinsic neuron
muscle cell
ocellus retinula cell
octopaminergic/tyraminergic neuron
olfactory projection neuron
photoreceptor cell
proximal medullary amacrine neuron
sensory neuron
tracheal cell
transmedullary Y neuron
transmedullary neuron
unannotated
uncharacterized CNS neuron
control
AB42
hTau
genotype
adfca_annotation_group

**fig. S6. AD-FCA data portal for surveying the AD-related changes in the head dataset.** AD-FCA web portal provides cell type-based, gene-based, and custom analyses for the head dataset.

fig. S7

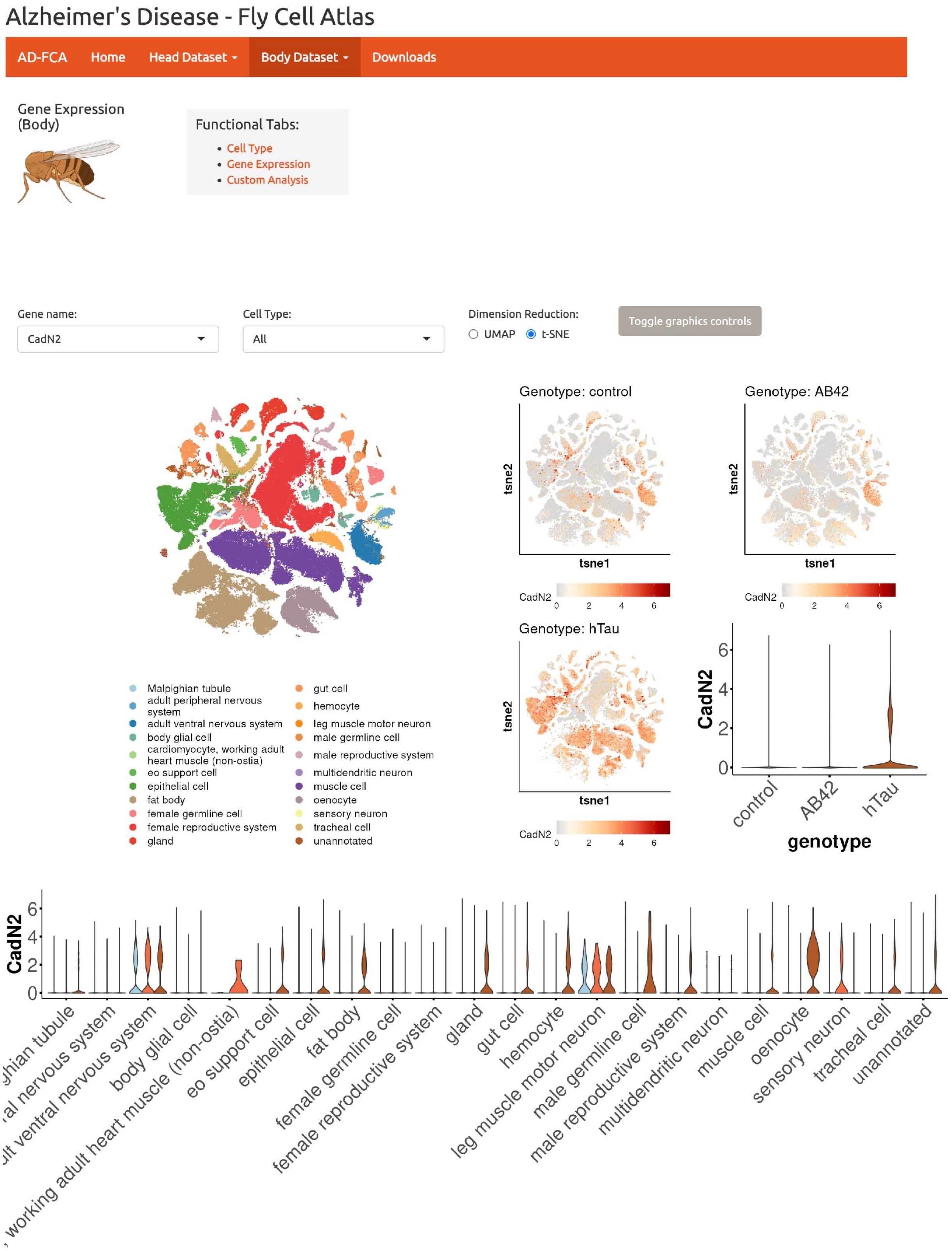

Alzheimer's Disease - Fly Cell Atlas
AD-FCA
Home
Head Dataset
Body Dataset
Downloads
Gene Expression (Body)
Functional Tabs:
Cell Type
Gene Expression
Custom Analysis
Gene name:
CadN2
Cell Type:
All
Dimension Reduction:
UMAP
t-SNE
Toggle graphics controls
Genotype: control
Genotype: AB42
Genotype: hTau
tsne1
tsne2
CadN2
Malpighian tubule
adult peripheral nervous system
adult ventral nervous system
body glial cell
cardiomyocyte, working adult heart muscle (non-ostia)
eo support cell
epithelial cell
fat body
female germline cell
female reproductive system
gland
gut cell
hemocyte
leg muscle motor neuron
male germline cell
male reproductive system
multidendritic neuron
muscle cell
oenocyte
sensory neuron
tracheal cell
unannotated
control
AB42
hTau
genotype

**fig. S7. AD-FCA data portal for surveying the AD-related changes in the head dataset.** AD-FCA web portal provides cell type-based, gene-based, and custom analyses for the body dataset.

**fig. S8**

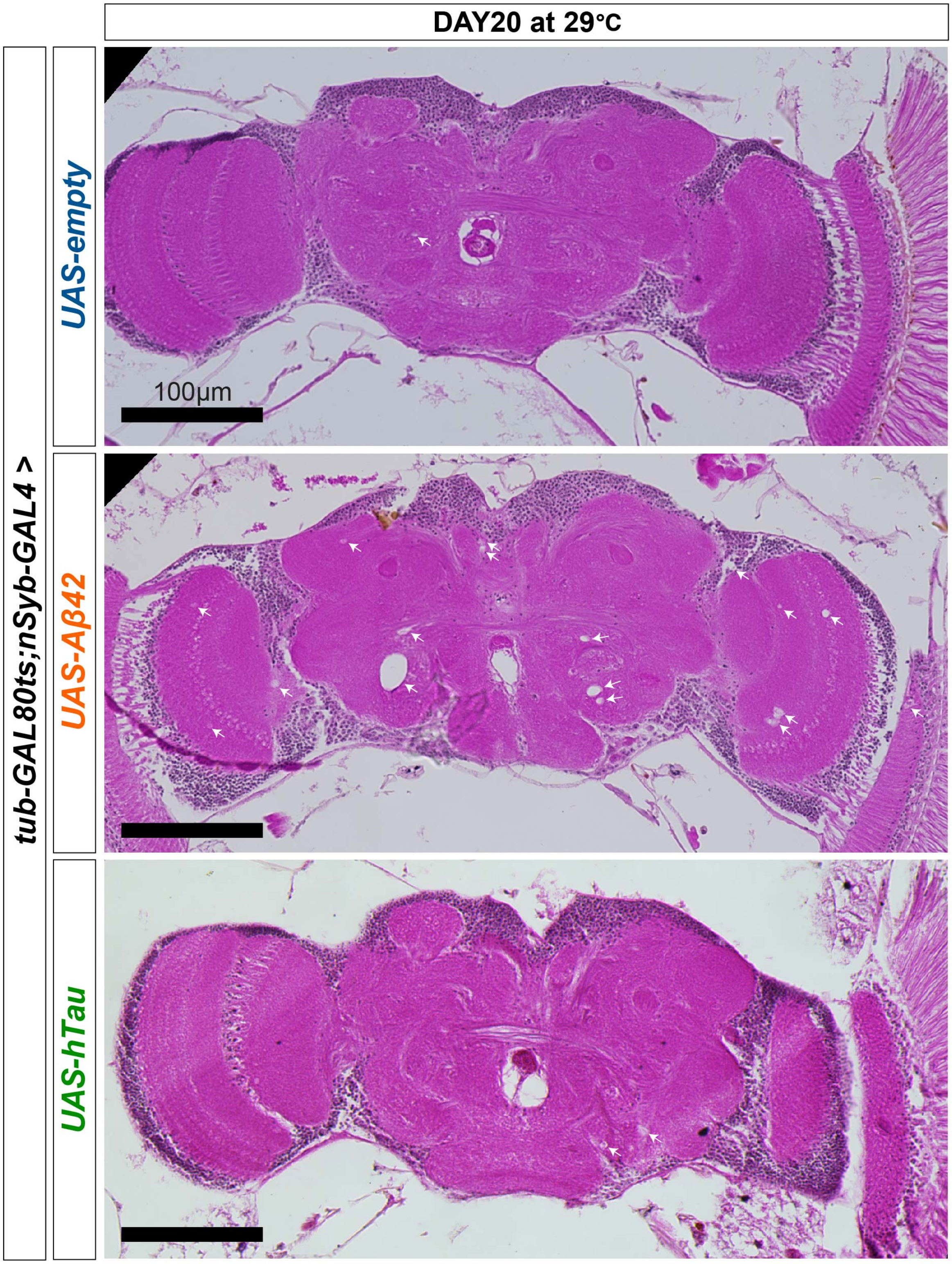

DAY20 at 29°C
tub-GAL80ts;nSyb-GAL4 >
UAS-empty
UAS-Aβ42
UAS-hTau
100µm

**fig. S8. Representative images of head sections showing vacuoles in the 20d fly brains**
Aβ42 fly brains show more dramatic vacuolization (white arrows) than hTau flies.

### fig. S9

A

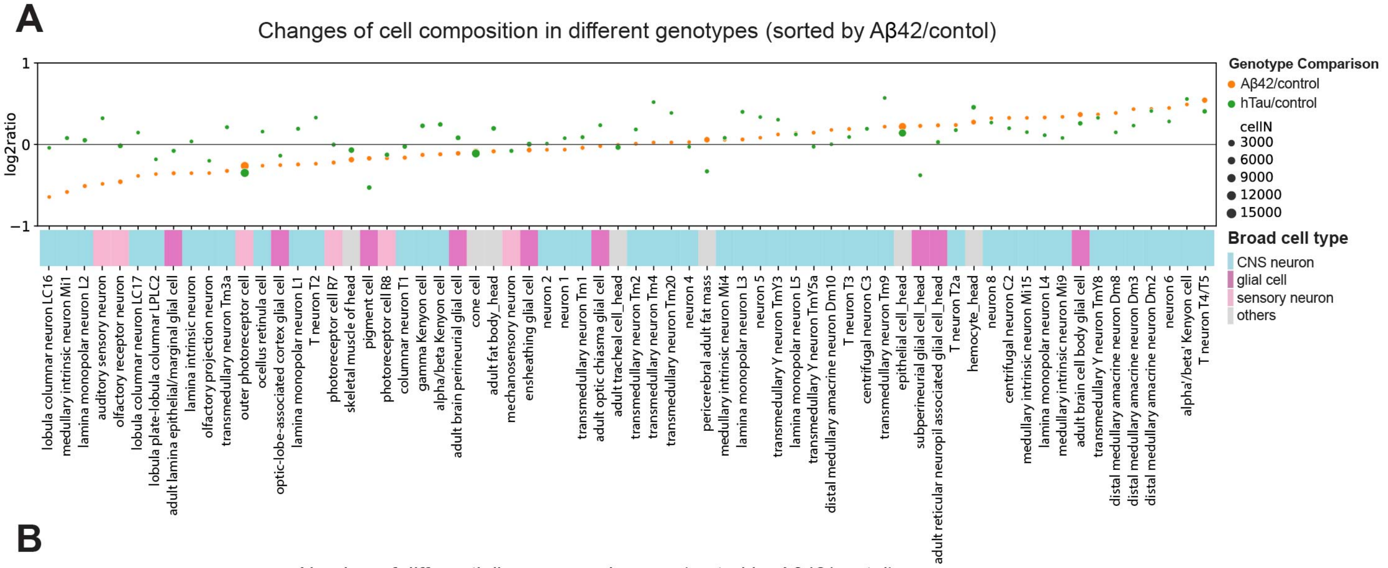
Changes of cell composition in different genotypes (sorted by Aβ42/contol)
log2ratio
1
0
−1
Genotype Comparison
Aβ42/control
hTau/control
cellN
3000
6000
9000
12000
15000
Broad cell type
CNS neuron
glial cell
sensory neuron
others
lobula columnar neuron LC16
medullary intrinsic neuron Mi1
lamina monopolar neuron L2
auditory sensory neuron
olfactory receptor neuron
lobula columnar neuron LC17
lobula plate-lobula columnar LPLC2
adult lamina epithelial/marginal glial cell
lamina intrinsic neuron
olfactory projection neuron
transmedullary neuron Tm3a
outer photoreceptor cell
ocellus retinula cell
optic-lobe-associated cortex glial cell
lamina monopolar neuron L1
T neuron T2
photoreceptor cell R7
skeletal muscle of head
pigment cell
photoreceptor cell R8
columnar neuron T1
gamma Kenyon cell
alpha/beta Kenyon cell
adult brain perineurial glial cell
cone cell
adult fat body_head
mechanosensory neuron
ensheathing glial cell
neuron 2
neuron 1
transmedullary neuron Tm1
adult optic chiasma glial cell
adult tracheal cell_head
transmedullary neuron Tm2
transmedullary neuron Tm4
transmedullary neuron Tm20
neuron 4
pericerebral adult fat mass
medullary intrinsic neuron Mi4
lamina monopolar neuron L3
neuron 5
transmedullary Y neuron TmY3
lamina monopolar neuron L5
transmedullary Y neuron TmY5a
distal medullary amacrine neuron Dm10
T neuron T3
centrifugal neuron C3
transmedullary neuron Tm9
epithelial cell_head
subperineurial glial cell_head
adult reticular neuropil associated glial cell_head
T neuron T2a
hemocyte_head
neuron 8
centrifugal neuron C2
medullary intrinsic neuron Mi15
lamina monopolar neuron L4
medullary intrinsic neuron Mi9
adult brain cell body glial cell
transmedullary Y neuron TmY8
distal medullary amacrine neuron Dm8
distal medullary amacrine neuron Dm3
distal medullary amacrine neuron Dm2
neuron 6
alpha'/beta' Kenyon cell
T neuron T4/T5

B

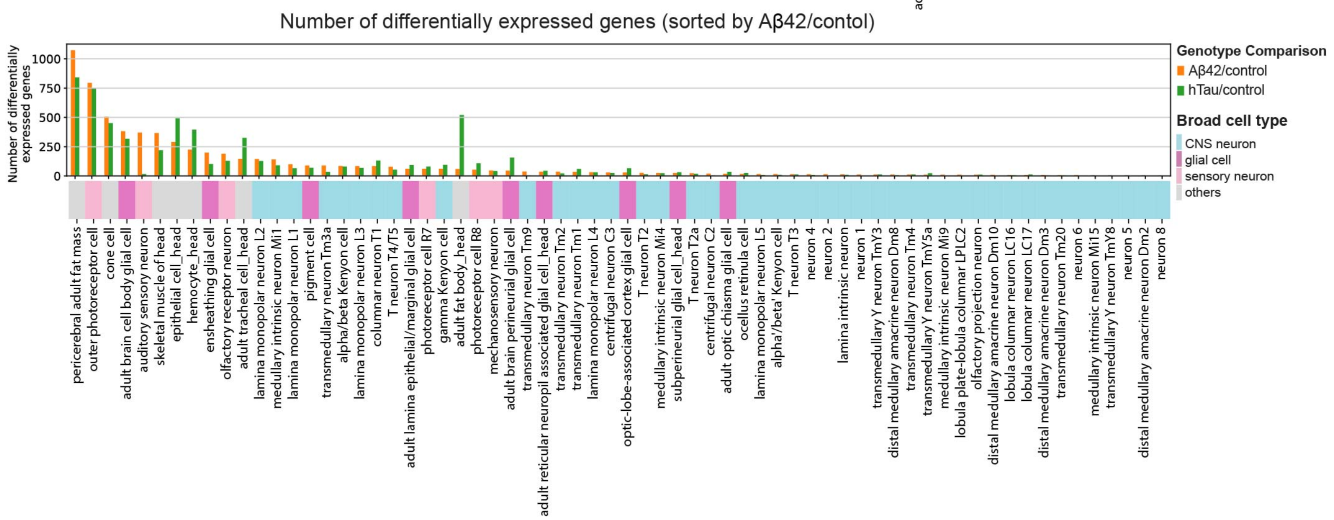
Number of differentially expressed genes (sorted by Aβ42/contol)
Number of differentially expressed genes
1000
750
500
250
0
Genotype Comparison
Aβ42/control
hTau/control
Broad cell type
CNS neuron
glial cell
sensory neuron
others
pericerebral adult fat mass
outer photoreceptor cell
cone cell
adult brain cell body glial cell
auditory sensory neuron
skeletal muscle of head
epithelial cell_head
hemocyte_head
ensheathing glial cell
olfactory receptor neuron
adult tracheal cell_head
lamina monopolar neuron L2
medullary intrinsic neuron Mi1
lamina monopolar neuron L1
pigment cell
transmedullary neuron Tm3a
alpha/beta Kenyon cell
lamina monopolar neuron L3
columnar neuron T1
T neuron T4/T5
adult lamina epithelial/marginal glial cell
photoreceptor cell R7
gamma Kenyon cell
adult fat body_head
photoreceptor cell R8
mechanosensory neuron
adult brain perineurial glial cell
transmedullary neuron Tm9
adult reticular neuropil associated glial cell_head
transmedullary neuron Tm2
transmedullary neuron Tm1
lamina monopolar neuron L4
centrifugal neuron C3
optic-lobe-associated cortex glial cell
T neuron T2
medullary intrinsic neuron Mi4
subperineurial glial cell_head
T neuron T2a
centrifugal neuron C2
adult optic chiasma glial cell
ocellus retinula cell
lamina monopolar neuron L5
alpha'/beta' Kenyon cell
T neuron T3
neuron 4
neuron 2
lamina intrinsic neuron
neuron 1
transmedullary Y neuron TmY3
distal medullary amacrine neuron Dm8
transmedullary neuron Tm4
transmedullary Y neuron TmY5a
medullary intrinsic neuron Mi9
lobula plate-lobula columnar LPLC2
olfactory projection neuron
distal medullary amacrine neuron Dm10
lobula columnar neuron LC16
lobula columnar neuron LC17
distal medullary amacrine neuron Dm3
transmedullary neuron Tm20
neuron 6
medullary intrinsic neuron Mi15
transmedullary Y neuron TmY8
neuron 5
distal medullary amacrine neuron Dm2
neuron 8

C

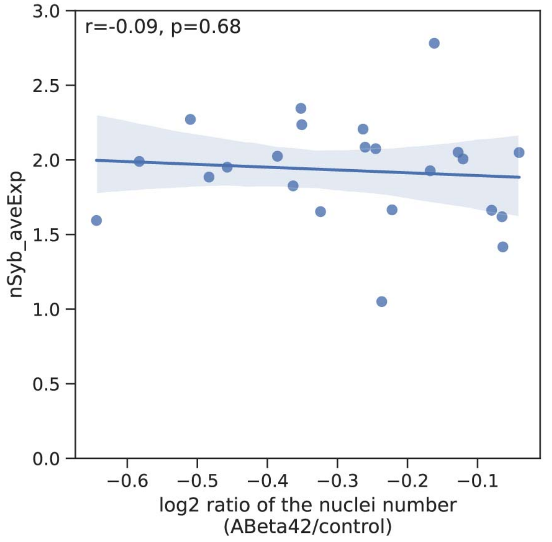
r=-0.09, p=0.68
nSyb_aveExp
3.0
2.5
2.0
1.5
1.0
0.5
0.0
−0.6
−0.5
−0.4
−0.3
−0.2
−0.1
log2 ratio of the nuclei number
(ABeta42/control)

**fig. S9. Analysis of cell composition and DEGs in head cell types**

(A) Cell composition changes in Aβ42 and hTau head cell types compared to controls.

(B) DEGs in Aβ42 and hTau head cell types versus controls.

(C) Pearson's correlation between nSyb expression levels and nuclear ratio changes in neuronal cells.

**fig. S10**

#### olfactory receptor neuron (ORN)

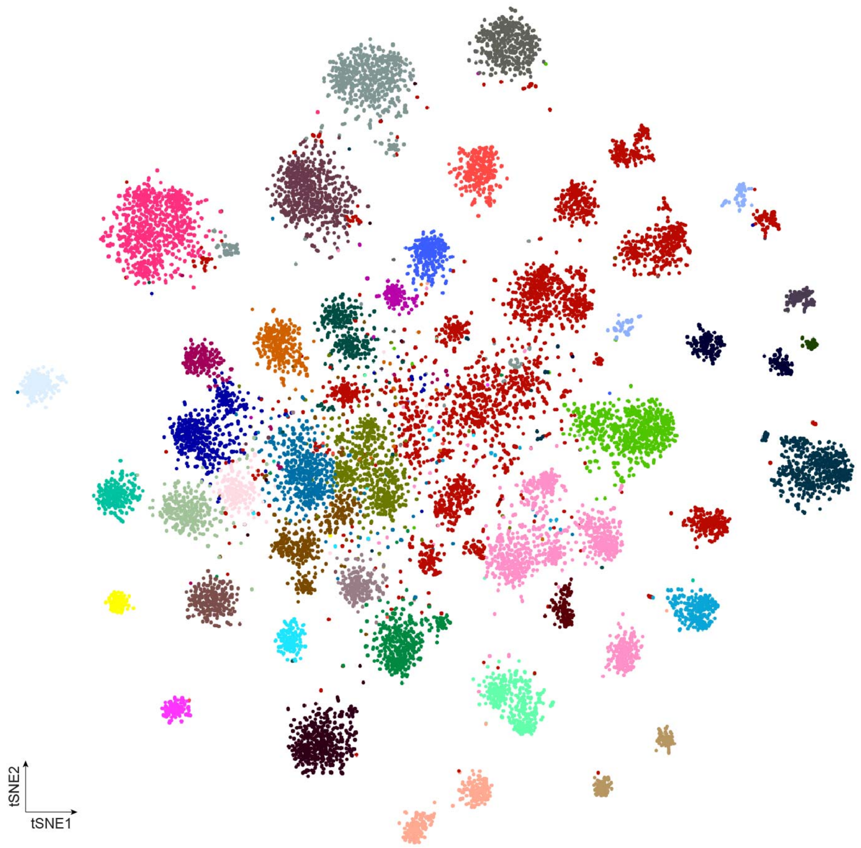


- adult olfactory receptor neuron uncharacterized
- adult olfactory receptor neuron Gr21a, Gr63a
- adult olfactory receptor neuron Gr39a
- adult olfactory receptor neuron Ir41a, Ir76b
- adult olfactory receptor neuron Ir56a
- adult olfactory receptor neuron Ir75d
- adult olfactory receptor neuron Ir92a, Ir84a, Ir31a, Ir76a, Ir76b, Ir8a, Or35a
- adult olfactory receptor neuron Or7a
- adult olfactory receptor neuron Or9a
- adult olfactory receptor neuron Or13a
- adult olfactory receptor neuron Or22a/b
- adult olfactory receptor neuron Or23a
- adult olfactory receptor neuron Or33a, Or56a
- adult olfactory receptor neuron Or42a, Or71a, Or85e
- adult olfactory receptor neuron Or42b
- adult olfactory receptor neuron Or43a/Or2a
- adult olfactory receptor neuron Or43b
- adult olfactory receptor neuron Or46a
- adult olfactory receptor neuron Or47a, Or85f
- adult olfactory receptor neuron Or47b
- adult olfactory receptor neuron Or49b, Or67b, Or69a
- adult olfactory receptor neuron Or59b
- adult olfactory receptor neuron Or65a/b/c
- adult olfactory receptor neuron Or67a
- adult olfactory receptor neuron Or67c
- adult olfactory receptor neuron Or67d
- adult olfactory receptor neuron Or82a
- adult olfactory receptor neuron Or83c
- adult olfactory receptor neuron Or85a
- adult olfactory receptor neuron Or88a
- adult olfactory receptor neuron Or92a
- adult olfactory receptor neuron Or98a
- adult olfactory receptor neuron acid-sensing Ir64a
- adult olfactory receptor neuron acid-sensing, Ir75a/b/c
- sacculus/arista neuron
- sacculus/arista neuron Gr28b
- sacculus/arista neuron Ir21a
- sacculus/arista neuron Ir40a

**fig. S10. tSNE of detailed cell type annotation of the ORNs.**
37 ORN cell types are shown in the tSNE plot and clusters are annotated based on olfactory receptor expression.

fig. S11

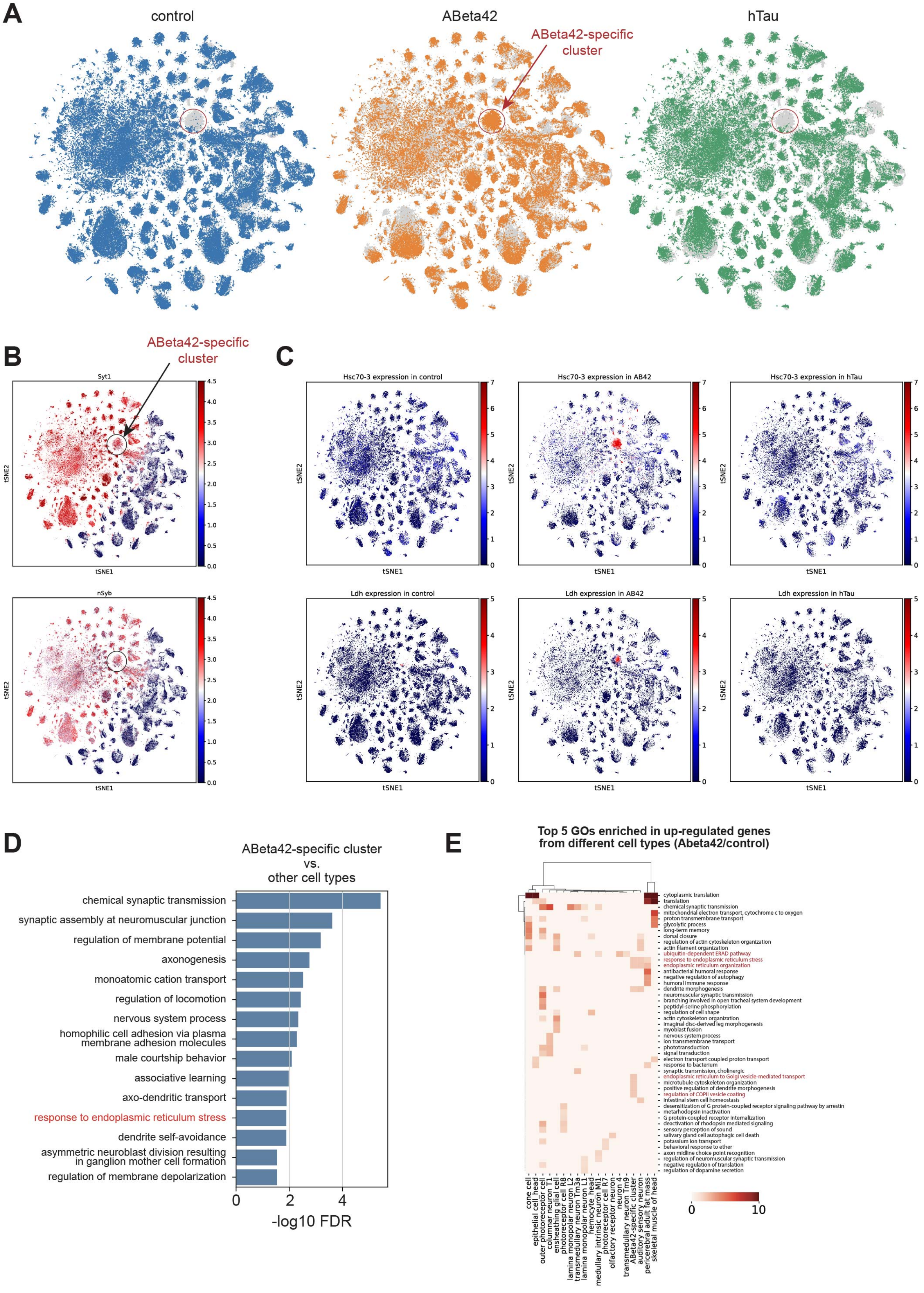
A
control
ABeta42
hTau
ABeta42-specific cluster
B
ABeta42-specific cluster
Syt1
nSyb
tSNE1
tSNE2
C
Hsc70-3 expression in control
Hsc70-3 expression in AB42
Hsc70-3 expression in hTau
Ldh expression in control
Ldh expression in AB42
Ldh expression in hTau
D
ABeta42-specific cluster vs. other cell types
chemical synaptic transmission
synaptic assembly at neuromuscular junction
regulation of membrane potential
axonogenesis
monoatomic cation transport
regulation of locomotion
nervous system process
homophilic cell adhesion via plasma membrane adhesion molecules
male courtship behavior
associative learning
axo-dendritic transport
response to endoplasmic reticulum stress
dendrite self-avoidance
asymmetric neuroblast division resulting in ganglion mother cell formation
regulation of membrane depolarization
0
2
4
-log10 FDR
E
Top 5 GOs enriched in up-regulated genes from different cell types (Abeta42/control)
cytoplasmic translation
translation
chemical synaptic transmission
mitochondrial electron transport, cytochrome c to oxygen
proton transmembrane transport
glycolytic process
long-term memory
dorsal closure
regulation of actin cytoskeleton organization
actin filament organization
ubiquitin-dependent ERAD pathway
response to endoplasmic reticulum stress
endoplasmic reticulum organization
antibacterial humoral response
negative regulation of autophagy
humoral immune response
dendrite morphogenesis
neuromuscular synaptic transmission
branching involved in open tracheal system development
peptidyl-serine phosphorylation
regulation of cell shape
actin cytoskeleton organization
imaginal disc-derived leg morphogenesis
myoblast fusion
nervous system process
ion transmembrane transport
phototransduction
signal transduction
electron transport coupled proton transport
response to bacterium
synaptic transmission, cholinergic
endoplasmic reticulum to Golgi vesicle-mediated transport
microtubule cytoskeleton organization
positive regulation of dendrite morphogenesis
regulation of COPII vesicle coating
intestinal stem cell homeostasis
desensitization of G protein-coupled receptor signaling pathway by arrestin
metarhodopsin inactivation
G protein-coupled receptor internalization
deactivation of rhodopsin mediated signaling
sensory perception of sound
salivary gland cell autophagic cell death
potassium ion transport
behavioral response to ether
axon midline choice point recognition
regulation of neuromuscular synaptic transmission
negative regulation of translation
regulation of dopamine secretion
cone cell
epithelial cell_head
outer photoreceptor cell
columnar neuron T1
ensheathing glial cell
photoreceptor cell R8
lamina monopolar neuron L2
transmedullary neuron Tm3a
lamina monopolar neuron L1
hemocyte_head
medullary intrinsic neuron Mi1
photoreceptor cell R7
olfactory receptor neuron
neuron 4
transmedullary neuron Tm9
ABeta42-specific cluster
auditory sensory neuron
pericerebral adult fat mass
skeletal muscle of head
0
10

**fig. S11. Features of Aβ42-specific cluster.**

(A) tSNE visualizations of the head samples from different genotypes.

(B) Expression of neuronal marker genes (*Syt1* and *nSyb*) in head cells shown by tSNE.

(C) Expression of Aβ42-specific marker genes (*Hsc70-3* and *Ldh*) in head cells shown by tSNE from different genotypes.

(D) Top 15 GOs terms from Aβ42-specific cluster compared to other cell types in head. All top GO terms identified are related to neuronal physiology and function with one exception, response to endoplasmic reticulum (ER) stress.

(E) Top 5 GOs enriched in up-regulated genes from different head cell types in Aβ42 flies compared to control flies.

fig. S12

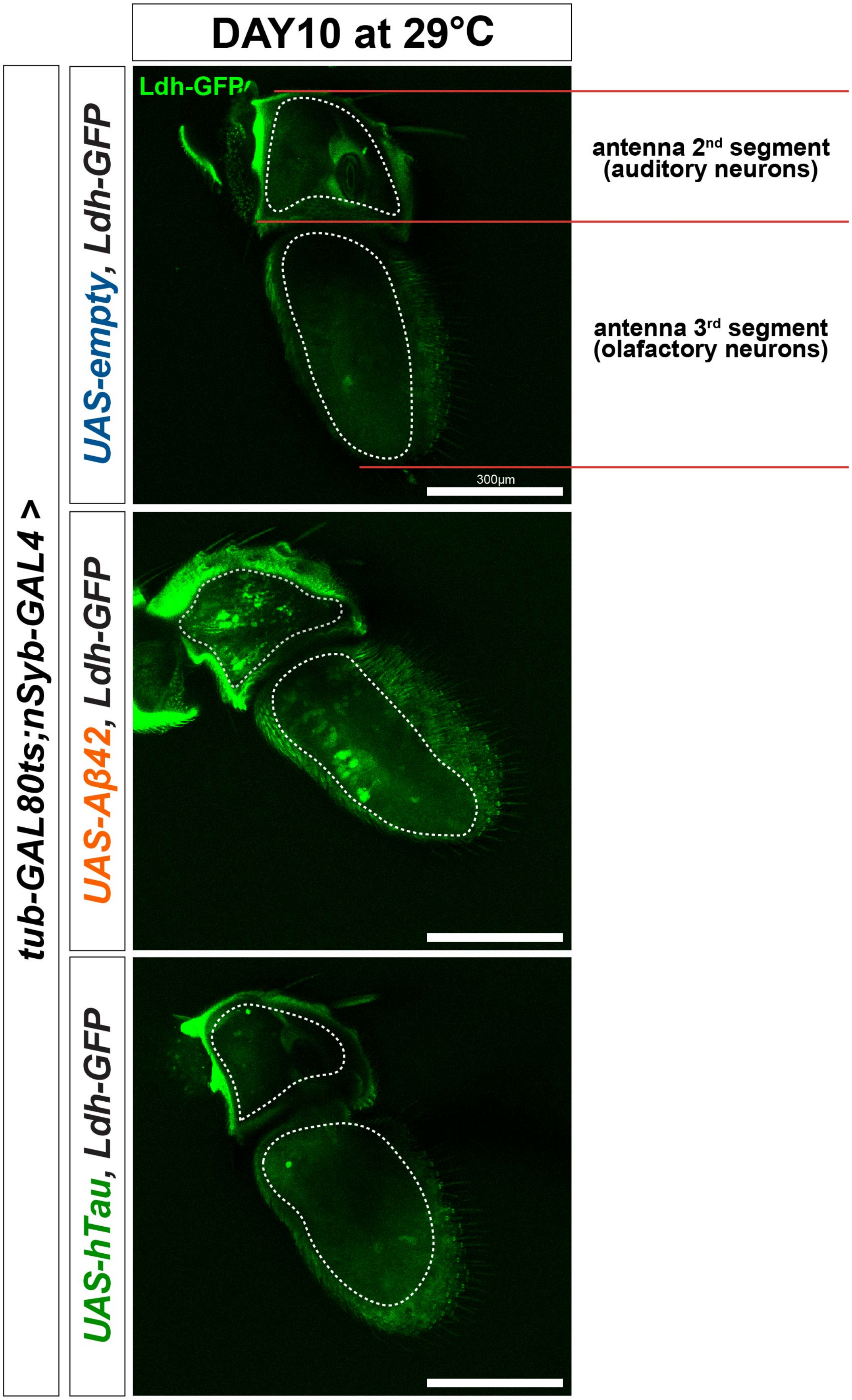

DAY10 at 29°C
tub-GAL80ts;nSyb-GAL4 >
UAS-empty, Ldh-GFP
UAS-Aβ42, Ldh-GFP
UAS-hTau, Ldh-GFP
Ldh-GFP
300µm
antenna 2nd segment
(auditory neurons)
antenna 3rd segment
(olafactory neurons)

**fig. S12. Endogenous Ldh-GFP signals in antennae of 10d flies**

Ldh-GFP (green signals in the dotted lines) is highly expressed in both 2$^{nd}$ segment (auditory neurons) and 3$^{rd}$ segment (olfactory receptor neurons) of the antenna in Aβ42 flies, but not in control or hTau flies. Endogenous GFP signals were immediately imaged after dissection without anti-GFP antibody staining. The representative images were generated by MAX intensity of middle 30 Z-planes. The green signals outside of the dotted lines are laser reflection on the cuticle.

### fig. S13

A

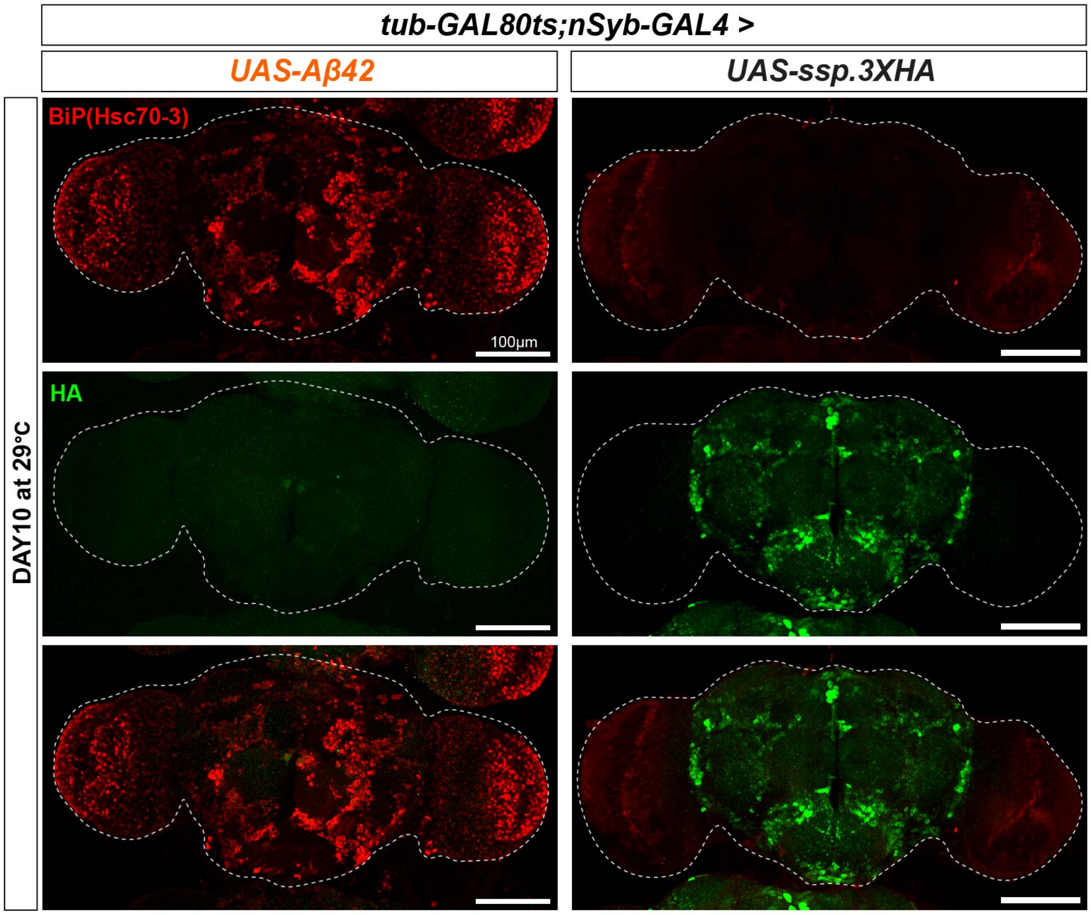
tub-GAL80ts;nSyb-GAL4 >
UAS-Aβ42
UAS-ssp.3XHA
DAY10 at 29°C
BiP(Hsc70-3)
HA
100μm

B

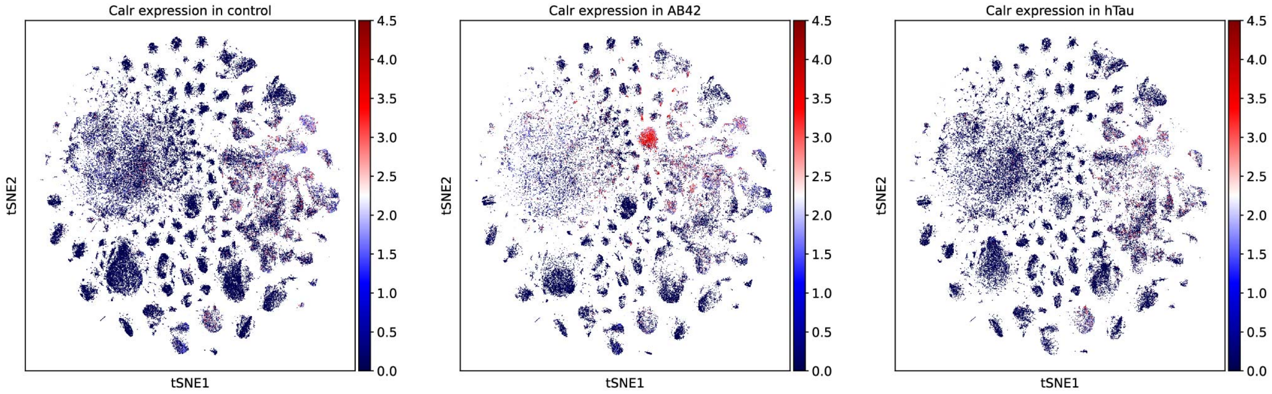
Calr expression in control
Calr expression in AB42
Calr expression in hTau
tSNE1
tSNE2
0.0
0.5
1.0
1.5
2.0
2.5
3.0
3.5
4.0
4.5

**fig. S13. ER stress response in Aβ42 fly brain is not due to the secretion process**

(A) Whole brain staining of 10d Aβ42 fly and *tub-Gal80$^{ts}$;nSyb-GAL4>UAS-ssp.3XHA* fly with anti-Hsc70-3(BiP) (red) and anti-HA (green) antibodies. The expression of ssp.3XHA protein in *ssp.3XHA* control fly brain is confirmed by anti-HA (green) antibodies. The *ssp.3XHA* control fly brain does not show upregulation of Hsc70-3 (BiP) protein, while the Aβ42 fly brain shows significantly increased Hsc70-3 protein levels. The representative images were generated by MAX intensity of whole Z planes.

(B) Expression of *Calr* (apoptosis-related gene) in head cells shown by tSNE from different genotypes.

### fig. S14

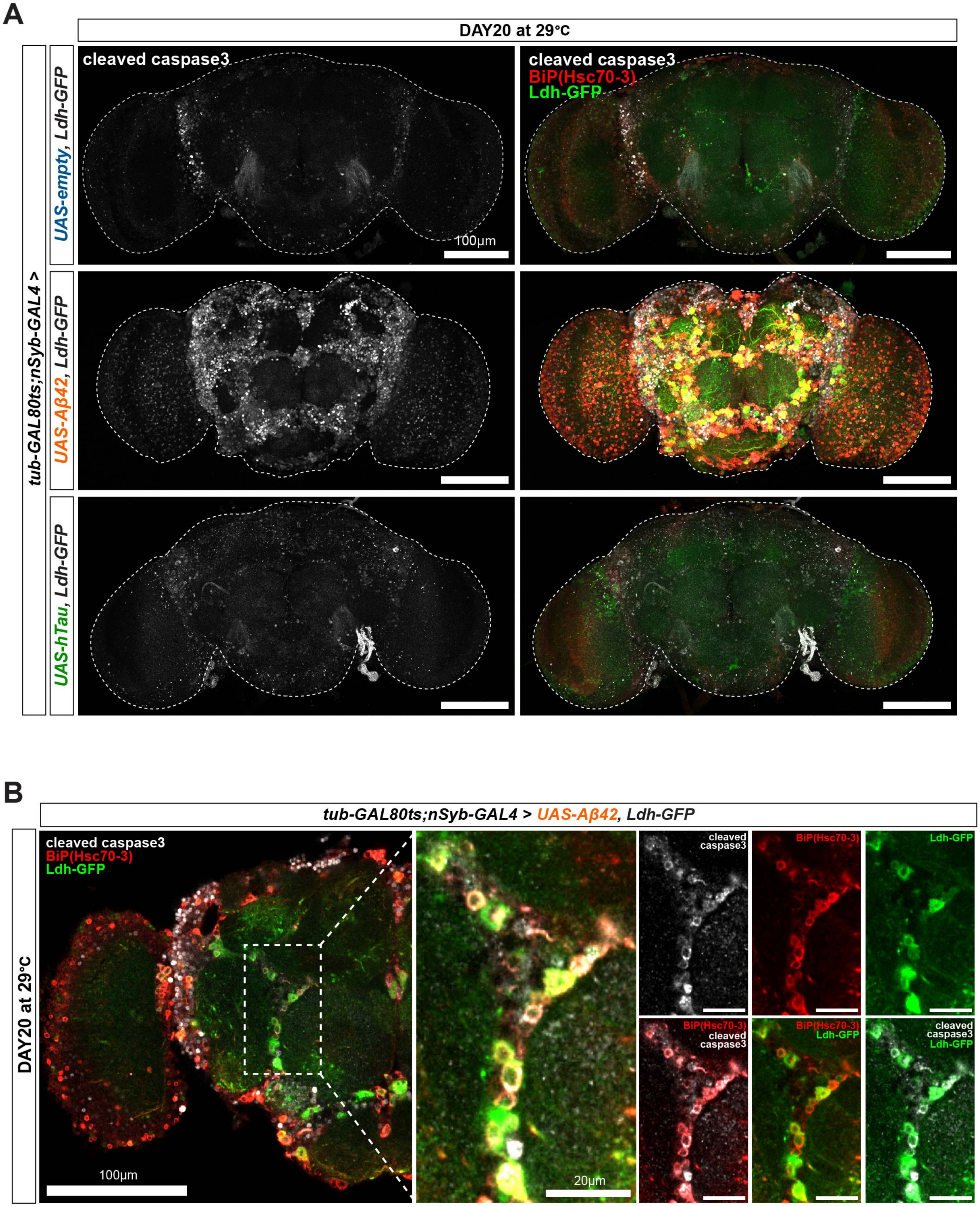
A
DAY20 at 29°C
tub-GAL80ts;nSyb-GAL4 >
UAS-empty, Ldh-GFP
UAS-Aβ42, Ldh-GFP
UAS-hTau, Ldh-GFP
cleaved caspase3
cleaved caspase3
BiP(Hsc70-3)
Ldh-GFP
100μm
B
tub-GAL80ts;nSyb-GAL4 > UAS-Aβ42, Ldh-GFP
DAY20 at 29°C
cleaved caspase3
BiP(Hsc70-3)
Ldh-GFP
100μm
20μm
cleaved caspase3
BiP(Hsc70-3)
Ldh-GFP
BiP(Hsc70-3) cleaved caspase3
BiP(Hsc70-3) Ldh-GFP
cleaved caspase3 Ldh-GFP

**fig. S14. Aβ42 brains show higher apoptosis marker signals compared to control and hTau brains.**

(A) Whole brain images of 10d flies (containing *Ldh-GFP*) showing apoptosis marker (Anti-cleaved-caspase3, gray), ER stress marker (anti-Hsc70-3(BiP), red), and endogenous Ldh-GFP (green). The representative images were generated by MAX intensity of whole Z planes.

(B) Magnified image of Aβ42 brain in fig. S14A. One representative plane of the Z series image was selected. Many BiP+ (and/or Ldh-GFP+) cells in the Aβ42 brain are positive for the cleaved-caspase3 signals.

### fig. S15

A

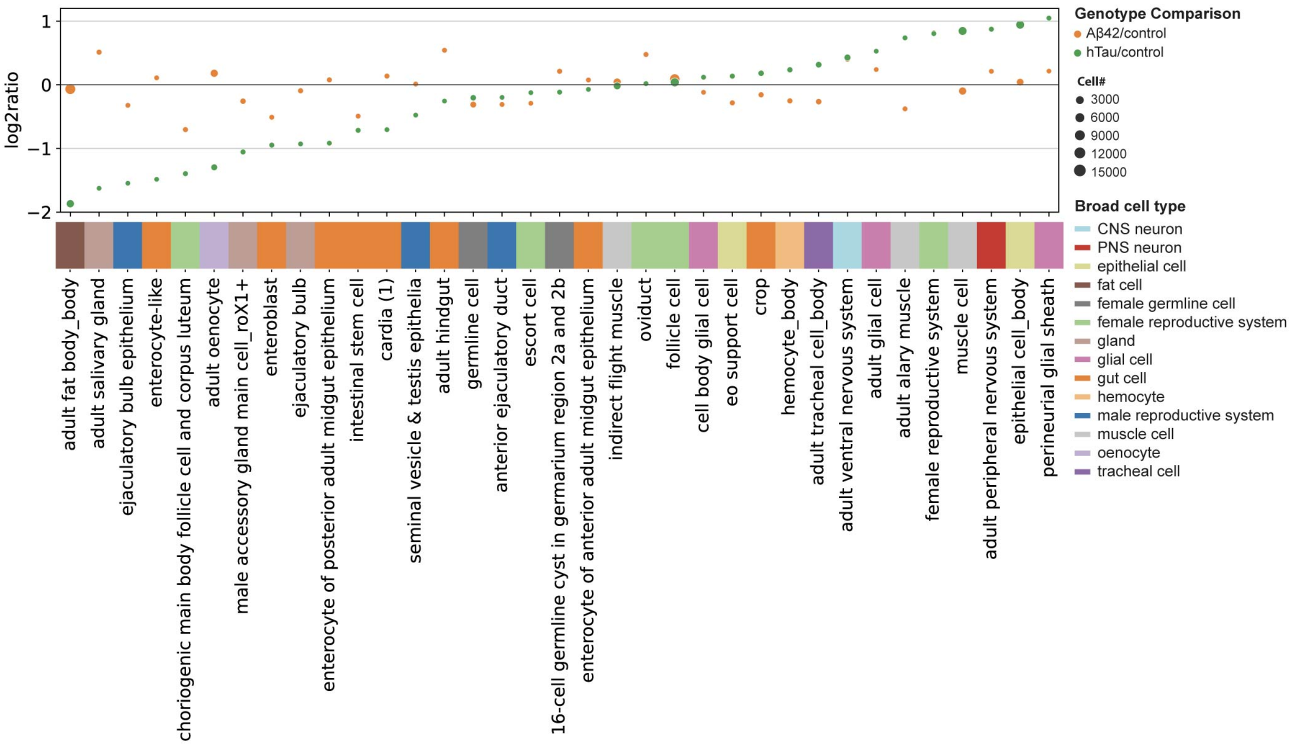
log2ratio
1
0
−1
−2
Genotype Comparison
Aβ42/control
hTau/control
Cell#
3000
6000
9000
12000
15000
Broad cell type
CNS neuron
PNS neuron
epithelial cell
fat cell
female germline cell
female reproductive system
gland
glial cell
gut cell
hemocyte
male reproductive system
muscle cell
oenocyte
tracheal cell
adult fat body_body
adult salivary gland
ejaculatory bulb epithelium
enterocyte-like
choriogenic main body follicle cell and corpus luteum
adult oenocyte
male accessory gland main cell_roX1+
enteroblast
ejaculatory bulb
enterocyte of posterior adult midgut epithelium
intestinal stem cell
cardia (1)
seminal vesicle & testis epithelia
adult hindgut
germline cell
anterior ejaculatory duct
escort cell
16-cell germline cyst in germarium region 2a and 2b
enterocyte of anterior adult midgut epithelium
indirect flight muscle
oviduct
follicle cell
cell body glial cell
eo support cell
crop
hemocyte_body
adult tracheal cell_body
adult ventral nervous system
adult glial cell
adult alary muscle
female reproductive system
muscle cell
adult peripheral nervous system
epithelial cell_body
perineurial glial sheath

B

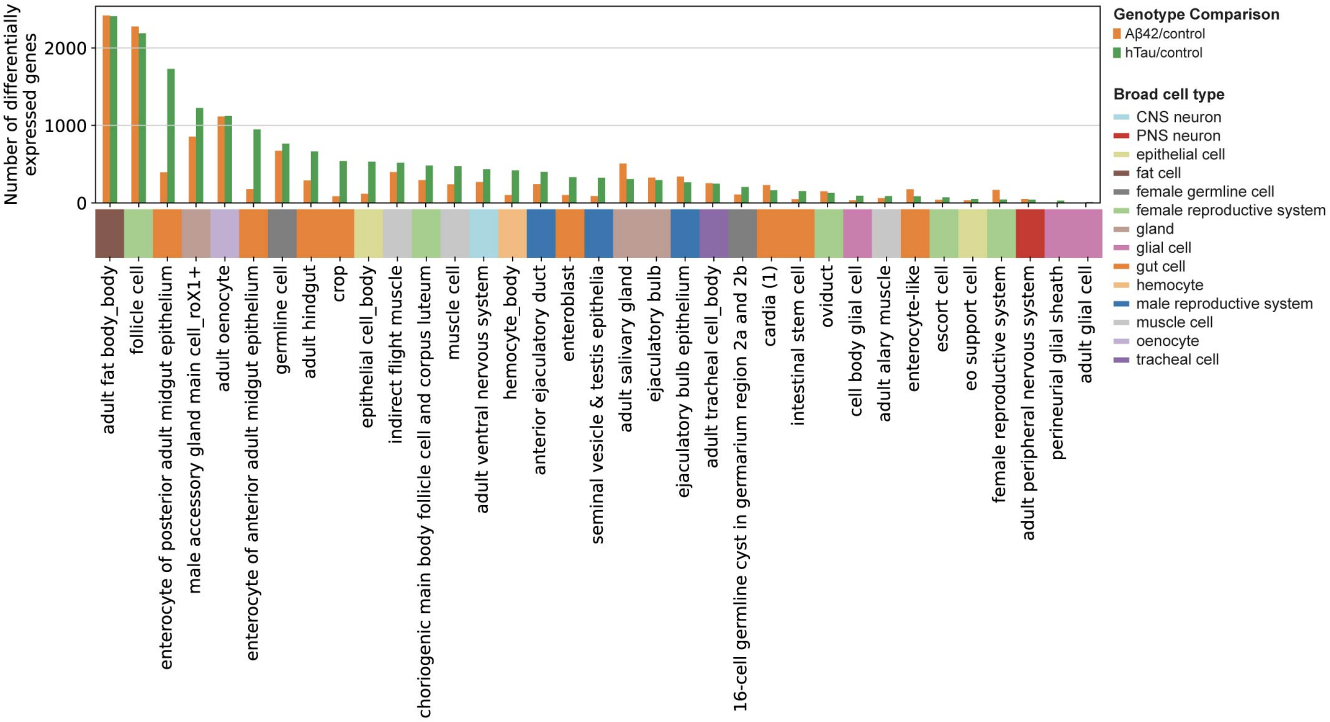
Number of differentially expressed genes
2000
1000
0
Genotype Comparison
Aβ42/control
hTau/control
Broad cell type
CNS neuron
PNS neuron
epithelial cell
fat cell
female germline cell
female reproductive system
gland
glial cell
gut cell
hemocyte
male reproductive system
muscle cell
oenocyte
tracheal cell
adult fat body_body
follicle cell
enterocyte of posterior adult midgut epithelium
male accessory gland main cell_roX1+
adult oenocyte
enterocyte of anterior adult midgut epithelium
germline cell
adult hindgut
crop
epithelial cell_body
indirect flight muscle
choriogenic main body follicle cell and corpus luteum
muscle cell
adult ventral nervous system
hemocyte_body
anterior ejaculatory duct
enteroblast
seminal vesicle & testis epithelia
adult salivary gland
ejaculatory bulb
ejaculatory bulb epithelium
adult tracheal cell_body
16-cell germline cyst in germarium region 2a and 2b
cardia (1)
intestinal stem cell
oviduct
cell body glial cell
adult alary muscle
enterocyte-like
escort cell
eo support cell
female reproductive system
adult peripheral nervous system
perineurial glial sheath
adult glial cell

**fig. S15. Cell composition and DEG analysis in body cell types.**

(A) Cell composition changes across Aβ42 and hTau body cell types against controls.

(B) Number of DEGs identified in Aβ42 and hTau body cell types relative to controls.

### fig. S16

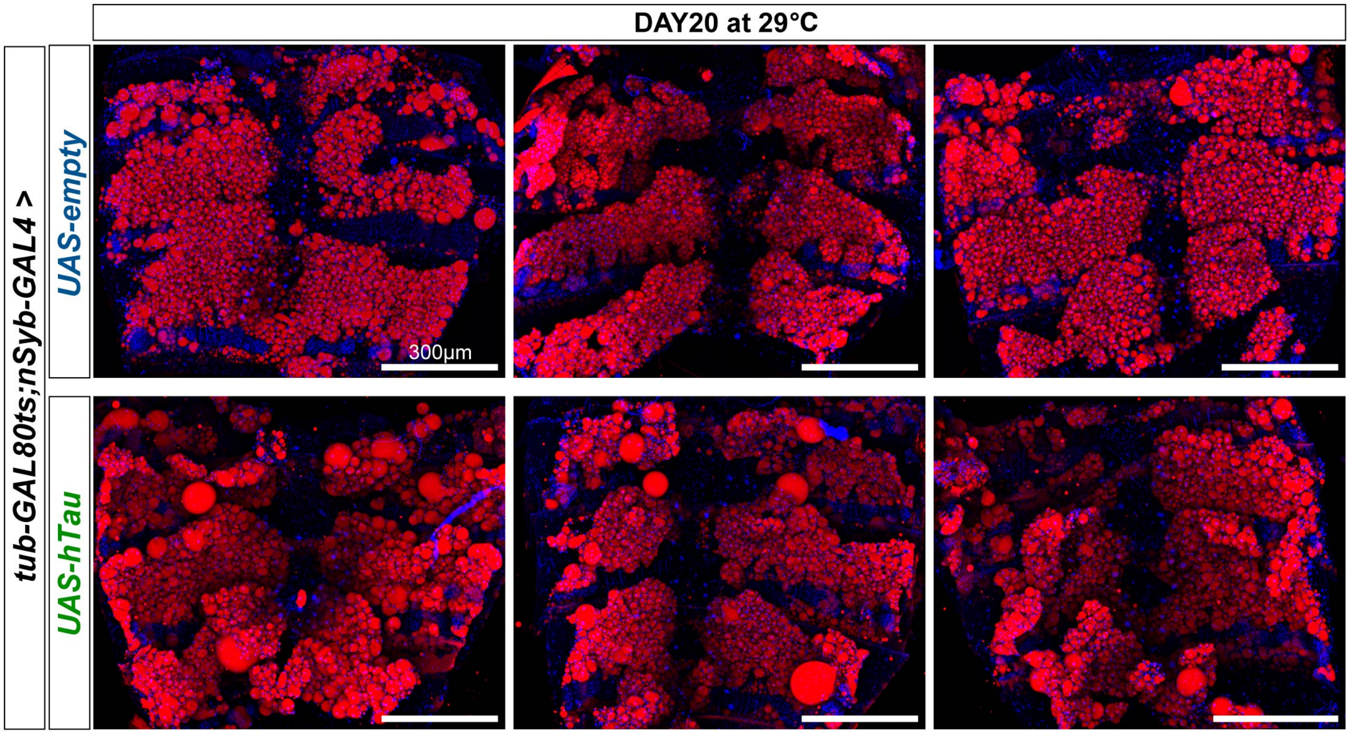

**fig. S16. Examples of fat body LD staining showing entire abdomen filets in 20d control and hTau flies.**

20d hTau flies show significantly enlarged LDs compared to LDs in control flies. LDs (Nile Red, red) and nuclei (DAPI, blue) staining. The fly abdomen filets were imaged using Leica tile scan. The representative images were generated by MAX intensity of whole Z planes.

### fig. S17

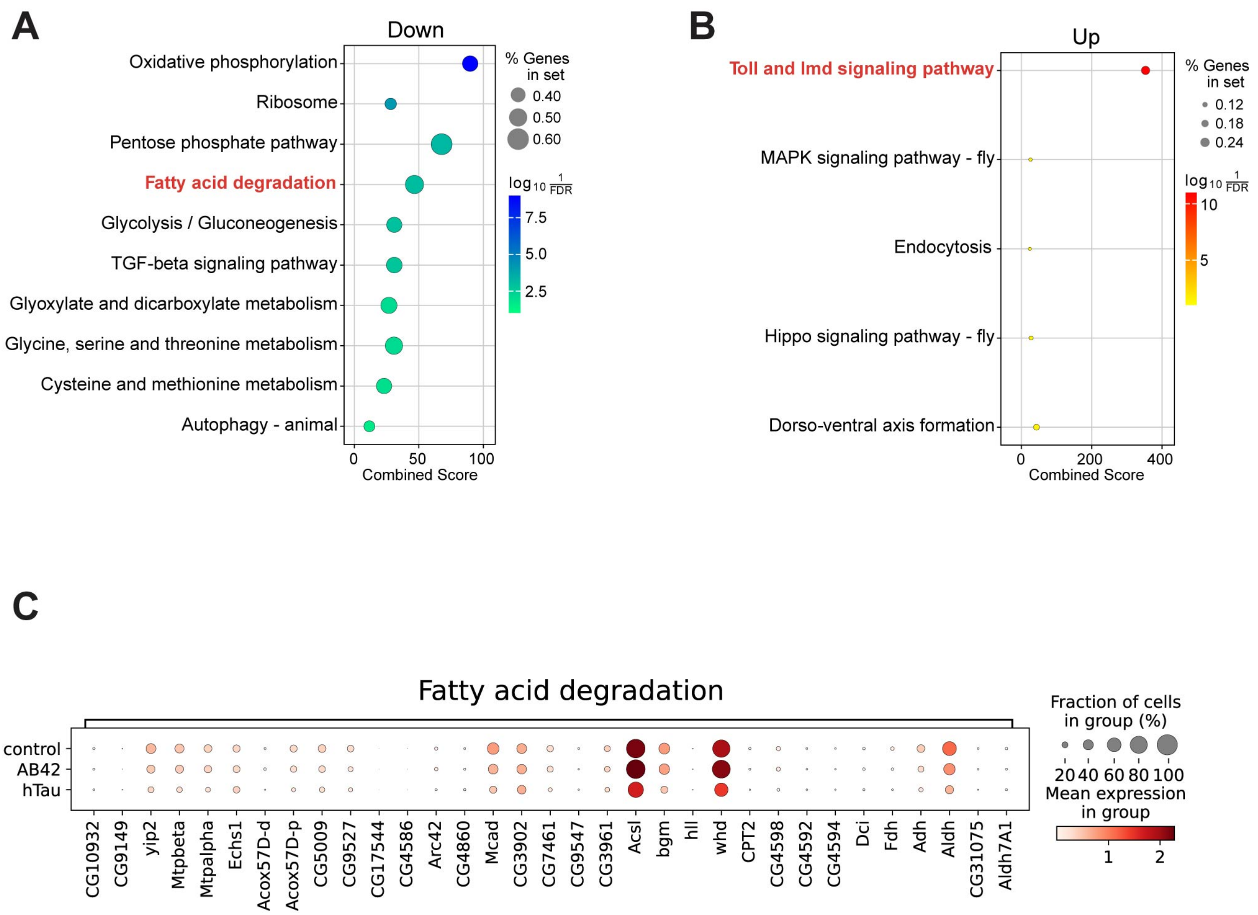

A
Down
Oxidative phosphorylation
Ribosome
Pentose phosphate pathway
Fatty acid degradation
Glycolysis / Gluconeogenesis
TGF-beta signaling pathway
Glyoxylate and dicarboxylate metabolism
Glycine, serine and threonine metabolism
Cysteine and methionine metabolism
Autophagy - animal
0
50
100
Combined Score
% Genes in set
0.40
0.50
0.60
log10 1/FDR
7.5
5.0
2.5
B
Up
Toll and Imd signaling pathway
MAPK signaling pathway - fly
Endocytosis
Hippo signaling pathway - fly
Dorso-ventral axis formation
0
200
400
Combined Score
% Genes in set
0.12
0.18
0.24
log10 1/FDR
10
5
C
Fatty acid degradation
control
AB42
hTau
CG10932
CG9149
yip2
Mtpbeta
Mtpalpha
Echs1
Acox57D-d
Acox57D-p
CG5009
CG9527
CG17544
CG4586
Arc42
CG4860
Mcad
CG3902
CG7461
CG9547
CG3961
Acsl
bgm
hll
whd
CPT2
CG4598
CG4592
CG4594
Dci
Fdh
Adh
Aldh
CG31075
Aldh7A1
Fraction of cells in group (%)
20 40 60 80 100
Mean expression in group
1
2

**fig. S17. Enrichment analysis of KEGG pathways in fat body DEGs**

(A) KEGG pathways enriched among genes down-regulated in hTau flies compared to controls.

(B) KEGG pathways enriched among up-regulated genes in hTau flies.

(C) Expression comparison of genes involved in “Fatty acid degradation” pathway,

fig. S18

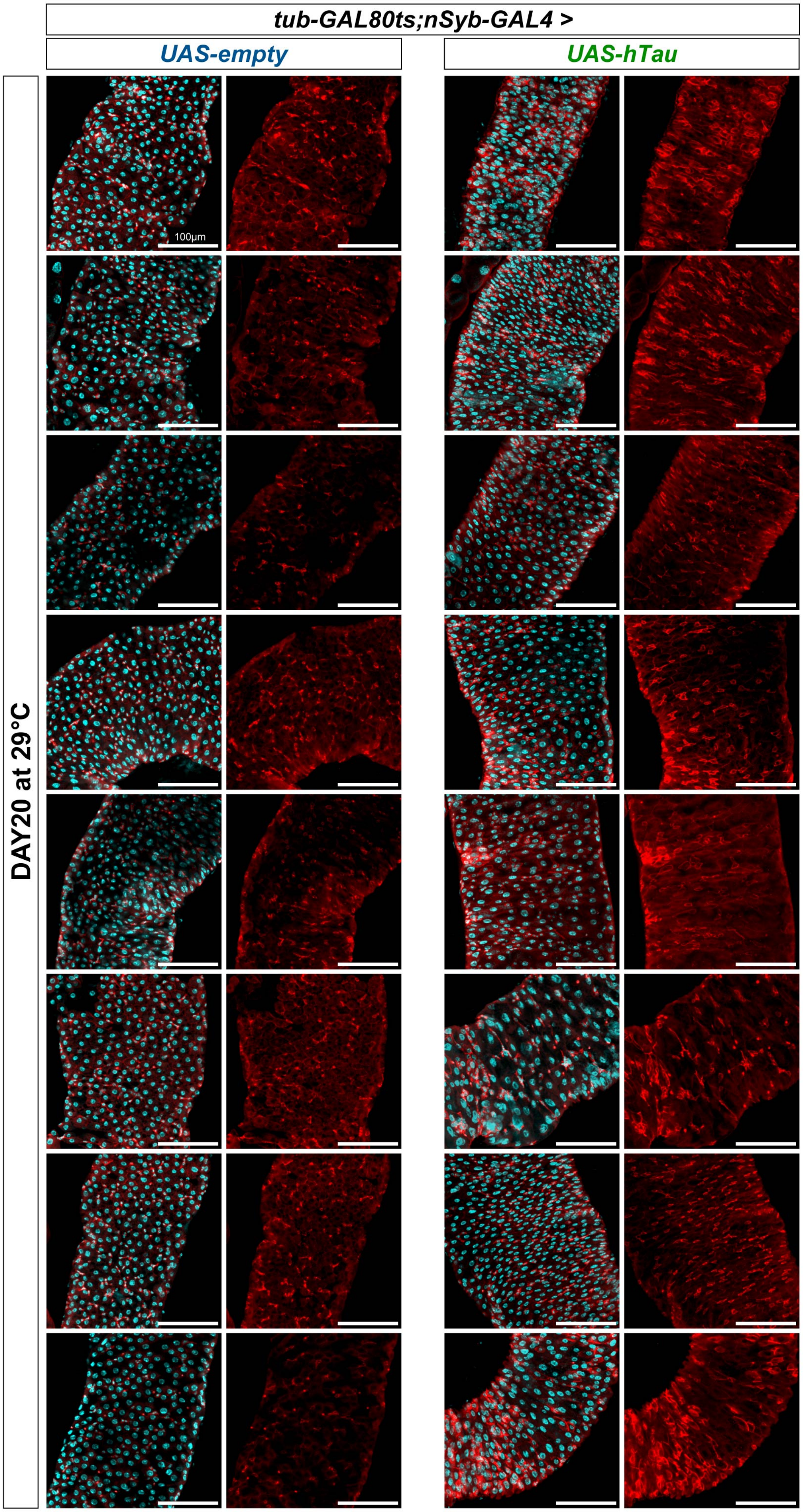

**fig. S18. Examples of posterior midgut in control and hTau flies**

20d hTau fly guts show a decrease of mature polyploidy ECs (large nuclei) and an increase of cells with intermediate-size nuclei with strong armadillo staining. DAPI (cyan), anti-Prospero (red signals in EE nuclei) & anti-armadillo (red signals in cell membrane, stronger in ISCs/EBs than ECs).

### fig. S19

A

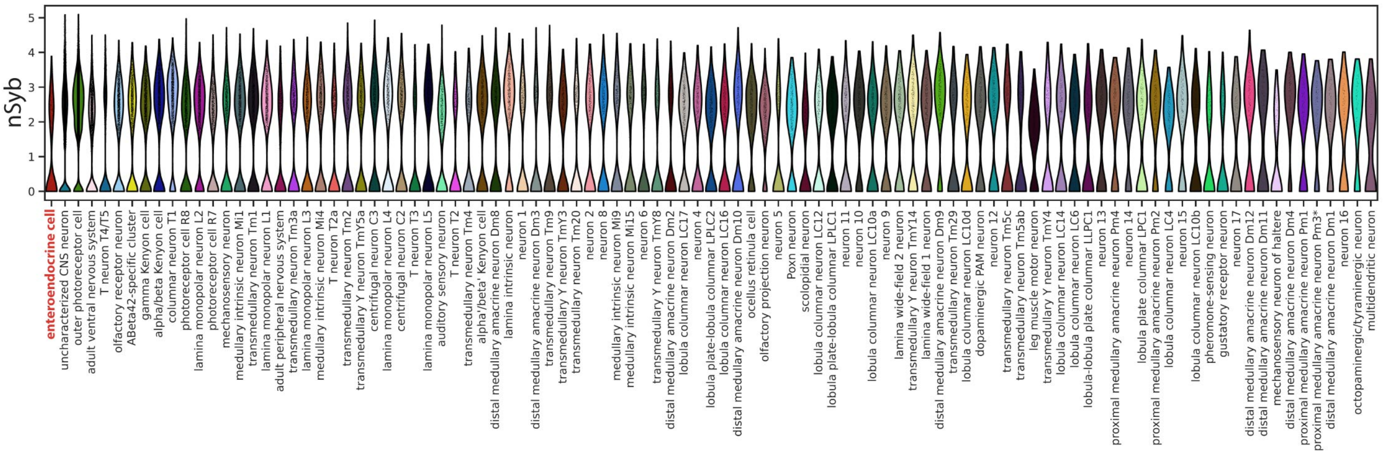
nSyb
0
1
2
3
4
5
enteroendocrine cell
uncharacterized CNS neuron
outer photoreceptor cell
adult ventral nervous system
T neuron T4/T5
olfactory receptor neuron
ABeta42-specific cluster
gamma Kenyon cell
alpha/beta Kenyon cell
columnar neuron T1
photoreceptor cell R8
lamina monopolar neuron L2
photoreceptor cell R7
mechanosensory neuron
medullary intrinsic neuron Mi1
transmedullary neuron Tm1
lamina monopolar neuron L1
adult peripheral nervous system
transmedullary neuron Tm3a
lamina monopolar neuron L3
medullary intrinsic neuron Mi4
T neuron T2a
transmedullary neuron Tm2
transmedullary Y neuron TmY5a
centrifugal neuron C3
lamina monopolar neuron L4
centrifugal neuron C2
T neuron T3
lamina monopolar neuron L5
auditory sensory neuron
T neuron T2
transmedullary neuron Tm4
alpha'/beta' Kenyon cell
distal medullary amacrine neuron Dm8
lamina intrinsic neuron
neuron 1
distal medullary amacrine neuron Dm3
transmedullary neuron Tm9
transmedullary Y neuron TmY3
transmedullary neuron Tm20
neuron 2
neuron 8
medullary intrinsic neuron Mi9
medullary intrinsic neuron Mi15
neuron 6
transmedullary Y neuron TmY8
distal medullary amacrine neuron Dm2
lobula columnar neuron LC17
neuron 4
lobula plate-lobula columnar LPLC2
lobula columnar neuron LC16
distal medullary amacrine neuron Dm10
ocellus retinula cell
olfactory projection neuron
neuron 5
Poxn neuron
scolopidial neuron
lobula columnar neuron LC12
lobula plate-lobula columnar LPLC1
neuron 11
neuron 10
lobula columnar neuron LC10a
neuron 9
lamina wide-field 2 neuron
transmedullary Y neuron TmY14
lamina wide-field 1 neuron
distal medullary amacrine neuron Dm9
transmedullary neuron Tm29
lobula columnar neuron LC10d
dopaminergic PAM neuron
neuron 12
transmedullary neuron Tm5c
transmedullary neuron Tm5ab
leg muscle motor neuron
transmedullary Y neuron TmY4
lobula columnar neuron LC14
lobula columnar neuron LC6
lobula-lobula plate columnar LLPC1
neuron 13
proximal medullary amacrine neuron Pm4
neuron 14
lobula plate columnar LPC1
proximal medullary amacrine neuron Pm2
lobula columnar neuron LC4
neuron 15
lobula columnar neuron LC10b
pheromone-sensing neuron
gustatory receptor neuron
neuron 17
distal medullary amacrine neuron Dm12
distal medullary amacrine neuron Dm11
mechanosensory neuron of haltere
distal medullary amacrine neuron Dm4
proximal medullary amacrine neuron Pm1
proximal medullary amacrine neuron Pm3*
distal medullary amacrine neuron Dm1
neuron 16
octopaminergic/tyraminergic neuron
multidendritic neuron

B

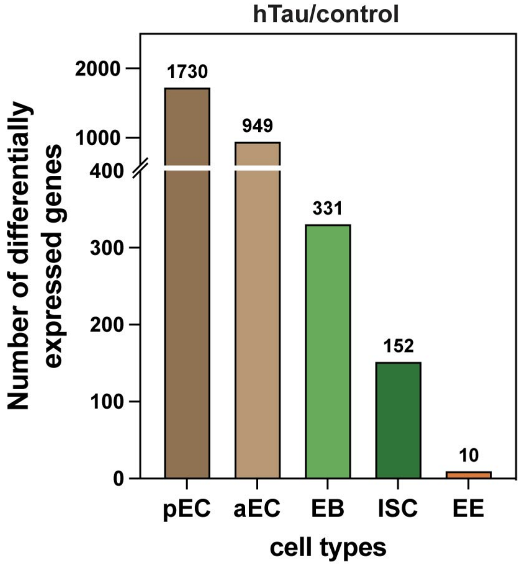
hTau/control
Number of differentially expressed genes
2000
1000
400
300
200
100
0
1730
949
331
152
10
pEC
aEC
EB
ISC
EE
cell types

**fig. S19. Expression of nSyb and DEG numbers in gut cell types.**
(A) The expression of nSyb in enteroendocrine cells is weaker than neuronal cells.
(B) Number of DEGs identified in different gut cell types.

**fig. S20**

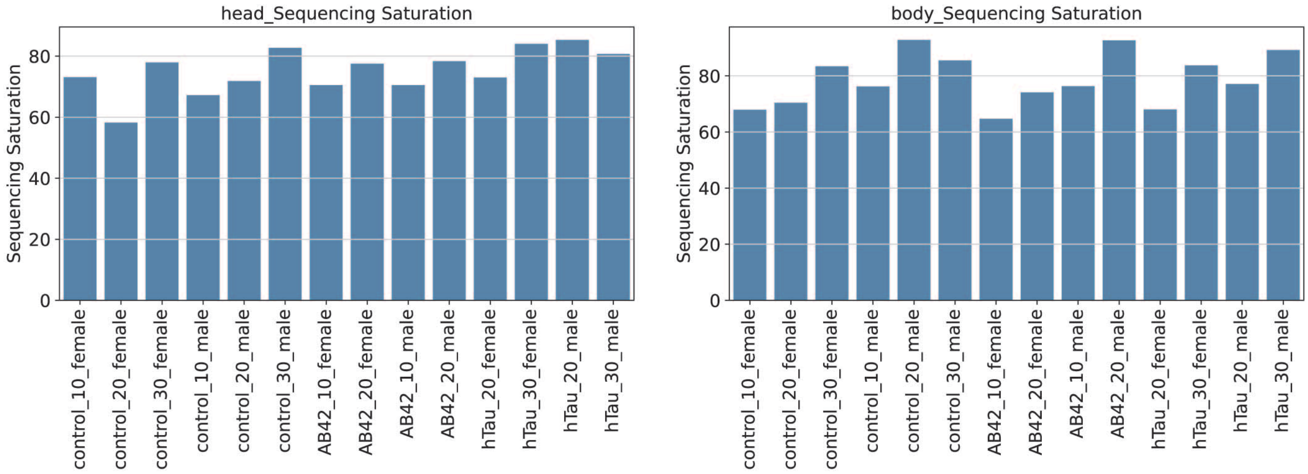

head_Sequencing Saturation
Sequencing Saturation
0
20
40
60
80
control_10_female
control_20_female
control_30_female
control_10_male
control_20_male
control_30_male
AB42_10_female
AB42_20_female
AB42_10_male
AB42_20_male
hTau_20_female
hTau_30_female
hTau_20_male
hTau_30_male
body_Sequencing Saturation
Sequencing Saturation
0
20
40
60
80
control_10_female
control_20_female
control_30_female
control_10_male
control_20_male
control_30_male
AB42_10_female
AB42_20_female
AB42_10_male
AB42_20_male
hTau_20_female
hTau_30_female
hTau_20_male
hTau_30_male

**fig. S20. Sequencing saturation of AD-FCA samples.**
Sequencing saturation levels for each sample, differentiated into head and body categories, highlighting the depth of sequencing achieved across the AD-FCA dataset.

### fig. S21

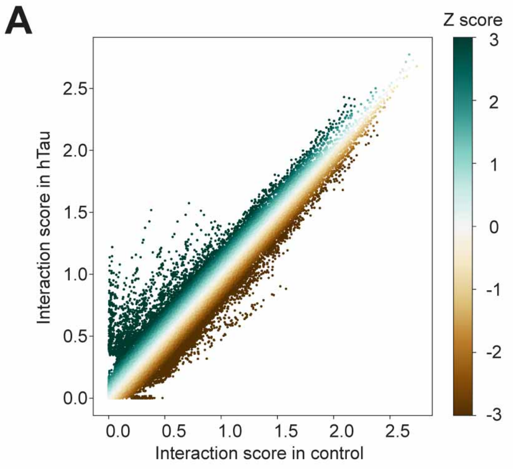
A
Z score
3
2
1
0
-1
-2
-3
2.5
2.0
1.5
1.0
0.5
0.0
Interaction score in hTau
0.0 0.5 1.0 1.5 2.0 2.5
Interaction score in control

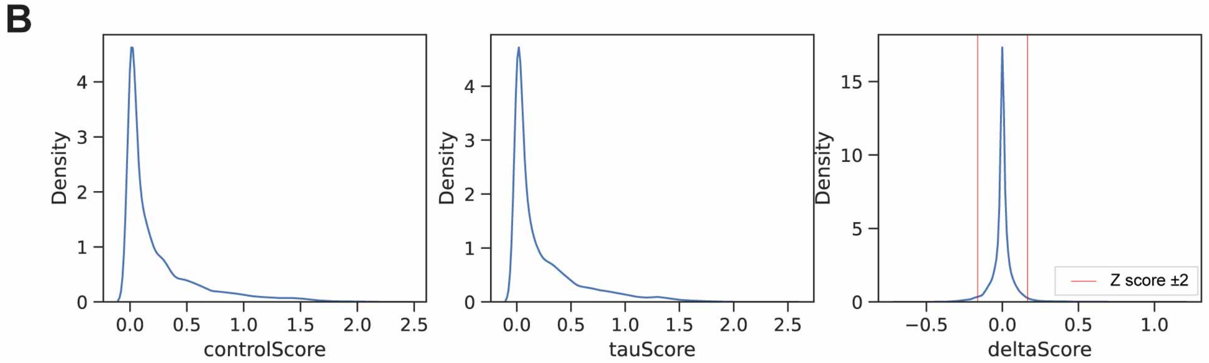
B
Density
4
3
2
1
0
0.0 0.5 1.0 1.5 2.0 2.5
controlScore
tauScore
15
10
5
−0.5 0.0 0.5 1.0
deltaScore
Z score ±2

**fig. S21. CCC scores from control and hTau flies**

(A) Comparison of Z score distributions between hTau and control flies.

(B) Distributions for control CCC scores, hTau CCC scores, and Δscores. Red lines in Δscore represent significant deviation thresholds (±2 Z score).
